# IFNγ-CXCL9-CXCR3 axis links T cell recruitment to skeletal muscle remodelling in human physical frailty

**DOI:** 10.64898/2026.10.27.762096

**Authors:** Vanda WT Ho, Chenshi Lin, Arnab Ray, Hui San Chin, Wei Siong Ong, Heng Boon Low, Dorcas MH Chia, Bhuvaneshwari Shunmuganathan, Chung Hwee Thiam, Paul Hutchinson, Frederick HX Koh, MAGNET group, Sylvie Alonso, Shifeng Xue, Arne N. Akbar, Jinmiao Chen, Catherine WM Ong, Paul A MacAry

**Affiliations:** Department of Medicine, National University of Singapore (NUS), Singapore; Immunology Translational Research Program, Department of Medicine, NUS; Department of Medicine, National University Hospital (NUH), Singapore; Department of Biological Sciences, NUS; Research Office, Sengkang General Hospital, Singapore; Immunology programme, Life Sciences Institute, NUS; Department of Microbiology and Immunology, NUS; Life Sciences Institute, NUS; SingHealth Duke-NUS Muscle Health Program, SingHealth, Singapore; Department of Colorectal Surgery, Sengkang Hospital, Singapore; Infectious Diseases Translational Research Programme, Department of Microbiology and Immunology, Department of Medicine, NUS; Communicable Diseases Agency, Singapore; Division of Infection and Immunity, Faculty of Medical Sciences, University College London, United Kingdom; Centre for Biomedical Data Science and Program in Cancer and Stem Cell Biology, Duke-NUS Medical School, Singapore, Singapore; Bioinformatics Institute (BII), Agency for Science, Technology and Research (A*STAR), Singapore, Singapore; Institute for Health Innovation and Technology, NUS

## Abstract

**Introduction:** Physical frailty is increasingly prevalent in our ageing society, and results in higher morbidity and healthcare costs. Yet, targeted therapeutics are lacking because underlying mechanisms are poorly defined. Human cohort studies have linked inflammation with frailty, but do not define the mechanisms by which the ageing immune system affects skeletal muscle and physical function. Defining this immune-muscle interaction is therefore crucial for identifying therapeutics for human physical frailty.

**Methods:** We first studied two independent human ageing cohorts (discovery n=22; validation n=33). Physical frailty was quantified with age-adjusted timed-up-and-go (TUG) residuals. Peripheral blood was analysed for plasma OLINK proteomics, high-dimensional spectral immunophenotyping and in a subset, single-cell RNA-sequencing of PBMCs with paired TCR profiling. We then examined an independent human skeletal-muscle cohort (n=340) by bulk RNA-sequencing to derive our proposed immune-muscle axis. Finally, we interrogated individual steps of this axis in primary human *in vitro* models using IFNγ-conditioned myotube-fibroblast co-cultures, PBMC transwell migration assays with CXCL9 neutralisation (anti-CXCL9) and receptor CXCR3 antagonism (AMG487). Macrophage co-cultured with CD8 were sequentially co-cultured with myotube/fibroblast co-culture, supernatant was then analysed for fibrosis mediators. We used fixed-effect meta-analysis, multivariable models and sequential mediation for analyses.

**Results:** Physical frailty, by single-cell analyses, was associated with a compartmentalised immune-remodelling state involving inflammatory, tissue-remodelling and trafficking programmes in myeloid and T-cells. This finding was validated with plasma proteomics of the same human cohort, where physical frailty was significantly positively associated with an interferon-gamma (IFNγ)/CXCR3 (IL-18R1, IFNγ, CXCL9) module, within which plasma CXCL9 was the strongest correlate. CD8-TEMRA and γδ T-cells were most responsive to CXCL9 in physical frailty. This expanded CD8-TEMRA compartment was polyclonal, consistent with bystander IFNγ production rather than a shared antigen-specific response. In human muscle, CXCL9 was significantly positively associated with T-cell module, and through this, fibrosis. Conversely, fibrosis was associated with macrophages. Sequential mediation was consistent with a CXCL9-T-cell-macrophage-fibrosis pathway. Finally, in *in-vitro* human primary cell experiments, IFNγ-conditioned myotube-fibroblast cultures produced CXCL9 and attracted CD8-TEMRA subsets. Anti-CXCL9 and CXCR3 antagonism significantly reduced this migration, indicating that recruitment of CD8-TEMRA cells was dependent on the CXCL9-CXCR3 axis. Incubating with activated CD8 cells shifted macrophages towards mixed CD163+CD86+ and pro-inflammatory CD163-CD86+ phenotypes. These CD8-co-cultured macrophages in myotube/fibroblast co-culture were associated with increased fibrosis markers transforming growth factor beta-1 and plasminogen activator inhibitor-1.

**Conclusion:** We identified a CD8-TEMRA T-cell migration to human muscle dependent on IFNγ-CXCL9-CXCR3 axis that is associated with human TUG-defined physical frailty which presents potential therapeutic avenues.

**Graphical abstract:** By integrating systemic immune profiling (discovery n = 22, validation n = 33), an independent human muscle cohort (n = 340) and primary human *in-vitro* model, we found that an IFNγ-CXCL9-CXCR3 axis was associated with timed-up-and-go–defined physical frailty. In blood, an IFNγ-CXCR3 programme, dependent on CXCL9, was enriched in clonally expanded but polyclonal CD8 TEMRA cells, consistent with antigen-independent IFNγ production. In muscle, CXCL9 was linked to fibrotic remodelling through sequential T-cell and macrophage signatures. *In vitro*, IFNγ-conditioned myotube/fibroblast co-culture produced CXCL9 and attracted CD8 TEMRA subsets. This migration was reduced by anti-CXCL9 and receptor CXCR3 antagonist AMG487. Incubating with activated CD8 cells induced mixed CD163+CD86+ and pro-inflammatory CD163-CD86+ macrophage phenotypes. These CD8-co-cultured macrophages in myotube/fibroblast co-culture were associated with increased fibrosis markers TGF-β1 and PAI-1, implicating a chronic, antigen-independent immune-trafficking circuit and a druggable CXCR3 node in human physical frailty.

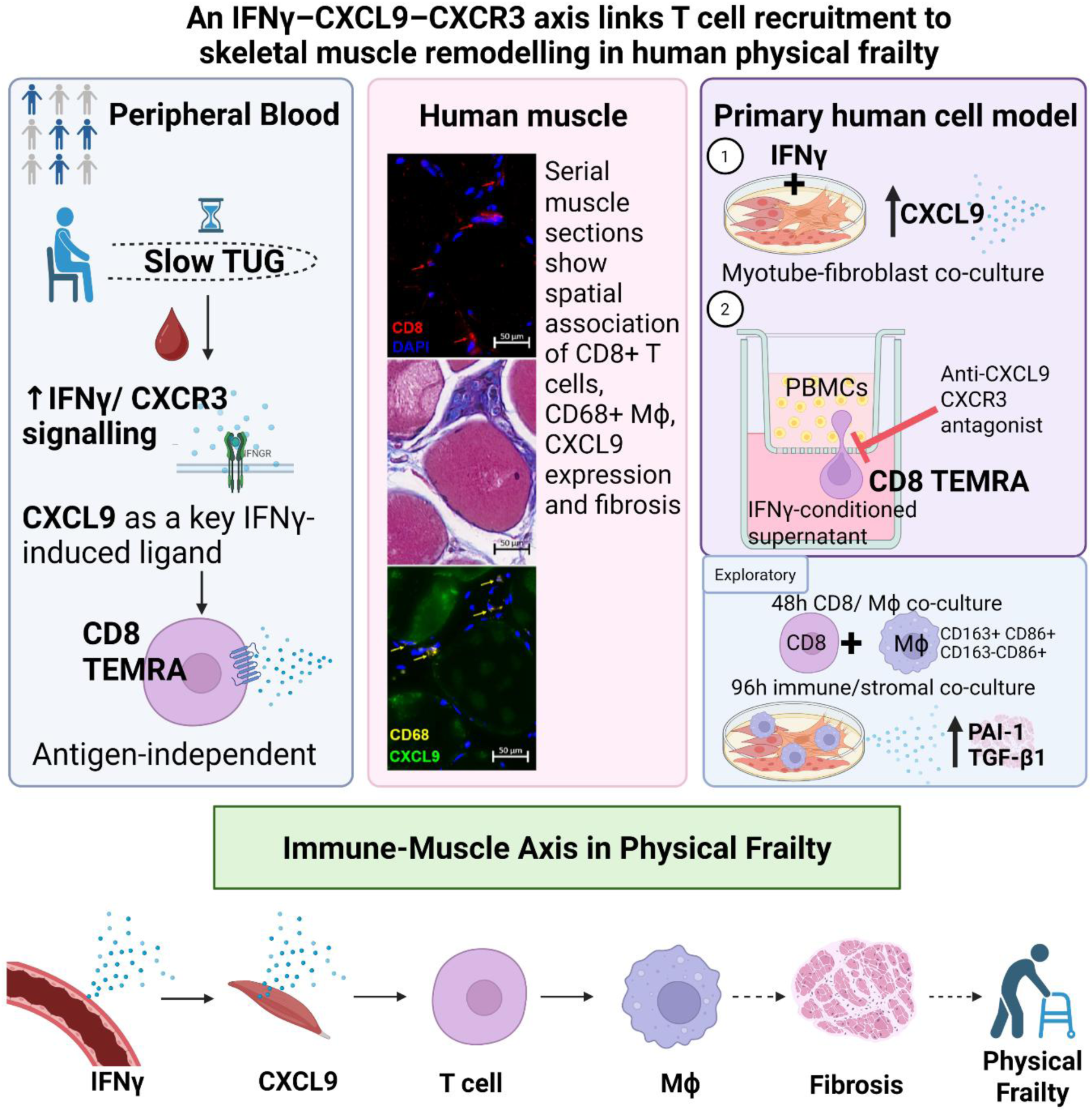

## Introduction

Physical frailty continues to increase in prevalence with rapidly aging societies^1^. Physical frailty increases vulnerability to disability, institutionalisation, and death^2^ and is associated with higher healthcare costs and care needs^3^ ^4^. Exercise, nutritional support, and medication optimisation remain the mainstays of frailty management, but benefits are heterogeneous and incremental^5^. This current state largely reflects a fundamental gap in understanding the mechanisms of why individuals of similar chronological age differ markedly in mobility and resilience. Determining these mechanisms is a prerequisite for precise intervention to physical frailty, and a growing body of evidence implicates the immune system as a key contributor^6–13^.

From human cohort studies, chronic low-grade inflammatory circulatory markers, including interleukin-6 (IL-6), C-reactive protein, and tumour necrosis factor (TNF), are upregulated in frail individuals^8–12^, and CMV-associated immunosenescence has likewise been linked to frailty^7^ ^14^ ^15^. In particular, CXCL9, as a blood-based marker of inflammaging had been associated with reduced muscle strength^16^. However, these signals are non-specific, inconsistently replicated and remain associative.

Separately in skeletal muscle, the end organ of physical frailty, a pro-inflammatory signal has been observed^17–20^. Single-cell and single-nucleus muscle atlases show that with age, human skeletal muscle undergoes coordinated immune and stromal remodelling, including altered macrophage states, chemokine signalling, and profibrotic fibro-adipogenic progenitor (FAP) programmes^17–19^. A more pronounced version of this immune-stromal niche is evident in inflammatory myopathies such as inclusion body myositis, where cytotoxic T cells, inflammatory macrophages, and profibrotic stromal populations coexist within diseased muscle^20^. In this pro-inflammatory setting, CXCL9 and CXCL10 are upregulated in affected muscle, and IFNγ has been shown to induce these chemokines^21^. Functionally, JAK inhibition in rheumatoid arthritis^22^ has been associated with improved muscle function, possibly via IFN–JAK signalling. However, where physical frailty lies on the spectrum between the expected low-grade remodelling in ageing and the antigen-specific autoimmunity of inflammatory myopathy remains unknown.

Clinical cohorts have shown elevated systemic inflammatory mediators in frailty^6–13^, while studies of aged and inflamed skeletal muscle have documented local immune-stromal remodelling^17–20^. Yet, the mechanism of how the blood and muscle compartments are connected in physical frailty remains unknown.

Here, we determined whether a defined, antigen-independent immune circuit links systemic immune ageing to skeletal muscle in human physical frailty. We first studied blood samples from two independent human ageing cohorts with physical frailty characterised by timed-up-and-go (TUG). We then examined muscle bulk transcriptomic data from a third independent human cohort with skeletal muscle biopsies to derive an immune-muscle axis. We interrogated components of this axis in primary human *in-vitro* models using IFNγ-conditioned myotube-fibroblast co-cultures, PBMC transwell migration assays and macrophage-stromal co-cultures. An IFNγ-CXCL9-CXCR3 axis was found to link clonally expanded but polyclonal CD8 TEMRA cells in blood to CXCL9-driven T-cell recruitment and macrophage–-stromal matrix remodelling in muscle, implicating a chronic, antigen-independent immune-trafficking circuit and a druggable CXCR3 node to mitigate human physical frailty.

## Results

Two human cohorts were recruited and were community-dwelling during the follow-up period of three years (n=55; Figure 1A). Comorbidity burden was low (median Charlson comorbidity index, CCI=0), and 78.20% participants (n=43) were CMV seropositive (Table S1).

**Figure 1.**
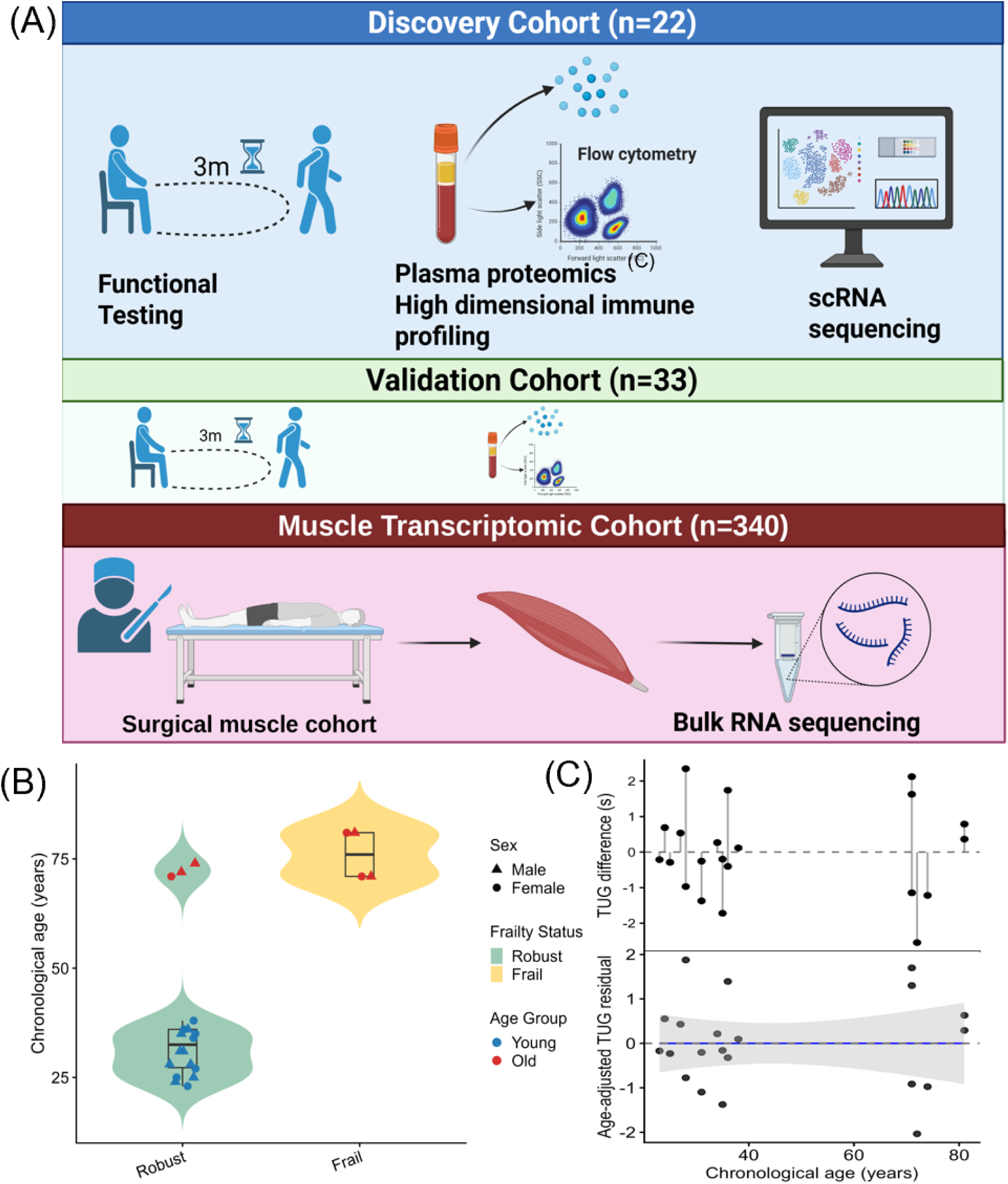
Study design and derivation of age-adjusted TUG residuals. (A) The discovery human cohort (n=22) underwent functional testing using timed-up-and-go (TUG), plasma proteomics, high-dimensional immunophenotyping and single-cell RNA sequencing of PBMCs collected at baseline. A separate human cohort (n=33) underwent identical readouts. An independent human cohort undergoing surgery (n=340) underwent abdominal muscle biopsy with muscle bulk RNA sequencing. Created with BioRender. (B) Chronological age by baseline TUG group in the discovery cohort. (C) Derivation of age-adjusted TUG residuals in the discovery cohort. **Top panel:** Each dot represents one participant, the vertical line shows the difference between their observed TUG and the TUG predicted for their chronological age (the age-expected value from a linear model of TUG on age); points above the dashed zero line walked slower than expected for their age. **Bottom panel:** the same deviations expressed as age-adjusted TUG residuals, plotted against chronological age. The fitted line and shaded 95% confidence band are flat, confirming the residuals carry no remaining age trend.

### Physical frailty does not coincide with chronological age

To measure frailty, we used TUG as the primary measure as it captures integrated motor performance including balance, lower-limb strength, gait and coordination, and has a locally validated cut-off^23^. Maximum handgrip strength and sit-to-stand time were pre-specified sensitivity analyses. Four of 22 discovery-cohort participants (18.2%) met the prespecified physical frailty criteria of slow TUG score, taking more than 9.45 seconds to complete the test^23^. Frail participants were older (Figure 1B), and baseline 3-metre TUG time increased with age (β=0.07s per year, 95% CI: 0.04-0.1, p<0.01), whereas sex was not associated (p=0.98; Figure S1A). Physical frailty was stable over 3 years, as all participants retained their status at recruitment and none crossed the robust or frail threshold (Figure S1B).

To distinguish physical frailty from the expected effects of chronological age, we used an age-residualisation approach. Within each cohort, we fitted a linear model predicting TUG score expected for a person of a given age. For each participant, we then took the residual, which was the difference between their actual TUG score and the score predicted from their age alone. A positive residual meant a person was slower than expected for their age and therefore frailer, capturing the portion of physical frailty that is not explained by chronological age. This age-adjusted residual was used as a continuous measure in all subsequent analyses. Sex was not included as it was not associated with TUG. Age accounted for majority, but not all, of the variation in TUG. In the discovery cohort, age explained 59.60% of the variance (R²=0.60, adjusted R²=0.58; p<0.001), leaving approximately 40% unexplained as the frailty residual we set out to study (Figure 1C). The validation cohort had similar results (Figure S2).

### Physical frailty was associated with altered gene expression in monocytes, CD8 TEMRA and γδ T cells

To determine differences between frail and robust participants in an unbiased manner, we profiled baseline PBMCs from an exploratory single-cell subset of the discovery cohort (Table S2). Major immune-lineage proportions did not differ in frailty (Figure 2A *inset*; Figures S3A and S3B). However, exploratory cell-level differential-expression analysis identified frailty changes within specific immune cell subsets. With the robust group as the reference, downregulated genes were predominantly in monocytes (Figure 2C), whereas fewer genes were upregulated and mainly in T cells, particularly CD8 TEMRA, CD4 naïve, and γδ T cells (Figure 2A). These counts indicated the number of differentially-expressed genes within each cell type and not corrected for donor-level differences. Hence, it is exploratory and primarily used to nominate cell types for the cohort-level analyses that follow. Gene-level differences between frail and robust groups included inflammatory and tissue-remodelling programmes in myeloid cells and stress-adaptation, trafficking, and cytotoxic programmes in lymphocytes (Figure S3C). Gene-ontology analysis highlighted reduced differentiation and regulatory pathways, particularly in CD8 TEMRA cells, and increased antigen-processing, MHC-I, and stress-response pathways in CD4 central-memory and γδ T cells in frailty (Figures 2B and 2D).

**Figure 2.**
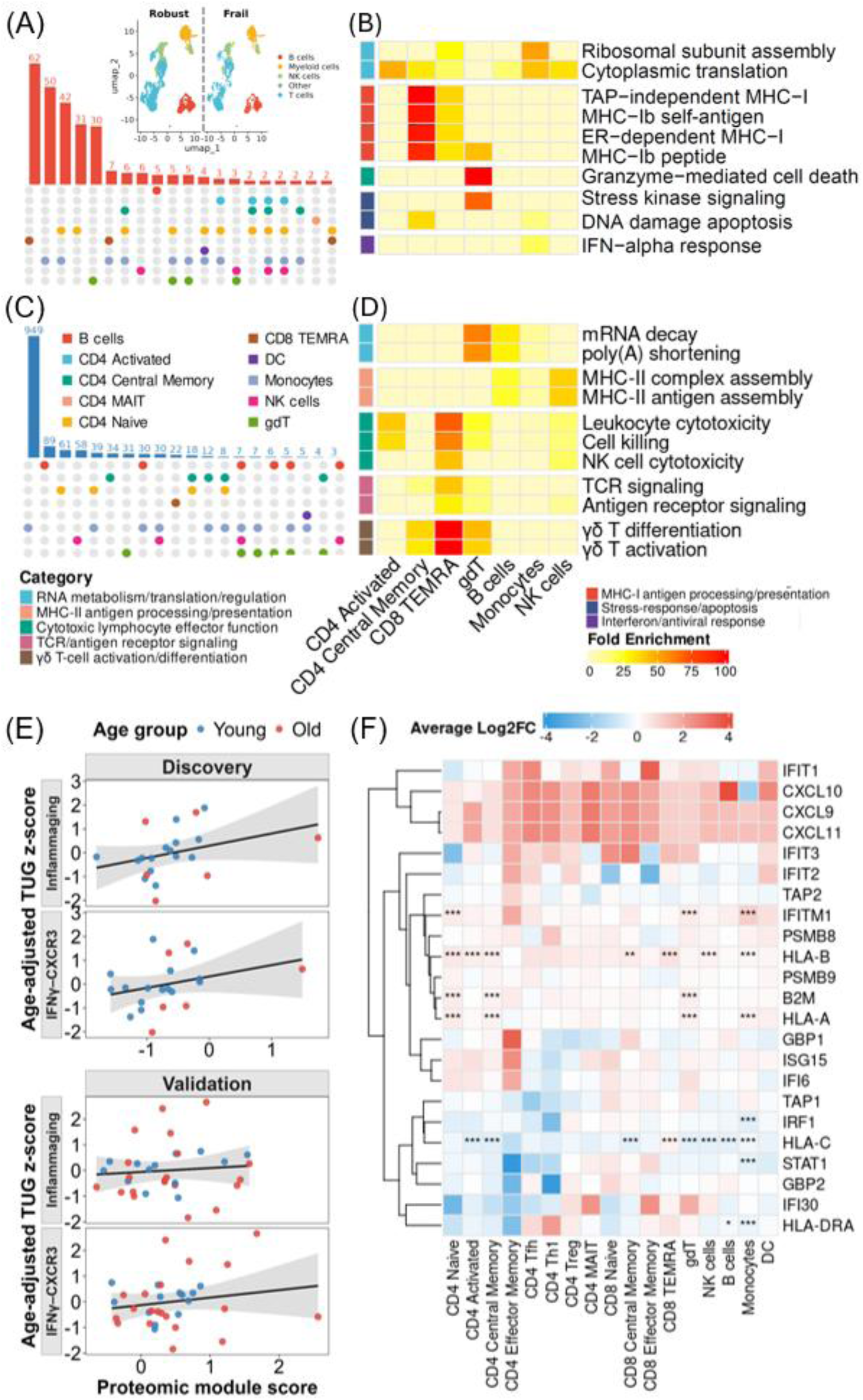
Physical frailty was associated with subset-specific immune transcriptional remodelling and an upregulated IFNγ–CXCR3 plasma programme. (A) UpSet plot showing overlaps of upregulated genes which were differentially expressed in frail versus robust adults (defined by slow-versus fast-TUG) across immune-cell subsets. Bars indicate shared or subset-specific genes; red and blue denote upregulation and downregulation, respectively (n=8). **Inset**: UMAP of baseline peripheral-blood immune cells from frail versus robust participants. (B) Cell-type-specific gene ontology enrichment in frail versus robust participants. Heatmap shows manually selected, significantly enriched GO terms associated with genes upregulated in frail relative to robust participants. These biological themes correlated with GeneAgent. Columns denote immune cell subsets; colour indicates enrichment score by clusterProfiler with values capped at 100 for visualization. GO term labels were abbreviated for readability, and coloured row annotations denote broader functional categories. (C) UpSet plot showing overlaps of downregulated genes, analysed as in (A). (D) Heatmap of significantly enriched GO terms associated with genes downregulated in frail versus robust participants, analysed as in (B). (E) Associations of plasma protein IFNγ–CXCR3 and inflammaging module scores with age-adjusted TUG residuals in the human discovery (n=21) and human validation cohorts (n=33). Each point represents an adult; lines and bands show cohort-specific linear fits and 95% confidence intervals. (F) Heatmap showing cell-type specific differential gene expression of IFNγ/CXCR3-associated response genes in frail versus robust participants. Rows denote selected genes and columns denote immune cell subsets; Asterisks denote FDR-adjusted significance: *FDR<0.05, **FDR<0.01, ***FDR<0.001.

Thus, physical frailty was associated with a compartmentalised immune-remodelling state in T cells and monocytes rather than a uniform pan-leukocyte transcriptional effect.

### Individuals who are physical frail had upregulated plasma IFNγ module expression

Given the enrichment of trafficking, inflammatory, and effector programmes across myeloid and T-cell subsets, we next asked whether related systemic immune signals were associated with physical frailty. We analysed two prespecified plasma-protein modules capturing inflammaging (CCL7, CCL3, and TNFSF14)^24^ and more specifically, IFNγ–CXCR3 signalling (IFNγ, CXCL9, and IL-18R1)^6^. In the discovery human cohort, both proteomic modules were positively associated with physical frailty (higher age-adjusted TUG residuals) after adjustment for CCI (IFNγ-CXCR3: β=0.53, 95%CI 0.04-1.03, p=0.036; inflammaging: β=0.55, 95%CI 0.07-1.02, p=0.028; Figure 2E; Table S3A). Findings were similar after additional adjustment for CMV serostatus. In the separate validation human cohort, both estimates remained positive but were attenuated and nonsignificant (Figure 2E; Table S3A). Sensitivity analyses with other physical measures, maximum handgrip strength and sit-to-stand tests, were directionally consistent (Table S3B).

Inverse-variance fixed-effect pooling supported a positive association for the IFNγ–CXCR3 module in the CCI-adjusted model (β=0.33, 95%CI 0.05-0.62, p=0.02, q=0.04) and after additional adjustment for CMV serostatus (β=0.33, 95%CI 0.05-0.61, p=0.02, q=0.04; Table S4). The inflammaging estimate was weaker and confidence intervals crossed zero (β=0.25, 95%CI -0.02-0.53, p=0.07). Bootstrap and influence analyses did not reverse the positive IFNγ-CXCR3 estimate (β=0.46; BCa 95%CI 0.07-0.93, p=0.01) nor identify a single observation that fully accounted for it (Tables S5 and S6), indicating that this was a stable association. We thus prioritised the IFNγ-CXCR3 module for downstream mechanistic study for three reasons: 1) its pooled association with frailty was larger than that of the broader inflammaging module; 2) it was more consistent across sensitivity analyses; and 3) it represents a defined signalling pathway, making it a more tractable therapeutic target.

At single-cell resolution, this IFNγ-response programme was engaged across many immune cell types in frail participants (Figure 2F). IFNγ-induced chemokines CXCL9, CXCL10 and CXCL11 were upregulated alongside interferon-stimulated and antigen-presentation genes (IFIT1/2/3, IFITM1, B2M) in T-cell subsets, monocytes and dendritic cells. Consistently, descriptive cell-level IFNγ-response scores were higher in frail participants in CD8 TEMRA and γδ T cells (Figure S4).

To test whether this elevated IFNγ signature reflected an antigen-specific expansion, we reconstructed TCR repertoires from paired 5′ single-cell VDJ libraries (23 samples; 93,443 cells with a productive TRB rearrangement, 81,508 with paired α/β). Clonotypes were defined by TRBV-CDR3β-TRBJ identity. Repertoires were polyclonal in 7 of 8 donors. The outlier donor was the single oligoclonal exception (clonality 0.34, largest clone 12.9% of T cells, D50=9) driven by a single private TRBV4-1/TRBJ1-5 clonotype, which had been reported in an unrelated COVID-19 cohort^25^ and consistent with an incidental public expansion rather than a frailty-specific response. Quantitative diversity metrics of the remaining 7 donors was low (normalised Shannon clonality median 0.07, range 0.04–0.12), the largest clone occupied only a median 2.9% of T cells (range 1.4–4.7%), and a median of 2,784 clones (range 1,685–4,885) was needed to account for half of all T cells. This diversity was inconsistent with an expansion dominated by one or a few antigen-specific clones. The expansions were stable across timepoints (16.2%, 12.8% and 10.4% at baseline, 3 months and 1 year), consistent with a chronic rather than acute response. This conclusion of no shared antigen-specific expansion holds with or without the outlier donor included.

To ask whether the expanded clones corresponded to known antigen-specific responses, we matched their CDR3β sequences against VDJdb, a curated database of TCRs with experimentally defined specificities. We focused on CMV, the dominant driver of CD8 TEMRA expansion in ageing^26^ ^27^. The repertoire was largely donor-specific and none of the expanded clonotypes matched known CMV-reactive TCRβ sequences (Figures S5B and C), indicating that the expanded effector cells were not driven by a single shared antigen and more specifically, not to CMV. This signal was more consistent with antigen-independent, cytokine-driven bystander activation.

Therefore, two independent data types from plasma and single-cell RNA sequencing analyses converge on the same signal of a systemic IFNγ–conditioned immune state present in frail individuals. As the associated CD8 TEMRA cells were polyclonal and lacked the clonal dominance or CMV-specificity expected of an antigen-driven expansion, this activation appears to be antigen-independent.

### CXCL9 was associated with a cytotoxic, terminal-effector immunophenotype

To identify downstream immune mechanisms linked to physical frailty, we focused on the IFNγ-CXCR3 module as a program-level representation of immune-inflammation. Within this module, CXCL9 had the largest pooled marker-level estimate (β=0.33, 95%CI 0.02-0.63, p=0.04; q=0.09; Table S7). After further adjustment for CMV serostatus and across the two independent human cohorts, CXCL9 remained consistently represented in individuals who are frail.

In circulating PBMCs, CXCL9 expression was sparse and largely present in monocytes (Figure S6A), whereas CXCR3 was upregulated across lymphocyte populations (e.g. activated CD4, naïve, central and effector memory CD8, CD8 TEMRA) (Figure S6B). CXCL9-response programmes, comprising CXCR3 signalling, downstream migratory and effector-response genes, were most evident in monocytes, activated and central-memory CD4 T cells, **CD8 TEMRA cells, and γδ T cells** (Figure 3A). Descriptive cell-level score distributions were higher in frail adults with upregulated CD8 TEMRA and γδ T cells (Figure 3B). These CD8 TEMRA and γδ T cells expressed higher cytotoxic-effector programmes (GZMB, GZMA, GZMH, GNLY, PRF1), lower CD28 and higher KLRG1, but not checkpoint/exhaustion genes (PDCD1, HAVCR2), which was characteristic of terminally-differentiated effector cells more than exhaustion (Figure S7). To test whether this reflected a shift along the CD8 differentiation trajectory, we ordered cells by pseudo-time. CD8 cells from frail participants were skewed toward a terminal-effector state (Figure 3C). Linking this trajectory to the CXCL9 axis, TCR analysis showed that expanded clonotypes within frail cytotoxic populations preferentially occurred in cells with higher CXCL9-response scores (Figure 3D). These CXCL9-responsive cytotoxic subsets were the same populations that showed the strongest transcriptional remodelling in our earlier single-cell analyses, reinforcing cytotoxic lymphocyte remodelling as a cellular correlate of frailty.

**Figure 3.**
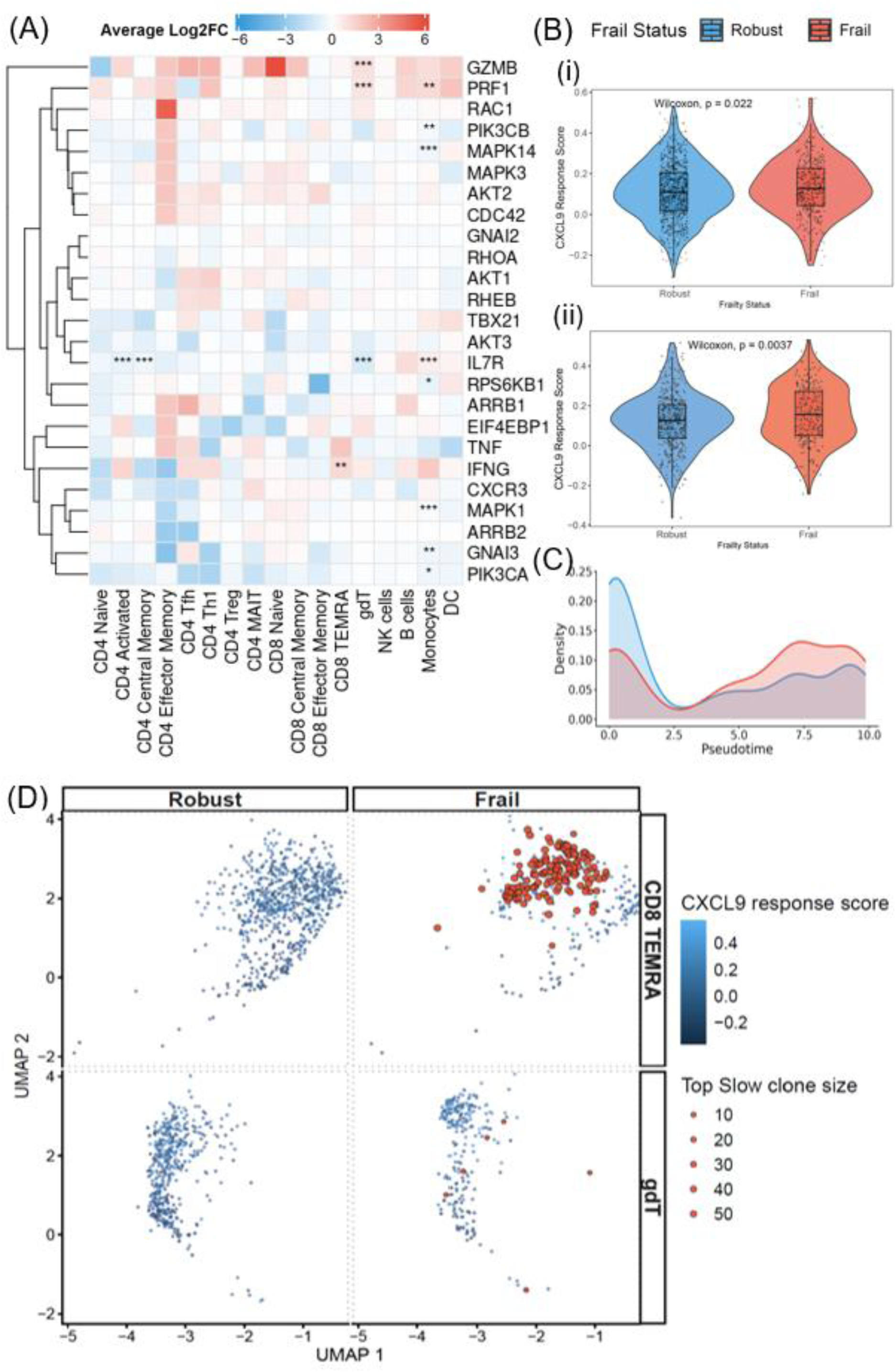
CXCL9-responsive cytotoxic lymphocyte states were enriched in frailty. (A) Heatmap showing cell-type specific differential expression of CXCL9-response genes in frail versus robust participants. Rows denote selected genes and columns denote immune cell subsets; colours indicate average log2 fold-change (red, higher in frail; blue, lower in frail). Asterisks denote FDR-adjusted significance: *FDR<0.05, **FDR<0.01, ***FDR<0.001. (B) Cell-level CXCL9-response scores in (i) CD8 TEMRA and (ii) γδ T cells by frailty group. Violin plots show descriptive single-cell distributions with embedded boxplots. Wilcoxon p values (CD8 TEMRA, p=0.022; γδ T cells, p=0.0037) are descriptive. (C) Density curves show a descriptive shift of cells from frail participants toward terminal pseudotime states. (D) UMAP overlay of the leading frail clonotypes on CXCL9-response scores across CD8 TEMRA, and γδ T cell compartments. TCR analyses remain descriptive due to the limited sample size. Blue intensity denotes CXCL9-response score; red points denote cells assigned to leading expanded clonotypes, with point size proportional to clone size. Leading clonotypes were donor specific. The panel does not establish a shared antigen-specific repertoire.

Flow cytometry provided protein-level support for this shift in cytotoxic T-cell subsets in frailty. In frailer participants, there were higher CD8 TEMRA and CD28-CD4-CD8-T-cell frequencies after adjusting for comorbidity and CMV serostatus, though this was insignificant after correction with the full flow panel (β = 0.55, 95%CI 0.05-1.05, p=0.03, q=0.62; β=0.57, 95%CI 0.12-1.02, p=0.017, q=0.61 respectively; Table S9A). No baseline CXCR3 measure was associated with frailty after correction (all BH q≥0.63; Table S9B and Figure S6C).

Thus, an IFNγ–CXCL9 programme was associated with the accumulation of terminally differentiated, cytotoxic CD8 TEMRA and γδ T cells in individuals who are physically frail.

### Muscle CXCL9 was associated with increased muscle T-cell and fibrosis module expression

We examined an independent adult cohort undergoing abdominal surgery and had their muscle biopsied (n=340; Table S10) to test whether the same IFNγ-CXCL9 axis was reflected in skeletal muscle. This cohort comprised different participants from the frailty cohorts and was phenotyped for sarcopenia defined by the Asian Working Group for Sarcopenia criteria^28^. Sarcopenia has been proposed as a biological substrate of physical frailty, and the two share low muscle strength and performance as core features^29^. Consistent with this relationship between sarcopenia and physical frailty, the systemic IFNγ-CXCR3 association was directionally consistent when indexed by maximum handgrip strength (Table S3B), a sarcopenia-defining measure, linking the systemic signal to the muscle-strength axis common to both. Because frailty was not assessed here with the same metric, sarcopenia status was treated as a covariate and effect-modifier. We then asked whether the same molecular axis recurred across these two independent cohorts. Our primary, pre-specified analyses were multivariate regression models relating immune and stromal gene modules to CXCL9 and fibrosis, adjusted for age, sex and sarcopenia status. Secondary interaction and sarcopenia-stratified analyses tested whether these pathway associations differed by sarcopenia status. Immune and stromal modules were specified a priori (Tables S11 and S12; Figure S8).

The cohort had a median age of 66 years (IQR 57-72), and 58.5% were male. 102 participants met criteria for sarcopenia^28^. In a multivariable model adjusted for age, sex, and sarcopenia status, muscle IFNγ-response activity was associated with leukocyte infiltration (β=0.40, 95%CI 0.30-0.49, p<0.001) and endothelial signal (β=0.37, 95%CI 0.28-0.47, p<0.001; adjusted R²=0.31). After accounting for these stromal modules, T-cell (β=0.41, 95%CI 0.31-0.51, p<0.001) and macrophage modules (β=0.10, 95%CI 0.00-0.19, p=0.05) remained significantly associated with residual IFNγ activity (Figure 4A; Table S13). Hallmark IFNγ single-sample gene set enrichment analysis (ssGSEA) gave similar results (Table S13). Exploratory lineage decomposition identified CD8A as the dominant measured T-cell correlate of IFNγ activity (β=0.37, 95%CI 0.28-0.46, p<0.001; Table S14). Because CD4 and CD8A were single-gene surrogates and the subset model only marginally improved fit, these results were secondary to the lineage-level analysis. Hence, after accounting for stromal signals, T cells were strongly and independently associated with muscle IFNγ activity, with exploratory analyses implicating CD8 as the leading T-cell subset.

**Figure 4.**
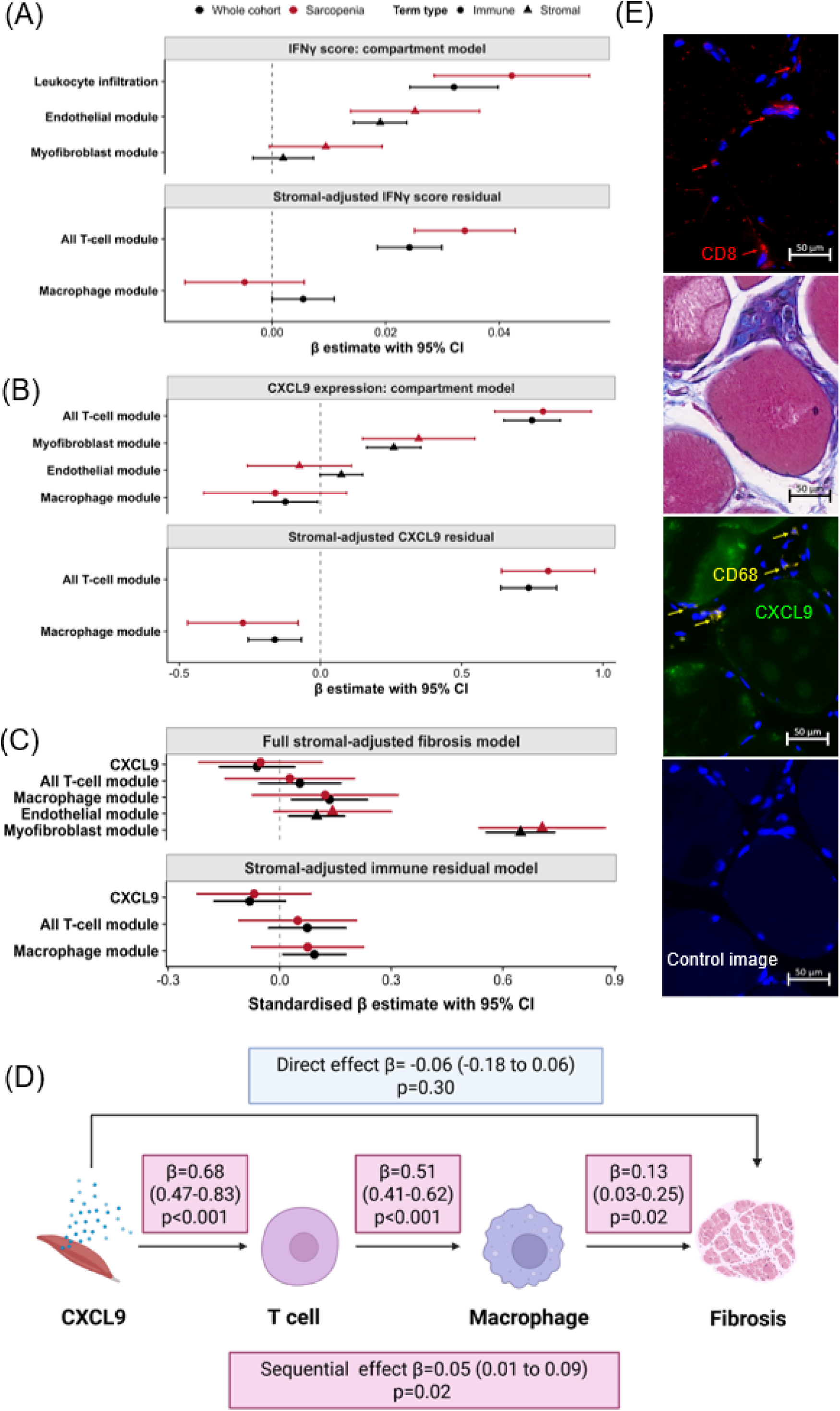
Muscle CXCL9 significantly associated with fibrosis through T-cell and macrophage signatures. (A) Forest plot of cellular correlates of the custom muscle IFNγ-response score in an immune and stromal compartment model and in a stromal-adjusted residual model. Points show standardised β estimates and horizontal bars 95% confidence intervals from multivariable linear models in the whole cohort (n=340) and sarcopenia-only subgroup (n=102); point shape denotes immune versus stromal terms. Whole-cohort models adjusted for age, sex, and sarcopenia status; subgroup models adjusted for age and sex. (B) Forest plot of cellular correlates of muscle CXCL9 expression, analysed as in (A). (C) Cellular and stromal correlates of the muscle fibrosis score. In the full model, macrophage (β=0.13, 95%CI 0.03-0.24, p=0.01), endothelial (β=0.10, 95%CI 0.02-0.17, p=0.010), and myofibroblast modules (β=0.65, 95%CI 0.56-0.74, p<0.001) were positive and significantly associated with fibrosis. In the stromal-adjusted immune-residual model, the macrophage module remained significantly associated with fibrosis residuals (β=0.09, 95%CI 0.01-0.18, p=0.03). (D) Cross-sectional sequential mediation model linking CXCL9 to T cells (0.68), to macrophages (0.51), and finally, macrophages to fibrosis (0.13). Standardised path coefficients and 95% confidence intervals are shown. The serial indirect effect was supported (β=0.05, 95%CI 0.01-0.09, p=0.02), whereas the direct effect CXCL9-to-fibrosis path was not (β= −0.06, 95%CI -0.18-0.06, p=0.30). The model evaluates pathway consistency, not temporal causality. Created with BioRender. (E) Representative sections of skeletal muscle from an individual with sarcopenia showing CD8 immunofluorescence (red) with DAPI nuclear counterstain (blue; top); Masson’s trichrome staining demonstrating endomysial and perimysial collagen deposition/fibrosis (blue; second from the top); and CD68 (yellow) and CXCL9 (green) immunofluorescence with DAPI (blue; second from the bottom). Images are from a single representative donor, stained on separate serial sections and not co-registered. Scale bars, 50 µm.

We then examined CXCL9, the chemokine our systemic data indicated to be the lead marker of the IFNγ-CXCR3 axis. Muscle CXCL9 expression was not associated with age or sarcopenia. Across the whole cohort, CXCL9 was strongly associated with the T-cell module (β=0.74, 95%CI 0.64-0.85, p<0.001) and more modestly with the myofibroblast module (β=0.25, 95%CI 0.14-0.37, p<0.001), whereas there was no independent association with macrophage (Figure 4B; Table S15). This CXCL9-T-cell association was stronger in sarcopenia (standardised slopes sarcopenia: β=0.80, 95%CI 0.68-0.91 versus non-sarcopenia: β=0.56, 95%CI 0.45-0.67; interaction β=0.23, 95%CI 0.08-0.39, p=0.003, BH q=0.017; Tables S16D and E). The T-cell module was the strongest independent correlate of CXCL9, accounting for 31.7% of its variance beyond other immune or stromal modules (β=0.68, 95%CI 0.58-0.77, p<0.001). An exploratory decomposition identified CD8A as the leading single-gene correlate (β=0.50, 95%CI 0.41-0.59, p<0.001), though the combined T-cell module fit the data substantially better than the CD4/CD8A subset model (Table S16).

Thus, CXCL9 expression was associated with a T-cell-rich muscle transcriptional signature, and this was more tightly linked in sarcopenic muscle.

Therefore, higher muscle IFNγ-response activity marked a mixed immune–stromal state, whereas muscle CXCL9 expression was most strongly associated with the T-cell signature and not with macrophages. This CXCL9-T-cell association was stronger in patients with sarcopenia.

### A CXCL9-T cell-macrophage-fibrosis pathway was seen from human muscle bulk RNA sequencing

We next asked whether human muscle upregulated CXCL9 expression was associated with muscle fibrosis, the histopathological finding linked to impaired muscle function^30^ ^31^. CXCL9 was associated with the muscle fibrosis score after adjustment for age, sex and sarcopenia (β=0.25, 95% CI 0.15–0.36, p<0.001). To test whether this association was direct or operated through intermediate cell populations, we additionally adjusted for the immune and stromal modules in our regression model. The CXCL9 association was fully attenuated (p=0.24), indicating the CXCL9-fibrosis link was explained by immune and stromal intermediate signals rather than acting directly.

We then examined which modules carried this link by testing the correlates of fibrosis directly. Fibrosis was most strongly positively associated with the myofibroblast module (β=0.65, 95% CI 0.56–0.74, p<0.001), significantly but more weakly with macrophage (β=0.13, p=0.01) and endothelial modules (β=0.10, p=0.01; adjusted R²=0.55). The macrophage association with fibrosis persisted against stromal-adjusted fibrosis residuals (β=0.09, p=0.03; Figure 4C; Table S17). An exploratory subset model demonstrated that resident macrophages had the strongest positive association with fibrosis (β=0.35, p<0.001), with a further positive contribution from recruited macrophages (β=0.22, p=0.01), whereas the inflammatory-macrophage module showed an inverse association (β=−0.25, p<0.001) with more inflammatory macrophages being linked with less fibrosis. These exploratory associations suggest that the fibrotic link is carried by reparative and resident macrophages rather than classical inflammatory ones.

Finally, we tested the full ordered pathway with a pre-specified serial mediation model (CXCL9 → T cell → macrophage → fibrosis). Each step was significantly positive. CXCL9 with the T-cell module (β=0.68, 95% CI 0.48-0.83), the T-cell module with the macrophage module (β=0.51, 0.42–0.62), and the macrophage module with fibrosis (β=0.13, 0.03-0.25). The serial indirect effect the path running through both intermediates was significant (β=0.05, 95% CI 0.01-0.09, p=0.02), whereas the direct CXCL9 to fibrosis path was not (p=0.30; Figure 4D; Table S19). Hence, the CXCL9-fibrosis association was statistically consistent with acting through sequential T-cell and macrophage signals rather than directly. An exploratory model using CD8A and a resident-macrophage signature reproduced the same indirect path (β=0.02, p=0.01; Table S21). Together, these cross-sectional analyses are consistent with a relay in which CXCL9 links T-cell and macrophage signals to muscle fibrosis. Sarcopenia did not alter these associations, and the sarcopenia-dependent amplification was confined to the CXCL9 to T-cell step (Tables S16, S20).

We proceeded to visualise this relationship *in situ*. Immunofluorescence supported the cell types and muscle fibrosis implicated by these analyses. Muscle from a sarcopenic individual showed interstitial CD8+ T cells and CD68+ macrophages, perimysial CXCL9 signal, and endomysial and perimysial collagen of fibrosis on Masson’s trichrome (Figure 4E). As serial sections were stained separately, these images show co-occurrence rather than quantified colocalisation.

Hence, in human muscle, upregulated CXCL9 and IFNγ-response activity were associated with T cells, and macrophage signatures with fibrosis, in a pattern consistent with a CXCL9 → T-cell → macrophage → fibrosis relay, with the CXCL9-T-cell coupling stronger in sarcopenic muscle. This tissue-level pattern mirrored the systemic finding of an IFNγ-CXCL9 programme enriched in CD8 TEMRA cells in frailty, providing convergent evidence across the blood and muscle compartments. To test whether these associated steps were individually plausible, we next examined each experimentally in primary human cell models.

### Primary human immune-muscle niche support CXCL9 → T cells → macrophages → fibrosis model

#### Myotube–fibroblast co-culture amplified IFNγ-induced CXCL9 secretion

Using a myotube-fibroblast co-culture model (Figure 5A, top panel), IFNγ-stimulation showed more CXCL9 than the sum of matched myotube-only and fibroblast-only cultures (82.97-190.25% above expected additive level; Figure 5B, Table S22), indicating synergistic stromal production.

**Figure 5.**
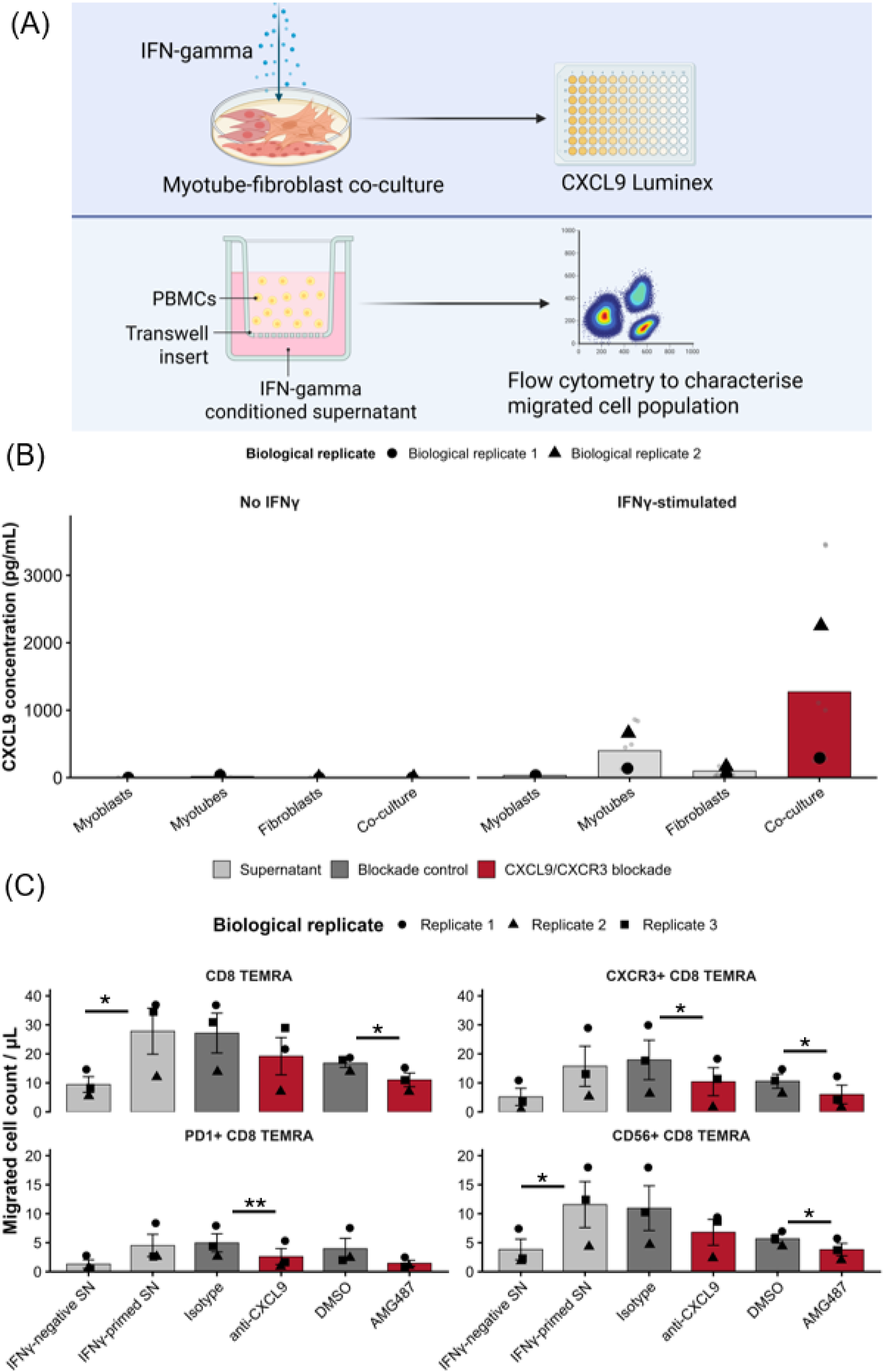
Primary human immune–muscle niche experiments support individual steps of a CXCL9-CD8-macrophage stromal-remodelling axis. (A) Experimental schematic. Created with BioRender. (B) CXCL9 secretion by primary human fibroblasts, myotubes, and myotube-fibroblast co-cultures (MT:FB 3:1) after 48h of IFNγ stimulation. Data are from n=2 biological replicates (circle, triangle) using the two distinct, separately expanded myotube and fibroblast donors; individual technical replicates (in grey) and condition means (bars) are shown. (C) PBMC transwell migration toward IFNγ-negative or IFNγ-conditioned myotube–fibroblast supernatant, with isotype/DMSO controls, CXCL9 neutralisation (anti-CXCL9) or CXCR3 antagonism (AMG487), shown for CD8 TEMRA, CXCR3+ CD8 TERMA, PD1+ CD8 TEMRA and CD56+ CD8 TEMRA populations as volume-normalised counts. IFNγ priming increased CD8 TEMRA (p=0.046) and CD56+ CD8 TEMRA migration (p=0.049). Anti-CXCL9 significantly reduced migration of CXCR3+ CD8 TEMRA (p=0.035) and PD1+ CD8 TEMRA (p=0.009), and AMG487 reduced migration of CD8 TEMRA (p=0.019), CXCR3+ CD8 TEMRA (p=0.031) and CD56+ CD8 TEMRA (p=0.016). Data from n=3 independent PBMC donors and n=2 myotube–fibroblast donor pairs, each with 2 technical replicates. Circle/triangle/square show per-donor means; bars show across-donor mean ± SEM. Statistics are one-sided paired t-tests on per-donor means. *p < 0.05.

#### Anti-CXCL9 and AMG487 reduced CD8 TEMRA migration toward IFNγ-conditioned stromal supernatant

We next reconstructed the CXCL9-T cell recruitment step using PBMC transwell migration toward IFNγ-conditioned myotube–fibroblast supernatant (Figure 5A, bottom panel). IFNγ-conditioned supernatant significantly increased migrated CD8 TEMRA cells from 9.44 to 27.81 (2.94-fold, p=0.046) and CD56+ CD8 TEMRA from 3.84 to 11.57 (3.01-fold, p=0.049). Relative to isotype control, neutralising anti-CXCL9 significantly reduced CXCR3+ CD8 TEMRA and PD1+ CD8 TEMRA migration (CXCR3+: -72.89%, 17.92 to 10.40 volume-normalised cells, p=0.03; PD1+: -90.60%, 4.97 to 2.61 volume-normalised cells, p=0.018 respectively). Relative to DMSO control, AMG487 (CXCR3 receptor antagonist) significantly reduced CD8 TEMRA, CXCR3+ CD8 TEMRA and CD56+ CD8 TEMRA migration (CD8 TEMRA: -52.30%, 16.80 to 11.03 volume-normalised cells, p=0.019; CXCR3+: -76.71%, 10.59 to 5.99 volume-normalised cells, p=0.031; CD56+: -49.17%, 5.69 to 3.81 volume-normalised cells, p=0.016), further substantiating the migration of CD8 TEMRA subsets was dependent on the CXCR3 pathway. Blocking either the ligand (anti-CXCL9) or the receptor (AMG487) reduced migration but did not abolish it. The concordant effect of both confirms a CXCL9-CXCR3-dependent component of recruitment (Figure 5C; Table S23), while the migration that persisted indicates that CXCL9– CXCR3 is one of several chemotactic signals rather than the sole driver.

#### CD8 co-culture shifts macrophages toward a pro-fibrotic secretome

To examine the downstream macrophage step, we co-cultured macrophages with CD8, and then introduced these macrophages into the myotube–fibroblast stroma (Figure S9A). Co-culturing with CD8 shifted macrophages toward a mixed (CD163+CD86+) and inflammatory (CD163-CD86+) phenotype, a consistent but non-significant trend across three donors (Figure S9B, Table S24). Adding these macrophages to the myotube-fibroblast stroma increased PAI-1 and total TGF-β1 secretion over 96h (Figure S9C, Table S25). Broader descriptive profiling of the macrophage-stromal secretome was consistent with this, showing highest CXCL9 secretion in the CD8-co-cultured macrophage condition at 96h and an accompanying rise in IL-6/IL-8 secretion (Figure S9D, Table S26).

Together, our primary human cell models support the upstream steps of the proposed pathway, CXCL9 → T-cells → macrophages → fibrosis. IFNγ-conditioned stroma produced CXCL9 synergistically, and CD8 TEMRA subsets migrate towards IFNγ-conditioned stromal supernatant in a partly CXCL9-CXCR3-dependent manner. The exploratory downstream macrophage arm demonstrated a shift towards a mixed-inflammatory macrophage phenotype following CD8 co-culture, and when added to the stromal co-culture, increased secretion of the matrix mediators of fibrosis PAI-1 and TGF-β1. These assays define the direction of effect at each step and show that the immune arm of the pathway, CXCL9 to T cell, converges on pro-fibrotic mediators whose downstream role in matrix remodelling is already established^32^ ^33^.

## Discussion

By integrating two human ageing cohorts, an independent muscle transcriptomic cohort, and primary human cell models, we found that frail participants had an upregulated systemic IFNγ-CXCL9 immune axis that was enriched in clonally expanded CD8 TEMRA cells. In the independent muscle cohort, the same axis was reflected in tissue as a CXCL9 → T-cell → macrophage → fibrosis pathway, with the CXCL9-T-cell association being stronger in sarcopenic muscle. In primary human cell models, the individual steps were supported experimentally: CD8 TEMRA subsets preferentially migrated towards IFNγ-conditioned myotube/fibroblast supernatant, with migration reduced by anti-CXCL9 and the CXCR3 receptor antagonist AMG487; CD8-co-cultured macrophages had a mixed-inflammatory phenotype and when added to myotube-fibroblast stroma, increased secretion of fibrosis mediators TGF-β1 and PAI-1.

Our first key finding was that IFNγ-CXCL9 axis was upregulated in both frail and a separate cohort of sarcopenic adults. In both the human discovery and validation cohorts, frail adults were associated with upregulated systemic IFNγ-CXCR3 module, while single-cell RNA transcriptomic analyses showed IFNγ-responsive, cytotoxic programmes particularly in CD8 TEMRA and γδ T cells in frail participants. Separately, upregulated IFNγ-response activity by bulk RNA sequencing of human muscle was significantly associated with leukocyte infiltration. These findings independently indicate that both the circulating immune system of frail adults and those with sarcopenia show an IFNγ-conditioned immune state. Prior literature is consistent, IFNγ-responsive chemokines are increased in inflammatory muscles^20^ ^21^ and can be induced directly in mouse models via acute injury^34^. In contrast, physical frailty and sarcopenia are characterised by chronic, low-grade inflammation^13^ ^31^ rather than the overt inflammatory disease or acute injury seen in current literature. Our data extends current understanding that in both frailty and sarcopenia, this sustained IFNγ-conditioned state may create a tissue environment for immune cell trafficking into muscle and shape muscle remodelling and fibrosis.

Our data also implicated bystander activation as the source for IFNγ. From our TCR data, the CD8 TEMRA population associated with the IFNγ-signal was polyclonal and lacked the clonal dominance expected of an antigen-specific response. We interpret this as antigen-independent, cytokine-driven bystander IFNγ production by terminally differentiated CD8 TEMRA cells, which can be triggered by innate cytokines such as IL-12, IL-15 and IL-18 in the absence of TCR engagement^35^ ^36^. Our macrophage-stromal co-culture data was consistent with such an environment: the IFNγ-conditioned, CD8-co-cultured macrophage niche accumulated pro-inflammatory IL-6 and IL-8, providing a cytokine-rich milieu which favours bystander IFNγ production. The persistence of the terminal-effector association after CMV serostatus adjustment further argues against simple CMV-driven CD8 TEMRA expansion as the explanation for this phenotype. This distinction is important. Most previous work on T cells in muscles come from autoimmune myositis, where cytotoxic CD8 T cells recognise and attack muscle fibres through specific muscle antigens^20^ ^37^. In ageing muscle, by contrast, the same IFNγ-producing effector machinery appears to be engaged without an antigen-specific autoimmune trigger. This bystander IFNγ production in frailty suggests that antigen-independent recruitment of terminally differentiated effector T cells may be a broader mechanism by which immune ageing contributes to end-organ remodelling, distinct from both antigen-specific responses to infection and organ-specific autoimmunity.

Within this broader IFNγ programme, CXCL9 emerged as a key trafficking signal for T cells into muscle. CXCL9 is strongly IFNγ-inducible and, through CXCR3, preferentially acts on activated lymphocytes^38–40^. In our first human cohort, plasma proteomics showed CXCL9 had the largest effect size to physical frailty within the IFNγ–CXCR3 module, while frailer adults had higher CD8 TEMRA and γδ T cells which showed stronger CXCL9-response programmes together with cytotoxic and terminal-effector features.

The bulk RNA sequencing of human muscle added complementary tissue-level observation. Significant association of CXCL9 and T cells was substantially stronger in sarcopenic muscle. What distinguished sarcopenic muscle was therefore not a higher level of CXCL9, but a tighter link between CXCL9 and T-cell infiltration, suggesting the tissue consequence of a CXCL9-rich environment depends on whether responsive T cells are present, not on CXCL9 abundance alone. As bulk transcriptomics cannot distinguish recruitment from local expansion or activation, this interaction should not itself be interpreted as direct evidence of trafficking. Our transwell experiments addressed this directly. IFNγ-conditioned myotube-fibroblast supernatant drove CD8 TEMRA subset migration. Blocking either the ligand (CXCL9 neutralisation) or receptor (chemical inhibition of CXCR3 receptor with AMG487^41^) significantly reduced this migration, indicating that recruitment was at least partially CXCL9/CXCR3-dependent. This apparent CXCL9/CXCR3 axis must be reconciled with reports that CXCR3 signalling can support, rather than harm, muscle. After acute injury, CXCL10 can promote the proliferation of muscle satellite cells and limit fibrosis^34^, whereas persistent CXCL9–CXCR3 signalling has been linked to fibrotic remodelling in other tissues^42^. Whether CXCR3 activity aids repair or sustains chronic inflammation therefore likely depends on which ligand predominates (CXCL10 versus CXCL9), which cell types are present to respond, and whether signalling is brief or sustained. Our data is within the chronic setting where the sustained, IFNγ-conditioned, CXCL9-rich niche acts on an expanded terminal-effector lymphocyte compartment.

Critically, these CD8 TEMRA subsets were the same terminal-effector populations our blood and muscle data had both nominated as the dominant IFNγ-expressing, clonally expanded compartment. Senescent-like CD56+ CD8 TEMRA cells can kill in an antigen-independent, NK-like manner^43^, and the PD1+ CD8 TEMRA cells in systemic circulation expressed effector and cytotoxic genes but limited expression of checkpoint genes, consistent with activation rather than exhaustion. We postulate that under the chronic conditions of ageing, these CXCL9-recruited activated, cytotoxic T cells can contribute to local muscle injury and the fibrotic remodelling we observed, though we did not directly test this downstream step. Instead, the strength of this inference lies in convergence. The same terminal-effector subsets recurred across independent blood and muscle cohorts and in the migration assay their recruitment depended on the CXCL9/CXCR3 axis inferred from those data, supporting the proposed circuit as connected.

The downstream consequence of this T-cell recruitment is matrix remodelling. Macrophages are central effectors of skeletal muscle fibrosis^30^ ^32^, a recognised consequence of failed muscle regeneration that contributes to the impaired contractile function^30^ ^31^ seen in sarcopenia, and CD8 T cells shape macrophage recruitment and phenotype in muscle^44^. This body of work, however, is largely set in acute injury, where CD8 T cells are pro-regenerative and support repair^44^. Our data indicate the direction of this relationship in a different setting. In ageing muscle, our cross-sectional models were consistent with a CXCL9-T-cell-macrophage relay, with macrophage signal associated with fibrosis after accounting for other immune and stromal modules. This sustained, CD8-associated macrophage state was hence associated with a pro-fibrotic output. This association in our human skeletal muscle cohort, together with the established role of macrophages in matrix remodelling,^30^ ^33^ is the principal basis for the macrophage arm of the pathway. Our cell experiments add preliminary, convergent support: CD8-educated macrophages showed a consistent shift from a reparative toward a mixed/inflammatory phenotype^45^ ^46^, which could be simultaneous repair and inflammatory programmes that become maladaptive when sustained^33^, and accumulated PAI-1 and TGF-β1 over time. PAI-1 limits plasmin-mediated matrix proteolysis and is linked to cellular senescence^47^, while TGF-β1 drives myofibroblast activation, consistent with the fibrosis^30^ reflected in the human muscle cohort. These co-culture data are preliminary and indicate direction of effect rather than causation. Overall, the findings are consistent with our immune-muscle axis in which recruited T cells shift macrophage state which leads to a matrix-remodelling secretome in muscle.

Our main strength is convergence across orthogonal datasets as no single association was sufficient to establish the proposed pathway. The IFNγ-CXCR3 module was thus mechanistically constrained based on cohort findings rather than derived post-hoc from a broad inflammatory screen. Our analysis distinguishes physical frailty from chronological age and treats sarcopenia as a related end-organ phenotype, and so the systemic-to-tissue link established convergence of a shared IFNγ-CXCL9 axis across syndromes and compartments rather than concordance of a single pathway within the same participants. These cohorts address the under-representation of Asian populations in ageing immunology. We used the cross-sectional mediation not as causal proof but to nominate and order the individual steps of the pathway. The primary in-vitro experiments supported the plausibility of these individual steps, including a secreted matrix-remodelling response. We prioritised antibody- and antagonist-based loss-of-function because these interrogate the same druggable nodes we propose as therapeutic targets, and we addressed specificity by orthogonal perturbation at two levels of the axis, ligand neutralisation (anti-CXCL9) and receptor antagonism (AMG487).

Our findings carry potential therapeutic implications. CXCR3 is druggable, with antagonists already in clinical development^48^, making this circuit a rational and tractable target for a geriatric syndrome that currently lacks any disease-modifying therapy. The stronger CXCL9– T-cell coupling in sarcopenic muscle further suggests that such interventions might be directed by tissue responsiveness depending on the individual’s frailty status.

Altogether, our data support an IFNγ-conditioned immune state in frailty that converges on CXCL9, linking cytotoxic terminal-effector T cells in the circulation to macrophage-associated matrix remodelling in skeletal muscle. To our knowledge, this is the first immune-muscle axis defined directly from human data for physical frailty, and it nominates CXCL9-CXCR3 as a druggable point of intervention.

## Declaration of AI use

Generative AI tools were used for language editing and troubleshooting R code for data analysis and figure preparation. All scientific content, study design, interpretation of results and final decisions were developed and verified by the authors.

## Acknowledgements and Funding

This work is supported by the National Medical Research Council (NMRC SEEDF23jul-0001; OFYIRG24jan-0013; CNIG25jan-0001), National Centre for Infectious Diseases (CAG-FY2024-014), Public Service Commission, Academy of Medicine and National University Health System (NCSP; CSDU). We thank Professors Bin Tean Teh, Weiping Han and Yibin Wang, theme investigators of the MAGNET programme, for their support (NMRC OFLCG24may-0020). VWTH is funded by NMRC RTF22jul-0010. ANA is funded by grants from The Medical Research Council grant (MR/T030534/1), Medical Research Council Grand Challenge in Experimental Medicine grant (MICA; MR/M003833/1), the LEO Foundation grant (LF-OC-19-000192) and the British Biotechnology and Biological Research Council grant (BB/Y003365/1). SX is funded by Ministry of Education (T2EP30125-0015). CWMO is funded by NMRC CSASI24jul-0005.

