## Supplemental Tables and Figures for "IFNγ-CXCL9-CXCR3 axis links T cell recruitment to skeletal muscle remodelling in human physical frailty"

### METHODS

#### Human subject recruitment

All studies involving human participants were approved by the National Healthcare Group Domain Specific Review Board (NHG DSRB 2021/00049; systemic immunophenotyping cohorts) the SingHealth Centralised Institutional Review Board (2020/2526; muscle cohort) and were conducted in accordance with the Declaration of Helsinki. Written informed consent was obtained before enrolment.

#### *Systemic circulation cohorts*

Two community-dwelling cohorts were studied (total n=55; Figure 1A). The discovery cohort included 22 participants (15 younger and 7 older adults) recruited from June to December 2021, and the validation cohort included 33 participants (13 younger and 20 older adults) was recruited from January 2023 to February 2024. The inclusion criteria for the human discovery and validation cohorts were community-dwelling adults aged 21 years or older; adults with active malignancy, immunosuppressed autoimmune disease, or known immunodeficiency were excluded. Functional testing and blood sampling were performed at baseline and longitudinal follow-up. Cohort characteristics are reported in Table S1.

#### *Muscle cohort*

The muscle cohort comprised of 340 patients undergoing colorectal surgery. Sarcopenia was classified using Asian Working Group for Sarcopenia 2019 criteria (102 sarcopenic; 238 non-sarcopenic). The cohort included 199 males (58.5%) and participants aged 22-90 years (Table S11). The inclusion criteria were patients undergoing elective colorectal surgery with intra-operative muscle biopsy of the rectus abdominis; participants with active malignancy in the biopsy site or active inflammatory myopathy were excluded.

#### Functional definition of physical frailty

Physical frailty was defined using the 3-metre timed up-and-go (TUG) test for a continuous index of mobility impairment<sup>1 2</sup> which is central to both physical frailty and sarcopenia. TUG is a performance-based assessment that integrates several physiological components relevant to the physical frailty phenotype, including lower-limb strength, balance, transfer ability, walking, turning, and motor coordination. It was therefore used to capture the locomotor expression of physical frailty most directly relevant to skeletal-muscle and immune–stromal biology. This choice was pragmatic and methodological. We have a locally prespecified threshold of $\geq 9.45$ s was used to define slow-TUG physical frailty<sup>1</sup>, and continuous age-adjusted TUG residuals could be used as the primary analytical outcome to preserve inter-individual variation and distinguish slower-than-expected physical performance from the effects of chronological age. This definition specifically captures the locomotor expression of physical frailty most directly relevant to skeletal muscle and immune–stromal biology, and does not represent multidomain frailty or deficit accumulation<sup>3</sup>. Sensitivity analyses were conducted for other physical tests used in physical frailty, namely maximum handgrip strength<sup>4</sup> and 5 times sit-to-stand test<sup>5</sup>. In contrast, categorical scales for physical frailty such as Clinical Frailty Scale<sup>6</sup> and FRAIL<sup>7</sup> score were considered but not appropriate as they were designed to discriminate across the full frailty spectrum including institutionalised and disabled older adults, and in a community sample they lack the dynamic range to detect graded biological associations.

Within each systemic immunophenotyping cohort, baseline TUG was regressed on chronological age, and the residual was used as the primary continuous mobility outcome. Positive values indicate slower-than-expected mobility. The dichotomous 9.45-s slow-TUG classification was used for descriptive group displays<sup>1</sup>. Continuous variables are reported as median (IQR) unless otherwise stated; categorical variables as n (%). Between-group comparisons of continuous variables were performed using Student's t-test where data were approximately normally distributed and Wilcoxon rank-sum test if otherwise. Categorical variables were compared using Pearson's $\chi^2$ test. Cohort-specific standardised $\beta$ estimates were pooled by inverse-variance fixed-effect meta-analysis. With only two small cohorts, pooled estimates were used to evaluate directional consistency rather than as definitive replication. Heterogeneity was summarised with Q and $I^2$. Bootstrap, influence, and leave-one-out analyses were treated as sensitivity analyses.

##### Peripheral blood mononuclear cell (PBMC) isolation and cryopreservation

Blood was collected into EDTA Vacutainer tubes (BD Biosciences) and centrifuged for 30 min at 400g to separate plasma from cells. Plasma was stored at -80°C. PBMCs were isolated by Ficoll-Paque Plus density-gradient centrifugation (Cytiva), cryopreserved in 90% heat-inactivated FBS (Hyclone) plus 10% dimethyl sulfoxide (Gibco), and stored in liquid nitrogen. Cells were thawed in pre-warmed RPMI 1640 (Gibco), washed twice, and used in the indicated assays.

##### Single-cell RNA sequencing and analysis

Cryopreserved PBMCs from eight participants (four younger and four older adults, Table S2) were profiled on recruitment and at one to three follow-up visits spanning approximately 3 months to 3 years, yielding 22 libraries. Single-cell suspensions of viability >80% were processed using the Chromium Next GEM Single Cell 5' Kit v2 (10x Genomics, PN-1000263) and T cell receptor amplification kits (PN-1000263) based on manufacturer's instructions. 10,000 cells were targeted per sample. Gene expression libraries were sequenced at a depth of 20,000 read pairs per cell, while TCR and BCR libraries were sequenced at 5,000 read pairs per cell. Libraries were sequenced on the Illumina NovaSeq X Plus platform using 150-bp paired-end reads. Only the baseline libraries were analysed for this study. QC parameters are provided in Methods Table M1.

Reads were aligned to GRCh38 with Cell Ranger v7.1.0 (10x Genomics) to generate feature–barcode matrices. Downstream analyses used R v4.5.3 and Seurat v5.4.0. Cells with >500 and <6,000 detected features and <20% mitochondrial transcripts were retained. Sample-level

quality-control plots are provided in Methods Figure M1. Data were log-normalised, 2,000 variable features were selected, and principal-component analysis used the first 30 components. Per-sample objects were integrated by Seurat anchor-based integration. Harmony was then applied using subject and timepoint covariates ($\theta=c(2,1)$; 30 dimensions; 20 iterations). The resulting Harmony-corrected embeddings were used for UMAP visualization, nearest-neighbour graph construction, and graph-based clustering at a resolution of 0.5.

#### *Cell-type annotation*

Initial labels were assigned with SingleR v2.12.0 using the Human Primary Cell Atlas. Automated annotations were manually reviewed against the expression of established canonical markers. Cells were subsequently assigned both broad-lineage and fine-subtype annotations. Cluster-specific marker genes were identified using Seurat's FindAllMarkers function with the Wilcoxon rank-sum test (`only.pos=TRUE`, `min.pct=0.25`, and `logfc.threshold=0.25`) and were used to support annotation decisions. Cells for which a confident annotation could not be assigned were classified as ambiguous and excluded from downstream cell-type-specific analyses.

#### *Gene-signature module scoring*

Single-cell module scores were calculated using Seurat's AddModuleScore function. Four predefined gene signatures were evaluated: a CXCL9/CXCR3-associated response signature comprising CXCR3, GNAI2, GNAI3, PIK3CA, PIK3CB, AKT1, AKT2, AKT3, RHEB, RPS6KB1, EIF4EBP1, RAC1, CDC42, RHOA, MAPK1, MAPK3, MAPK14, GZMB, PRF1, IFNG, TNF, IL7R, TBX21, ARRB1, and ARRB2; a T-cell effector signature comprising GZMB, GZMA, GZMK, GZMH, GNLY, PRF1, IFNG, EOMES, and TBX21; an inhibitory-receptor/exhaustion-associated signature comprising PDCD1, LAG3, TIGIT, and HAVCR2; and an IFNγ-response signature comprising STAT1, IRF1, CXCL9, CXCL10, CXCL11, GBP1, GBP2, IFITM1, IFI30, TAP1, TAP2, PSMB8, PSMB9, B2M, HLA-DRA, HLA-A, HLA-B, HLA-C, ISG15, IFI6, IFIT1,

IFIT2, and IFIT3. For each signature, the module score was calculated as the average expression of the signature genes relative to matched control genes selected from similar expression-level bins.

##### *Pseudotime analysis:*

CD8 T cells were converted from a Seurat object to a Monocle 3 (v1.4.27) cell\_data\_set, while retaining the existing UMAP coordinates and cluster assignments. A principal trajectory graph was learned in UMAP space using learn\_graph with partitions disabled. Cells annotated as naïve CD8 T cells were specified as the trajectory root, and pseudotime values were assigned using order\_cells. Pseudotime was mapped back to the Seurat metadata for visualization and comparison. Distributions of pseudotime between frail vs robust frailty groups were displayed using kernel-density plots after excluding missing or non-finite values.

##### *T-cell receptor repertoire analysis:*

Productive, high-confidence TRB contigs from D0 samples were matched to scRNA-seq metadata using sample-specific cell barcodes. Cells lacking complete CDR3, V-gene, or J-gene information were excluded, and clonotypes were defined by the combined V gene, J gene, and CDR3 amino-acid sequence. Clone size was calculated as the number of TRB-positive cells per clonotype within each T-cell subset and frailty group. Analyses focused on CD8 TEMRA and CD8 effector-memory cells, with $\gamma\delta$ T cells included in selected visualizations. Public clonotypes were detected in at least two donors, and putative CMV-associated clonotypes were identified by matching against VDJdb. CXCL9-response-high cells were defined by the top quartile of a Seurat module score. All TCR analyses were considered descriptive because of the limited sample size.

Analyses: Group comparisons were descriptive. Primary inferential analysis was donor-level, these analyses were performed for cell-type proportions, DEGs, CD8 pseudotime and TCR

analyses. For descriptive single-cell-level comparisons, scores were compared between frail and robust groups within each cell type using the Wilcoxon rank-sum test.

#### Protein quantification

##### *OLINK*

Frozen plasma was analysed in two batches by the OLINK service laboratory using the Target 96 Inflammation proximity-extension assay. Normalised Protein eXpression (NPX) values are reported on a log2 scale. Samples with OLINK QC-warning flags were excluded, and proteins with >50% of values below the limit of detection (LOD) were removed. For the remaining 89 proteins, below-LOD values were retained at the assay LOD. Bridge samples were used to assess and correct plate effects. Proteins were z-standardised before calculation of prespecified module means. Ranking considered pooled effect magnitude and precision, cross-cohort direction, multiplicity-adjusted q value, heterogeneity, and biological interpretability. CXCL9 was treated as an exploratory representative marker because $q=0.08$.

##### *Luminex bead array*

Frozen cell culture supernatants were analysed for CXCL9, PAI-1, CXCL8, IL-6 and IFNγ using customised ProcartaPlex assays (Thermofisher Scientific; EPX01A-1028 5-901; customised Procartaplex PPX-03) according to the manufacturer's instructions. Technical replicate numbers are reported for each experiment and figure.

##### *TGF-β1 ELISA*

Frozen supernatants were analysed for total TGF-β1 with ELISA (Thermofisher Scientific; BMS249-4) according to the manufacturer's instructions. Technical replicate numbers are reported for each experiment and figure.

Each protein was z-standardised within the analytical dataset, module members were averaged, and the resulting score was re-standardised. Prespecified OLINK modules were

IFNγ-CXCR3 (IL-18R1, IFNγ, CXCL9) and inflammaging (CCL7/MCP-3, CCL3, TNFSF14). Associations with age-adjusted TUG residuals were estimated by multivariable linear regression. Primary models adjusted for CCI; secondary models also adjusted for CMV serostatus. Two-sided paired t-tests were used for Luminex and ELISA readouts.

##### CMV IgG ELISA

CMV IgG serostatus was measured with the Abcam Human anti-CMV IgG ELISA kit (ab108724). Plasma was diluted 1:100, incubated in CMV-antigen-coated wells, and detected with HRP-conjugated anti-human IgG and tetramethylbenzidine. Absorbance was read at 450 nm with a 620-nm reference. Manufacturer-defined index thresholds were used: >1.1, positive; 0.9–1.1, equivocal; and <0.9, negative. Each plate was accepted only when all manufacturer-specified control criteria were met; failed plates were repeated.

##### Flow cytometry

High-dimensional flow cytometry was performed on discovery-cohort PBMCs, migrated transwell cells, and macrophage-polarisation samples. Cryopreserved PBMCs were thawed and washed in FACS buffer (PBS, 2% FBS, 1 mM EDTA); transwell and macrophage assays used fresh cells. Cells were stained with the panels in Methods Tables M2–M4 and Live/Dead Fixable Near-IR dye, resuspended in 300 μL FACS buffer, and acquired on a Cytex Aurora. UltraComp eBeads single-stain controls were used for spectral unmixing in SpectroFlo; fluorescence-minus-one controls were used to set CXCR3 gates in migration assays. Data were analysed in FlowJo v10. Gating strategies are shown in Methods Figures M2–M4. Flow-cytometry frequencies and fluorescence intensities were related to the age-adjusted TUG residual by linear regression adjusted for CCI and CMV serostatus. Predictors were standardised for effect-size comparison. Benjamini–Hochberg correction was applied across the full tested flow panel (Methods Table M2).

##### Bulk RNA sequencing of human muscle

Approximately 1 cm<sup>3</sup> of rectus-abdominis muscle was collected intra-operatively at the specimen-extraction site, snap-frozen in liquid nitrogen, and stored at -80°C. Total RNA was extracted using TRIzol/chloroform according to the manufacturer's protocol. Bulk RNA-sequencing data were pre-processed using the nf-core/rnaseq pipeline v3.10.1<sup>8</sup>. Reads were aligned to the GRCh38/hg38 reference genome using STAR v2.7.10a<sup>9</sup> with GENCODE v38 gene annotations<sup>10</sup>, and gene expression was quantified with RSEM v1.3.1<sup>11</sup>. Library quality was assessed using RSeQC v3.0.1<sup>12</sup>, Qualimap v 2.2.2-dev<sup>13 14</sup> and MultiQC<sup>15</sup>, including sequencing depth, mapping rate, gene-body coverage, duplication and strandedness metrics. Samples failing prespecified quality-control criteria were excluded before downstream analysis. Expression values were corrected for sequencing batch effects across sequencing runs using ComBat<sup>26,27</sup>. Enriched Hallmark pathways in each sample were identified using ssGSEA (v10.1.0)<sup>16 17</sup>.

The custom IFNγ-response module comprised 23 genes: STAT1, IRF1, CXCL9, CXCL10, CXCL11, GBP1, GBP2, IFITM1, IFI30, TAP1, TAP2, PSMB8, PSMB9, B2M, HLA-DRA, HLA-A, HLA-B, HLA-C, ISG15, IFI6, IFIT1, IFIT2, and IFIT3. It correlated with the Hallmark IFNγ-response ssGSEA score ($r=0.84$). The fibrosis score comprised COL1A1, COL1A2, COL3A1, FN1, CTGF, ACTA2, TGFB1, and VIM.

A priori lineage and stromal modules were: T cell (CD3E, TRAC, TRBC2, CD2, CD247), macrophage (CD14, ITGAM, CSF1R, LST1, AIF1, TYROBP, FCER1G, SPI1, CD68), myofibroblast (POSTN, LOX, TNC, FAP), and endothelial (PECAM1, VWF, CDH5). Coherence was evaluated with Cronbach $\alpha$, mean inter-gene correlation, and first-principal-component variance (Table S14). PTPRC, CD4, and CD8A were treated as single-gene readouts. Genes did not overlap across the primary modules. Exploratory macrophage-state modules were defined as recruited (CCL2, CCL4, CCL8, FCN1, VCAN, S100A8, S100A9), resident (FOLR2, LYVE1, STAB1, SELENOP, C1QA, C1QB, C1QC), inflammatory (IL1B, TNF, IL6, NLRP3, CXCL2, CXCL3, PTGS2), and reparative-associated (MSR1, MERTK, CCL18,

MAF, CD163, MRC1). Related modules were fitted separately because of collinearity and were not used in the primary mediation model.

Multivariable linear regression was adjusted for age, sex, and sarcopenia status in the whole cohort and age and sex in the sarcopenic subgroup. Exploratory effect modification by sarcopenia was assessed using interaction terms between sarcopenia status and the focal exposure; group-specific slopes were estimated from the corresponding interaction models, with BH correction across the five interaction tests. Collinearity was evaluated with pairwise and partial correlations and variance-inflation factors. The prespecified serial model was CXCL9 → T-cell module → macrophage module → fibrosis score, fitted with lavaan v0.6-19. Bias-corrected bootstrap confidence intervals were generated for specific indirect, serial indirect, direct, total indirect and total effects using 5000 resamples. Because exposure, mediators, and outcome were measured cross-sectionally from the same bulk transcriptome, mediation results are interpreted as pathway consistency, not temporal or causal mediation.

#### Immunofluorescence

Serial sections of rectus-abdominis muscle biopsy from a representative sarcopenic donor were used to relate immune-cell localisation to fibrosis. Formalin-fixed, paraffin-embedded blocks were cut at 5 µm; adjacent serial sections were assigned to Masson's trichrome staining, immunofluorescence for CXCL9 and CD68, and CD8a immunofluorescence.

Sections were then deparaffinised, rehydrated, and subjected to heat-induced epitope retrieval in EDTA buffer (pH 9.0). Sections were permeabilised in 0.1% Tween-20 in PBS and blocked in 5% BSA for 1 h at room temperature, then incubated overnight at 4 °C with primary antibodies against CD8a (clone 4SM16, rat, 1:25, Invitrogen), CD68 (clone KP1, mouse, 1:200, DAKO) and CXCL9 (clone 11H1L14, rabbit, 1:200, Invitrogen). Sections were washed and incubated for 2h at room temperature with species-matched, high affinity cross-adsorbed

fluorophore-conjugated secondary antibodies (Alexa Fluor 647 for CD8, CY3 for CD68, and Alexa Fluor 488 for CXCL9). Nuclei were counterstained with DAPI and slides were mounted in DAKO antifade mounting medium. To control for muscle autofluorescence in the green channel, a secondary-only antibody control section was processed in parallel and imaged at identical exposure and gain settings to the CXCL9 acquisition; the green signal was negligible in this control (Figure 4E). Masson's trichrome was performed on the adjacent serial section according to the manufacturer's protocol (Abcam).

##### *In-vitro* human cell models

Ethical approval was obtained for use of primary human cells (NUS IRB LN-20-054E). Human primary skeletal muscle myoblasts and primary dermal fibroblasts were maintained in their respective growth media (F10+20% FBS for myoblast, DMEM+10% fetal bovine serum (FBS) for fibroblast) at 37°C in 5% CO<sub>2</sub>. To induce myotube differentiation, myoblasts at 80–90% confluence were switched to DMEM without added FBS for 5 days, with confirmation of multinucleated myotube formation by light microscopy. All cells were used between passages 5-10 and passages were recorded from the first time the original cell vials were thawed. Myotube formation was detected by observing multinucleated cells under light microscopy. Primary myoblasts and fibroblasts were each derived from two independent donors; myoblasts and fibroblasts were obtained from different individuals.

Differentiated myotubes, primary fibroblasts, and myotube–fibroblast 3:1 co-cultures were seeded in 6-well plates and maintained in DMEM plus 10% FBS for 24h (Figure 5A, top panel). Cultures were then exposed to recombinant human IFNγ (10 ng/mL; R&D Systems, 285-IF-025) or sterile water for 48 h. Supernatants were collected and stored at –80°C until use. Technical replicates were first averaged within each independent experiment. The expected additive CXCL9 concentration was calculated as the sum of CXCL9 concentrations measured in matched IFNγ-stimulated myotube-only and fibroblast-only cultures. Excess over additivity was defined as the observed CXCL9 concentration in the myotube–fibroblast co-culture minus this additive expectation, and percentage excess was calculated as $[(\text{co-culture} - \text{additive expectation}) / \text{additive expectation}] \times 100$. Excess over additivity was tested against zero across independent biological replicates using two-sided T-test.

For the transwell assays (Figure 5A, bottom panel), we used three independent PBMC donors. PBMC migration was assessed in 24-well Transwell plates with 5-$\mu\text{m}$ polycarbonate membranes (Corning 3421). The lower chamber contained 600 $\mu\text{L}$ of the indicated control medium, recombinant CXCL9 or IFNγ control, unstimulated or IFNγ-conditioned myotube–fibroblast supernatant, or conditioned supernatant with anti-CXCL9, matched isotype, or AMG487. Anti-CXCL9 (5 $\mu\text{g}/\text{mL}$; R&D Systems MAB392-SP) or isotype (mouse IgG1; in-house) was pre-incubated with supernatant for 30 min. PBMCs were pretreated with AMG487 (10 $\mu\text{M}$; MedChemExpress HY-15319) for 30 min at 37°C. PBMCs ($7.5 \times 10^5$ in 100 $\mu\text{L}$) were added to the upper chamber for 6 h at 37°C and 5% CO<sub>2</sub>. Migrated cells were collected and characterised by flow cytometry using Flowjo 10 (Figure M3, Table M3). Transwell counts were volume-normalised. Given the apriori hypothesis that IFNγ-conditioned supernatant increases migration, one-sided paired t-tests were conducted.

CD8 T cells and monocytes were magnetically isolated from three PBMC donors, matched to the transwell assays. Monocytes were differentiated for 7 days with M-CSF (50 ng/mL; PeproTech 300-25). CD8 cells were activated for 24 h with plate-bound anti-CD3 (2 $\mu\text{g}/\text{mL}$; Thermo Fisher Scientific 16-0037-81) and soluble anti-CD28 (2 $\mu\text{g}/\text{mL}$; Thermo Fisher Scientific 16-0289-85). Macrophage conditions were: M0, M-CSF only; IL-4 control (20 ng/mL); activated-CD8 co-culture at a 10:1 CD8:macrophage ratio; CD8 co-culture with anti-IFNγ (5 $\mu\text{g}/\text{mL}$); and CD8 co-culture with matched mouse IgG2a isotype (in-house). These monocyte-derived macrophages were used as a tractable in vitro proxy and were not assumed to reproduce resident muscle macrophages.

On day 9, cells were washed, incubated with pre-warmed Accutase (STEMCELL Technologies 07922) for 20 min at 37°C, dissociated by pipetting after addition of cold medium, washed, and resuspended. Immune cells were characterised by flow cytometry (Figure M4, Table M4). Immune cells were added to myotube–fibroblast monolayers at a 1:3 immune:stromal ratio in 1:1 DMEM:RPMI with 10% FBS. This deliberately immune-enriched ratio was used as an inflammatory stress test and is not intended to reproduce physiological cell abundance. M-CSF (25 ng/mL) was replenished in macrophage conditions at 72 h. Supernatant was collected at 24 and 96 h (Figure 9A). Luminex and ELISA cytokine readouts were normalised by cell count and averaged across replicates.

#### Statistics

All analyses were conducted in R (v4.5.3). P-values and 95% confidence intervals were reported. Benjamini–Hochberg correction was applied within prespecified analysis families, with nominal but non-FDR-significant results identified explicitly. Sample sizes were feasibility based rather than determined by formal power calculations. Significance was defined as $p < 0.05$, results are reported to 2 decimal places.

#### Data and code availability statement

Raw scRNA-seq data and the analysis code are available here: http://github.com/lincsii/IFNg-CXCL9-CXCR3-frailty.git.

**Methods Table M1.**

| **ID** | **Cell Count** | **Total UMI** | **Median Unique Molecular Identifier/Cell** | **Median Genes/Cell** | **Median Mitochondrial Percent** |
| --- | --- | --- | --- | --- | --- |
| 10 | 2020 | 10566151 | 4444.5 | 1952.5 | 3.39 |
| 11 | 1267 | 5946460 | 4489 | 1848 | 2.91 |
| 13 | 1876 | 7968623 | 3959 | 1719 | 3.25 |
| 14 | 3407 | 12411486 | 3100 | 1369 | 3.05 |
| 15 | 1655 | 8698679 | 4427 | 1926 | 2.86 |
| 18 | 658 | 3278143 | 5013.5 | 1958 | 3.47 |
| 6 | 1727 | 7495686 | 4067 | 1726 | 2.49 |
| 9 | 1975 | 9542376 | 4270 | 1885 | 3.02 |

**Methods Table M2. Surface-staining panel for systemic T cells**

| **Marker** | **Fluorochrome** | **Clone** | **Manufacturer** | **Catalogue number** |
| --- | --- | --- | --- | --- |
| CD14 | APC-Cy7 | Mfp9 | BD, Sparks, MD, USA | 557831 |
| CD19 | APC-Cy7 | SJ25C1 | BD, Sparks, MD, USA | 557791 |
| CD3 | BUV395 | UCHT1 | BD, Sparks, MD, USA | 563546 |
| CD4 | BUV496 | SK3 | BD, Sparks, MD, USA | 612936 |
| CD8 | PerCP-Cy5.5 | SK1 | BD, Sparks, MD, USA | 565310 |
| CXCR3<br>(CD183) | PE | 1C6 | BD, Sparks, MD, USA | 557185 |
| PD-1 | BV711 | EH12.2H7 | Biolegend, San Diego, California, USA | 329928 |
| CD45RA | BV510 | HI100 | Biolegend, San Diego, California, USA | 304142 |
| CD27 | AlexaFluor700 | O323 | Biolegend, San Diego,<br>California, USA | 302814 |
| CD28 | BUV805 | CD28.2 | BD, Sparks, MD, USA | 742037 |

**Methods Table M3. Surface-staining panel for migrated transwell cells**

| Marker | Fluorochrome | Clone | Manufacturer | Catalogue<br>number |
| --- | --- | --- | --- | --- |
| CD3 | BUV395 | UCHT1 | BD, Sparks, MD, USA | 563546 |
| CCR7 | BUV563 | B-LY6 | BD, Sparks, MD, USA | 749679 |
| KLRG1 | BUV661 | Z7-205 | BD, Sparks, MD, USA | 753693 |
| CD45 | BUV805 | HI30 | BD, Sparks, MD, USA | 612891 |
| CD45RA | BV510 | HI100 | Biolegend, San Diego,<br>California, USA | 304142 |
| CD8 | RV544 | RPA-T8 | BD, Sparks, MD, USA | 573421 |
| PD-1 | BV711 | EH12.2H7 | Biolegend, San Diego,<br>California, USA | 329928 |
| CD56 | BV786 | NCAM16.<br>2 | BD, Sparks, MD, USA | 564058 |
| CD57 | FITC | HNK-1 | Biolegend, San Diego,<br>California, USA | 359603 |
| CXCR3<br>(CD183) | PE | 1C6 | BD, Sparks, MD, USA | 557185 |

**Methods Table M4. Surface-staining panel for macrophages**

| Marker | Fluorochrome | Clone | Manufacturer | Catalogue<br>number |
| --- | --- | --- | --- | --- |
| HLA-DR | BV650 | G46-6 | BD, Sparks, MD,<br>USA | 564231 |
| CD163 | PE-CF594 | GHI/61 | BD, Sparks, MD,<br>USA | 562670 |
| CD45 | BUV 496 | HI30 | BD, Sparks, MD,<br>USA | 750179 |
| CD11b | BV785 | ICRF44 | Biolegend, San<br>Diego,<br>California, USA | 301346 |
| CD86 | BUV 737 | FUN-1 | BD, Sparks, MD,<br>USA | 612785 |

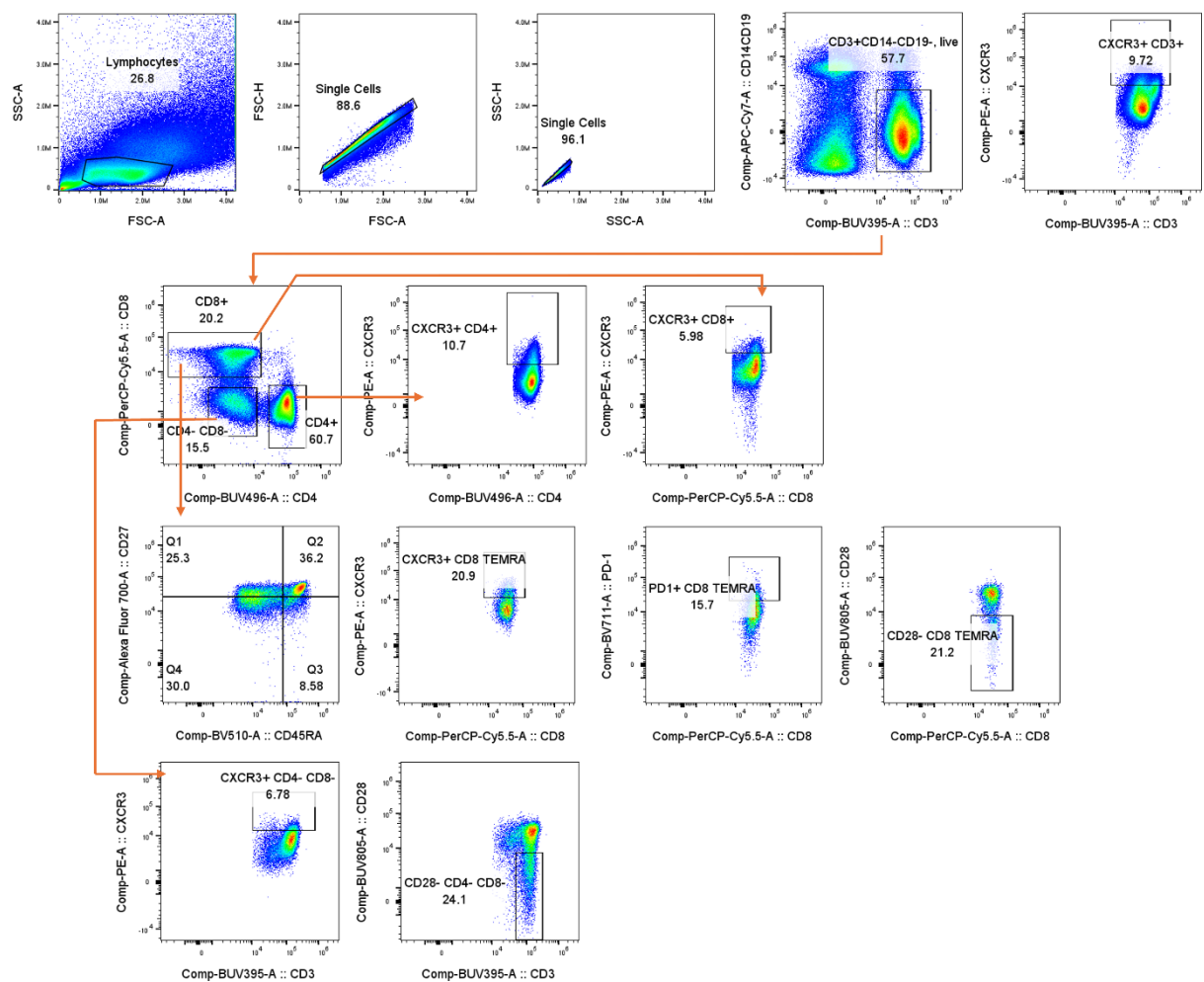

**Methods Figure M2. Gating strategy for systemic T cells**

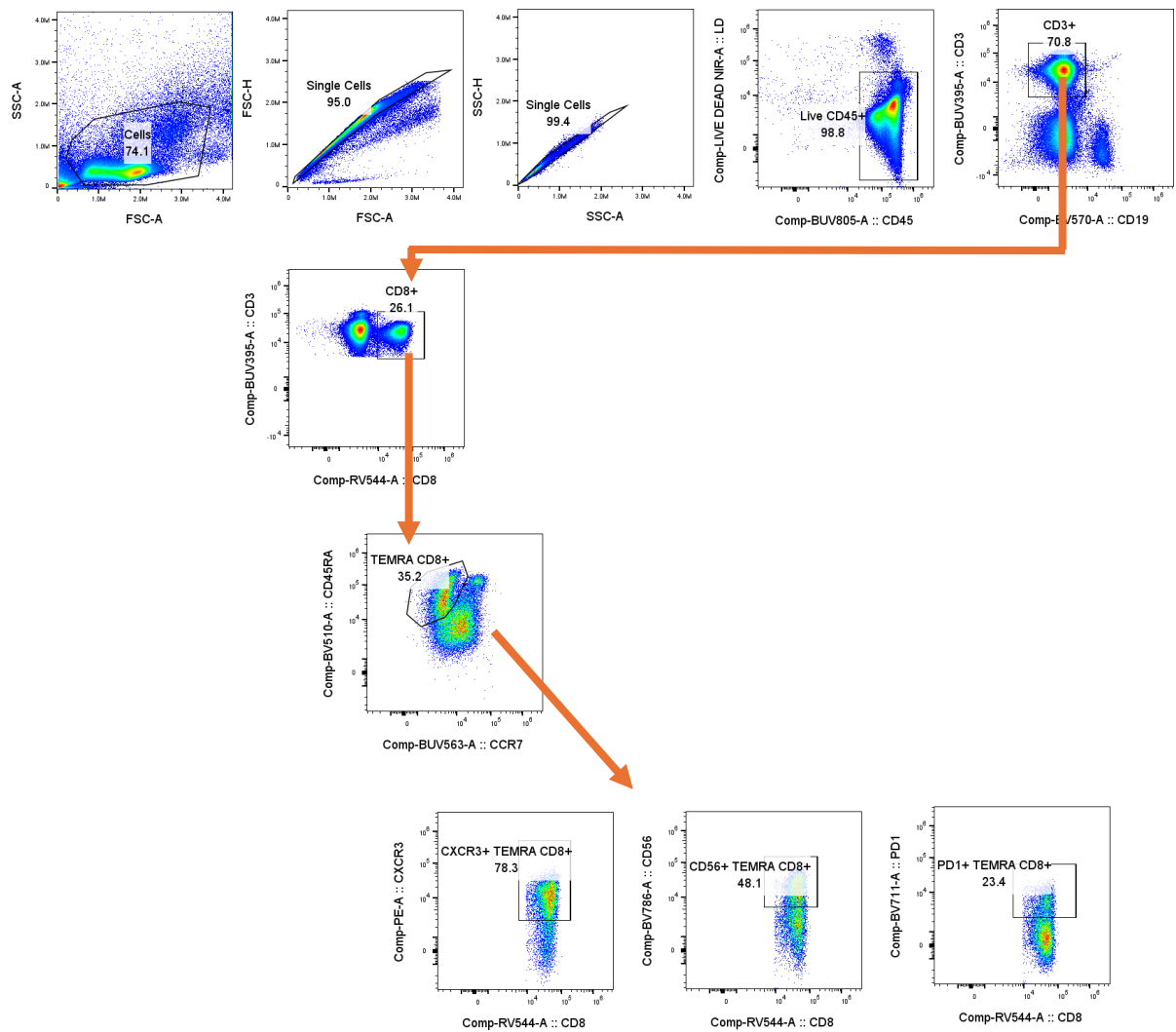

**Methods Figure M3. Gating strategy for migrated transwell cells**

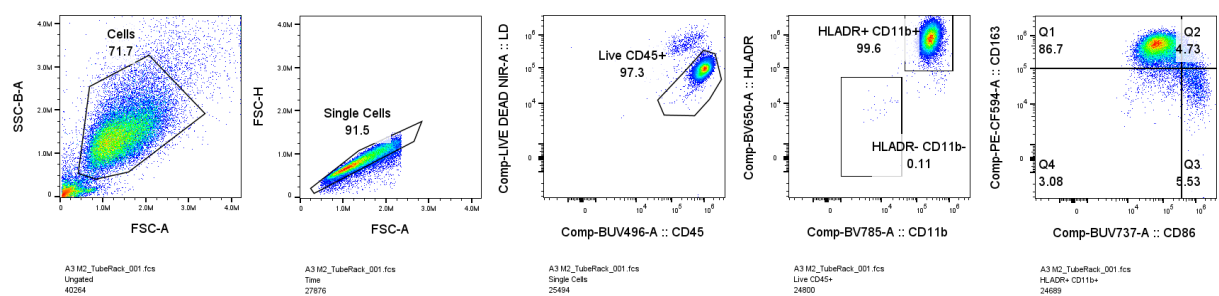

**Methods Figure M4. Gating strategy for macrophages. HLA-DR+CD11b+ macrophages were resolved into CD163/CD86 quadrants.**

### Supplementary Figures

(A)

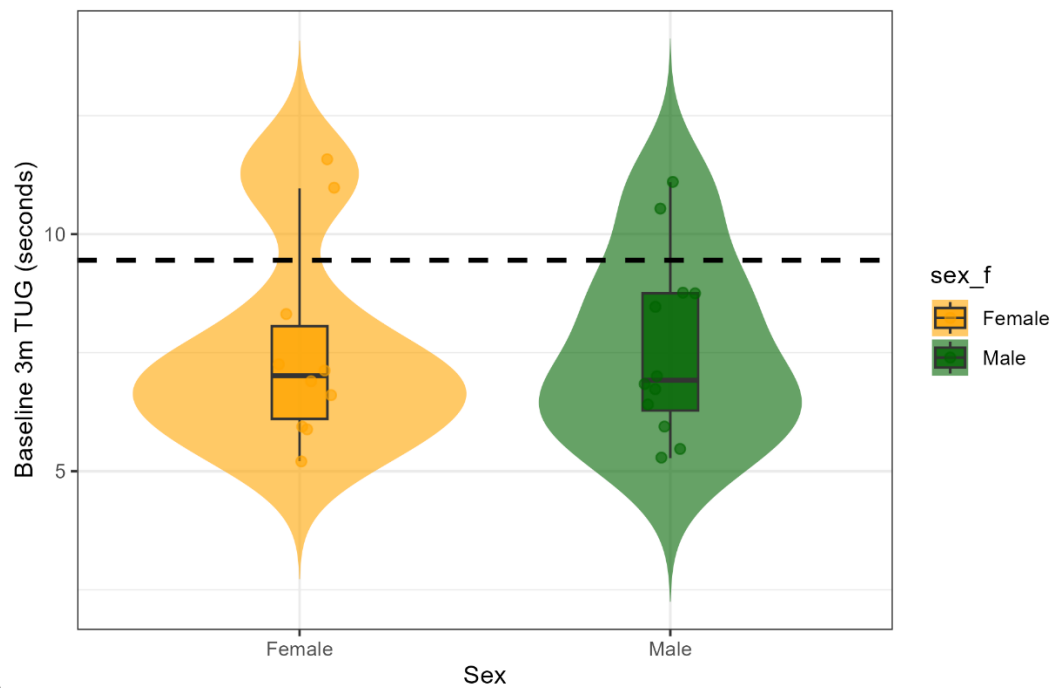

(B)

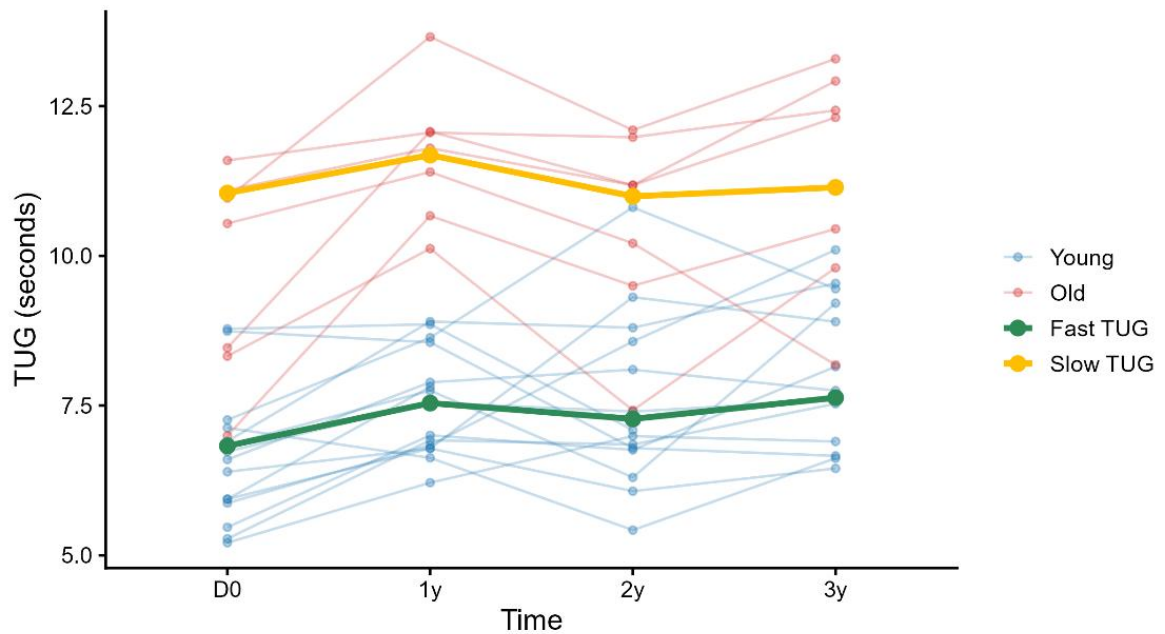

**Figure S1. Baseline TUG distributions and longitudinal trajectories in the discovery cohort.** (A) Baseline 3-min TUG by sex. The dashed line denotes the 9.45-s slow-TUG threshold. (B) Participant-level TUG trajectories across visits, with the thick line representing group mean. Points represent participants. Colours identify the groups shown in each panel.

(A)

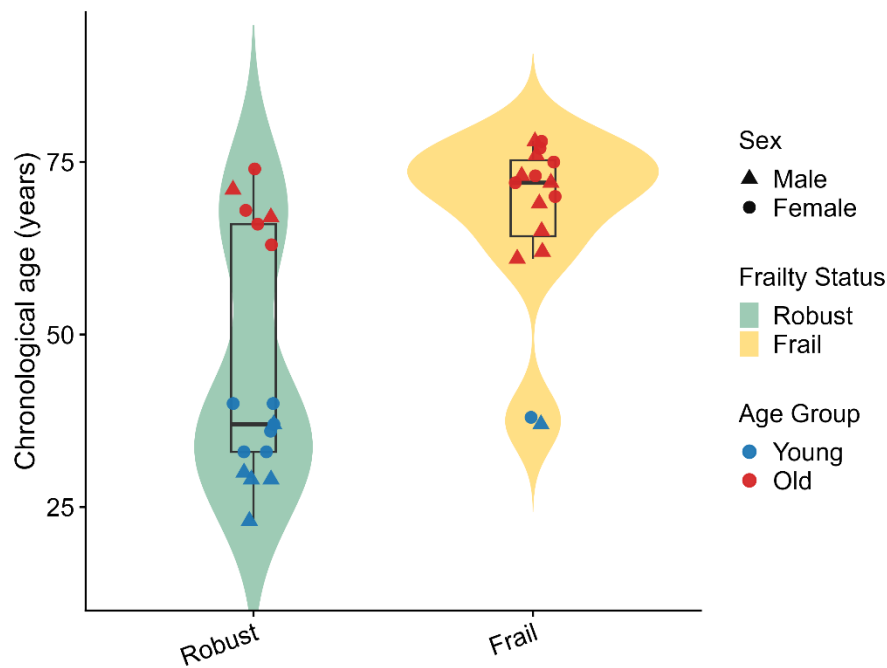

(B)

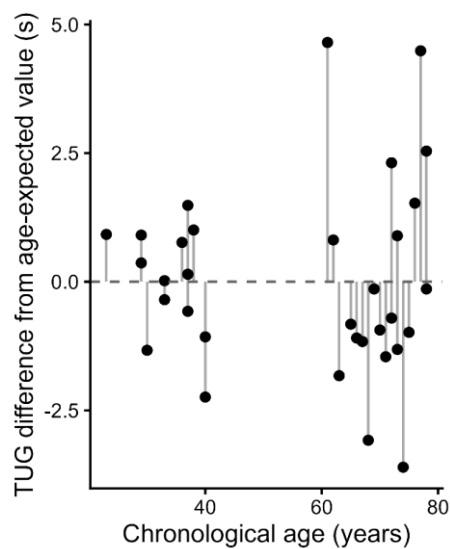

**Figure S2. Baseline TUG and age residualisation in the validation cohort.** (A) Baseline TUG by age and sex. Young adults had faster TUG than older adults (8.33 [7.89–8.87] versus 10.20 [9.42–12.10] s, $p < 0.001$). Age was associated with baseline TUG ($\beta = 0.07$, 95% CI 0.03–0.10, $p < 0.001$), whereas sex was not ($p = 0.35$). (B) TUG differences from age-expected values derived from a linear model of baseline TUG on chronological age;

positive values indicate slower-than-expected TUG. The model explained $R^2 = 0.35$ (adjusted $R^2 = 0.31$); residualised TUG was uncorrelated with age ($r = 0.00$, 95% CI $-0.34$ to $0.34$, $p = 1.00$).

(A)

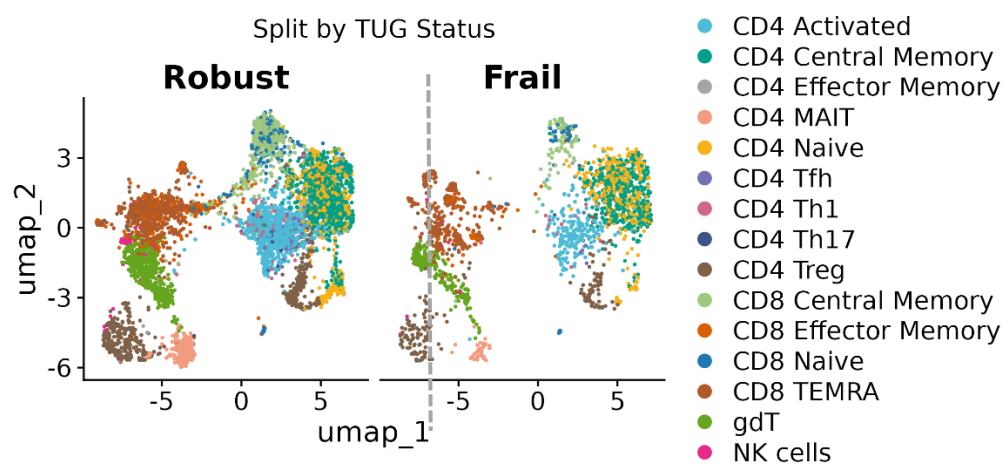

(B)

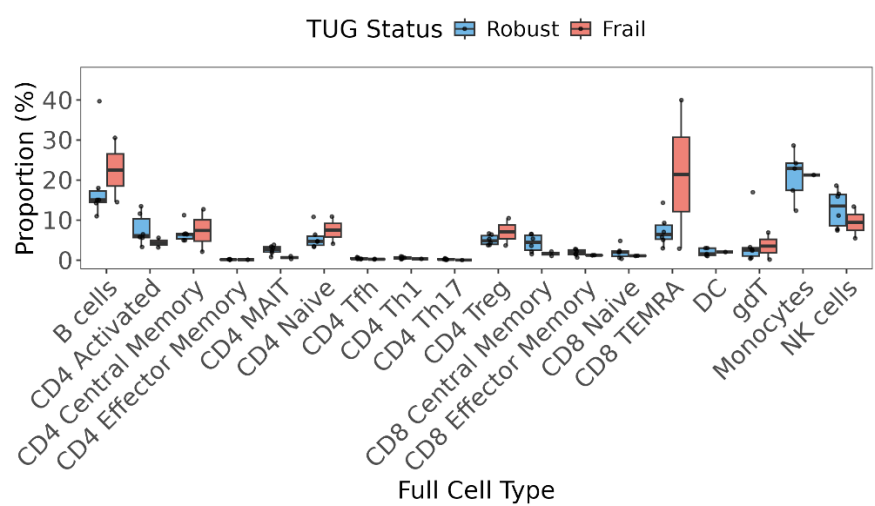

(C)

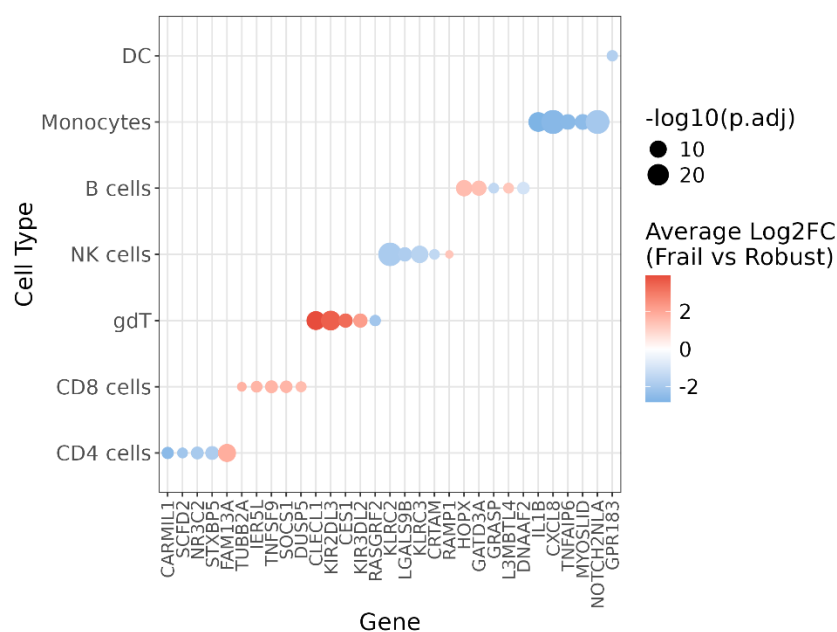

**Figure S3. Single-cell T-cell composition and differential expression by frailty status (A)**

UMAP of T-cell states stratified by frail versus robust individuals. (B) Cell-type proportions at baseline. (C) Top unique differentially expressed genes in monocytes, CD8 T cells, CD4 T cells, and $\gamma\delta$ T cells in by frail versus robust individuals. Cluster marker genes were identified using Seurat's FindAllMarkers function with Wilcoxon rank-sum test. P values were adjusted for multiple testing using Bonferroni correction.

(A)

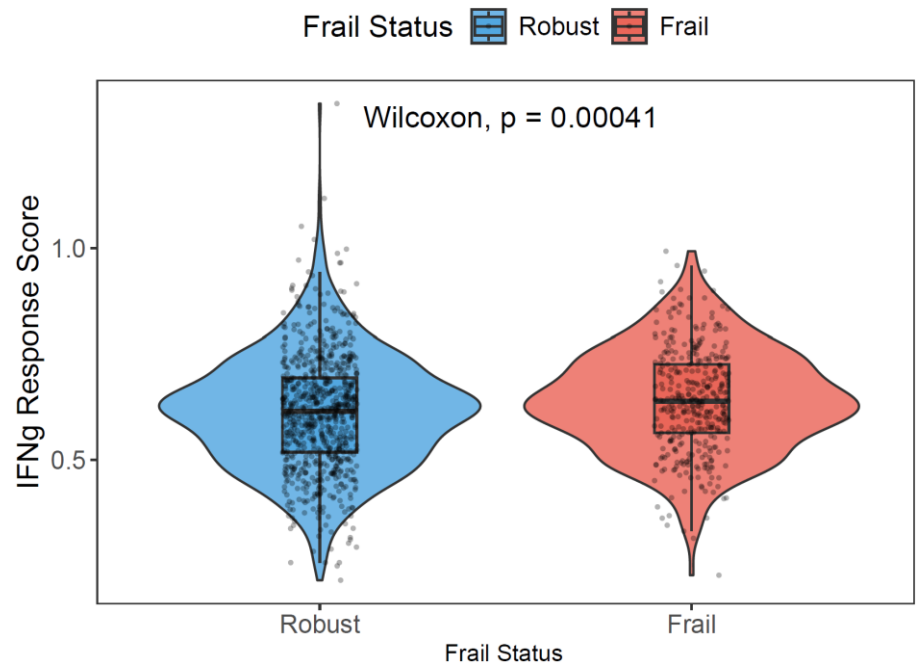

(B)

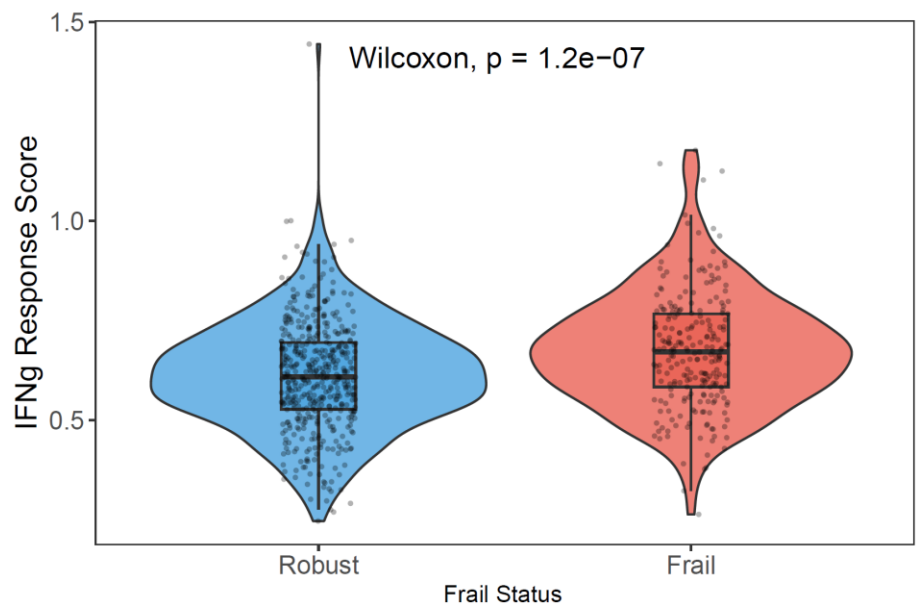

**Figure S4. IFNγ-response activity in (A) CD8 TEMRA and (B) $\gamma\delta$ T-cells by frailty status.** Cell-level distributions and Wilcoxon p values are shown for descriptive visualisation.

(A)

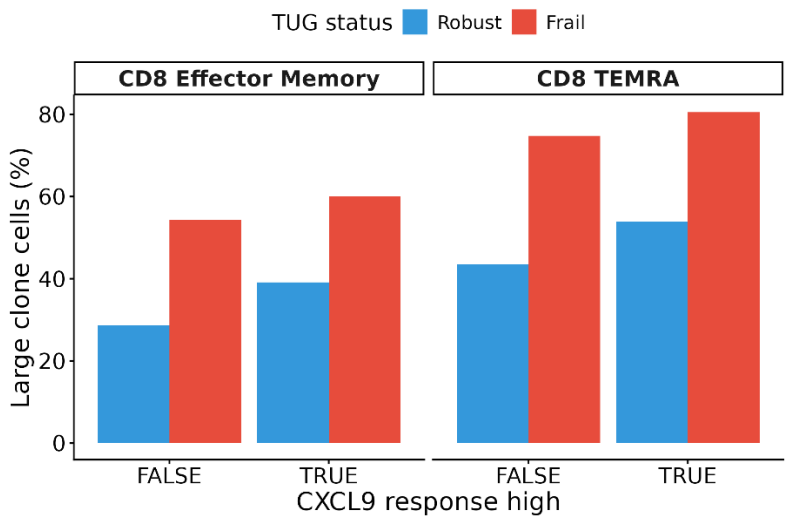

(B)

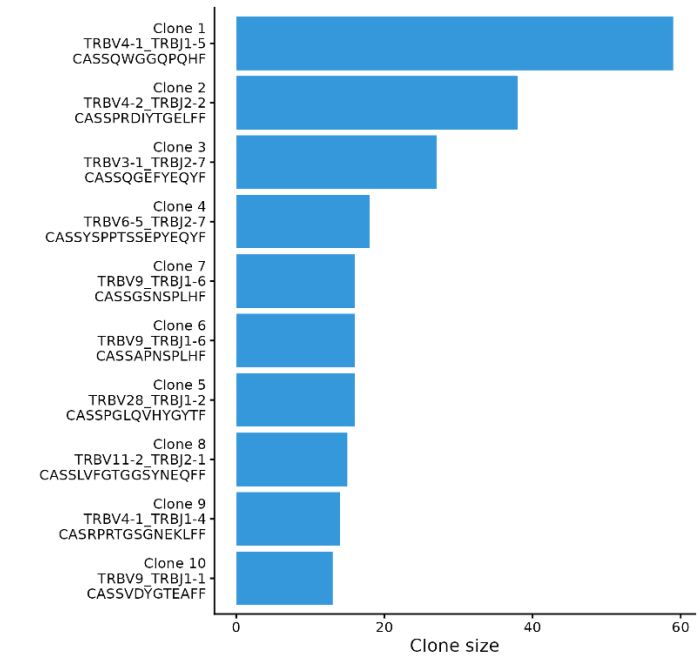

(C)

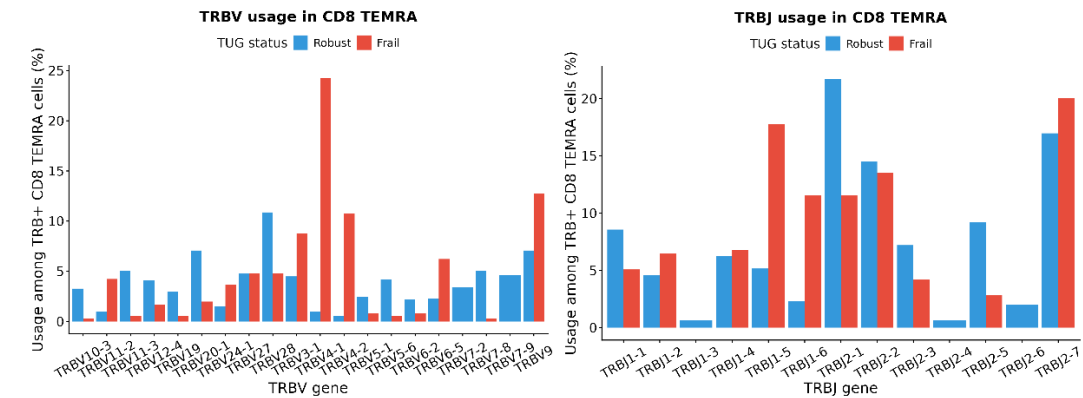

**Figure S5. T-cell receptor clonality in cytotoxic subsets by frailty status.** (A) Large-clone fractions in CD8 effector-memory and CD8 TEMRA cells by CXCL9-response status and frailty group. (B) Top 10 expanded CD8 TEMRA clonotypes. The largest apparent expansions were confined to individual donors. (C) TRBV and TRBJ usage in CD8 TEMRA cells. Analyses are descriptive and do not support a shared antigen-specific repertoire.

(A)

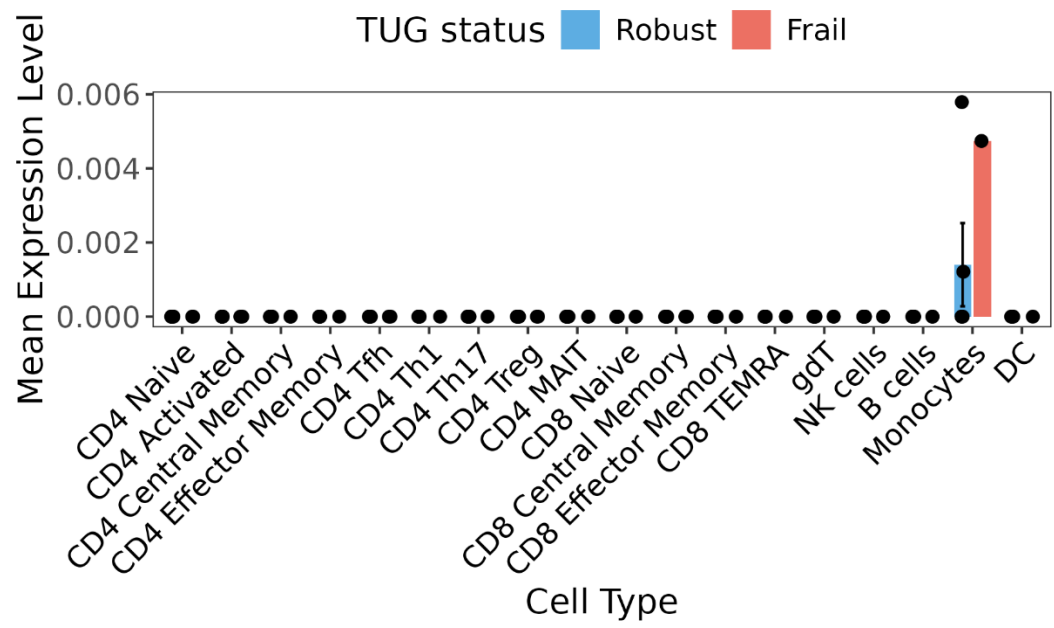

(B)

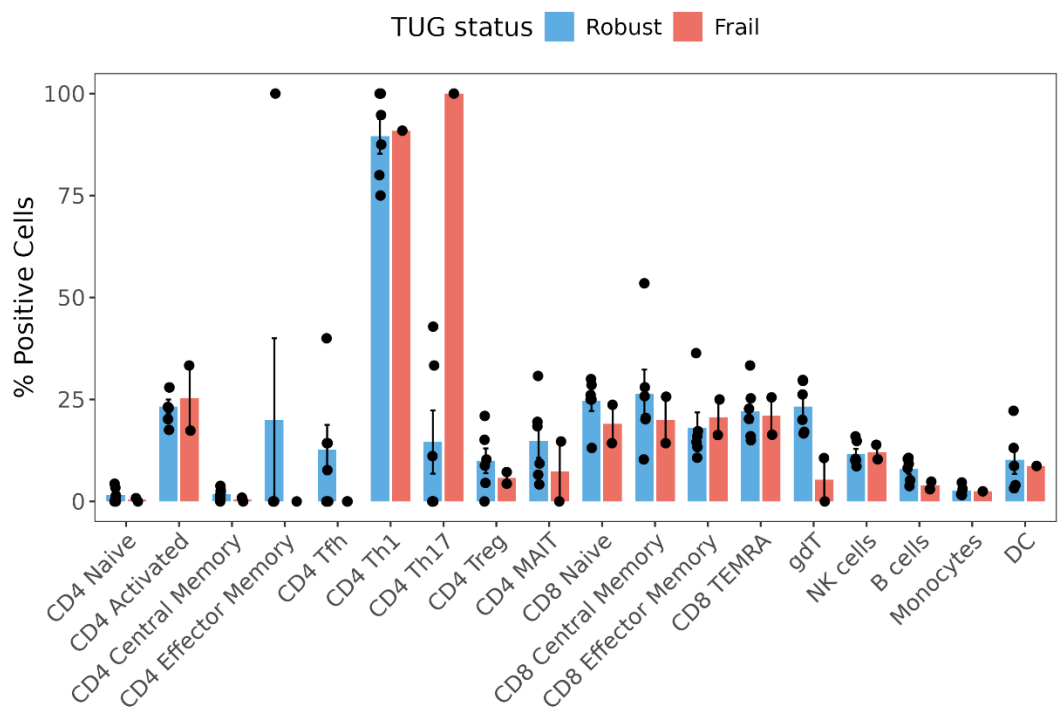

(C)

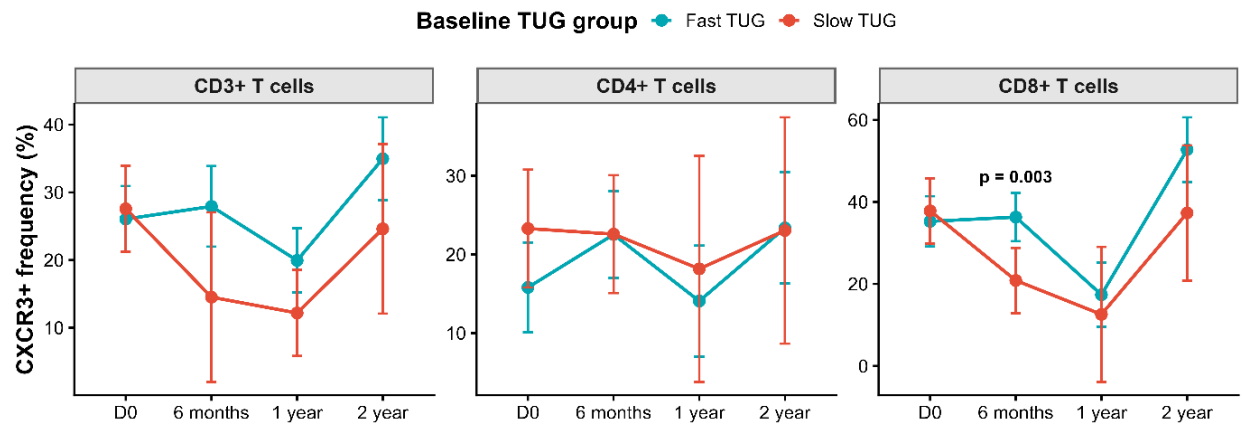

**Figure S6. CXCL9 and CXCR3 immunophenotyping in T-cell subsets.** (A) CXCL9-related signal across immune-cell subsets by frailty status. (B) CXCR3+ frequencies across T-cell subsets. (C) Adjusted longitudinal CXCR3+ frequencies in total CD3, CD4, and CD8 T cells. Baseline CXCR3+ frequencies in total CD3+, CD4+, and CD8+ T cells were not associated with frailty (standardised $\beta = -0.24$, 95% CI $-0.78$ to $0.30$, $p = 0.355$, $q = 0.625$; $\beta = -0.28$, 95% CI $-0.82$ to $0.26$, $p = 0.289$, $q = 0.625$; and $\beta = -0.14$, 95% CI $-0.69$ to $0.42$, $p = 0.601$, $q = 0.763$, respectively). No baseline CXCR3 measure survived correction across the full flow panel (Table S9B).

(A)

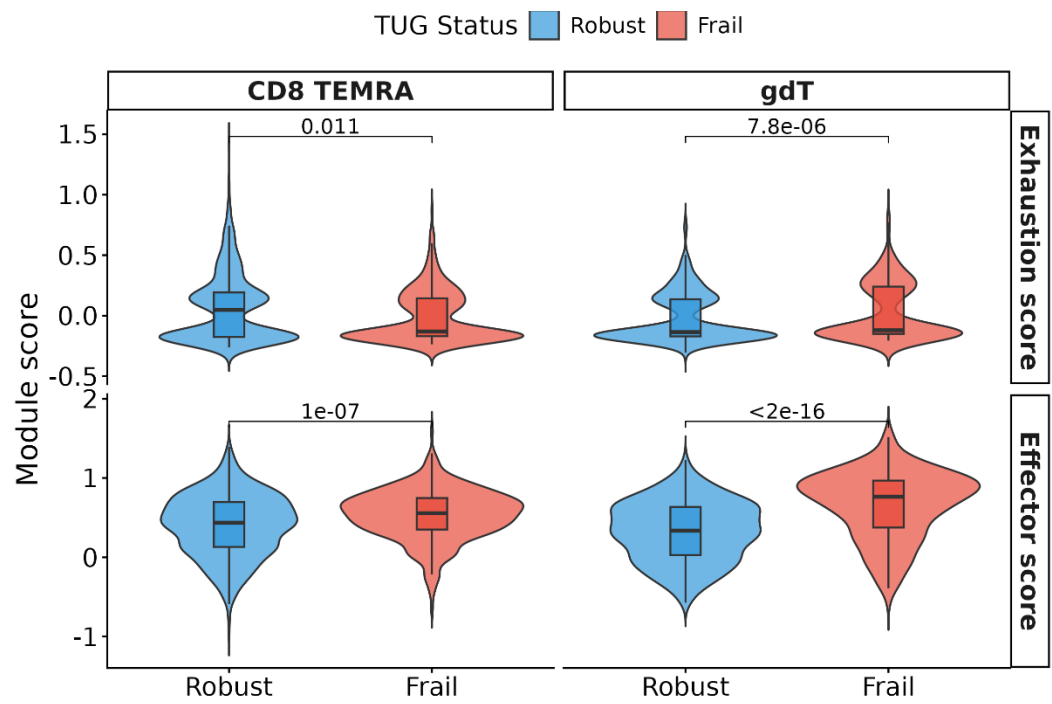

(B)

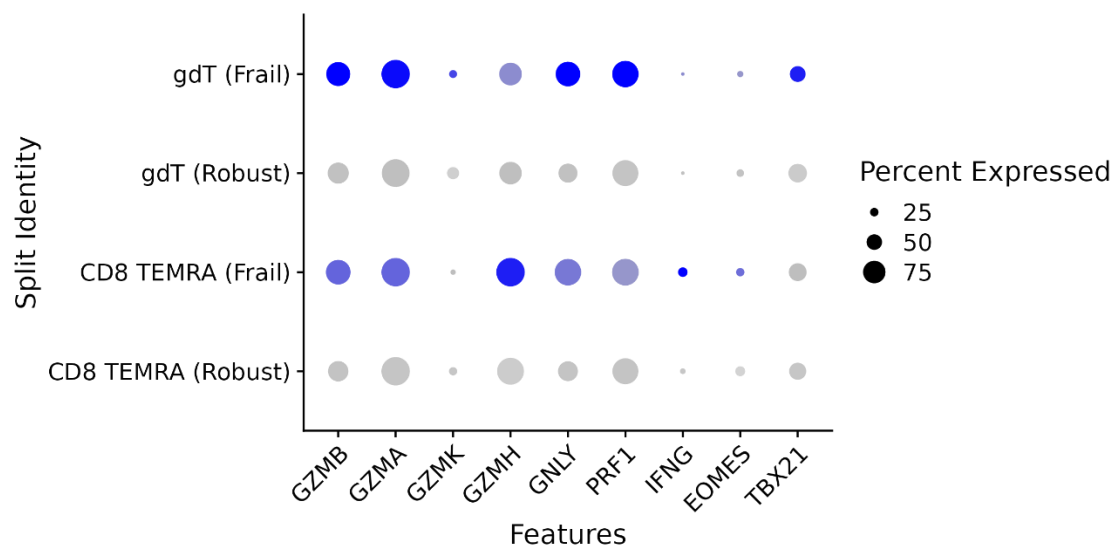

(C)

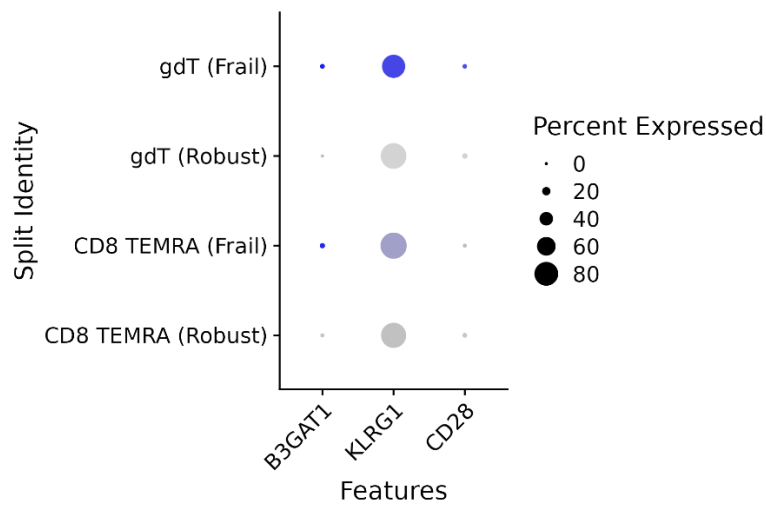

(D)

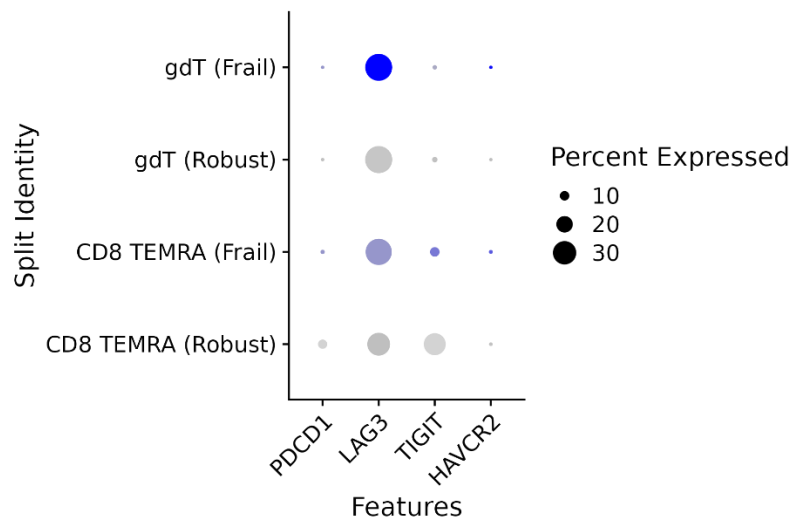

**Figure S7. Cytotoxic-effector, exhaustion, and terminal-differentiation signatures in CD8 TEMRA and $\gamma\delta$ T cells.** Frail donors had higher descriptive (B) cytotoxic-effector and (C) senescence-module scores, with limited expression of the (D) canonical checkpoint genes shown. Module scores were calculated with Seurat AddModuleScore. Cell-level Wilcoxon p values are descriptive and do not establish donor-level significance.

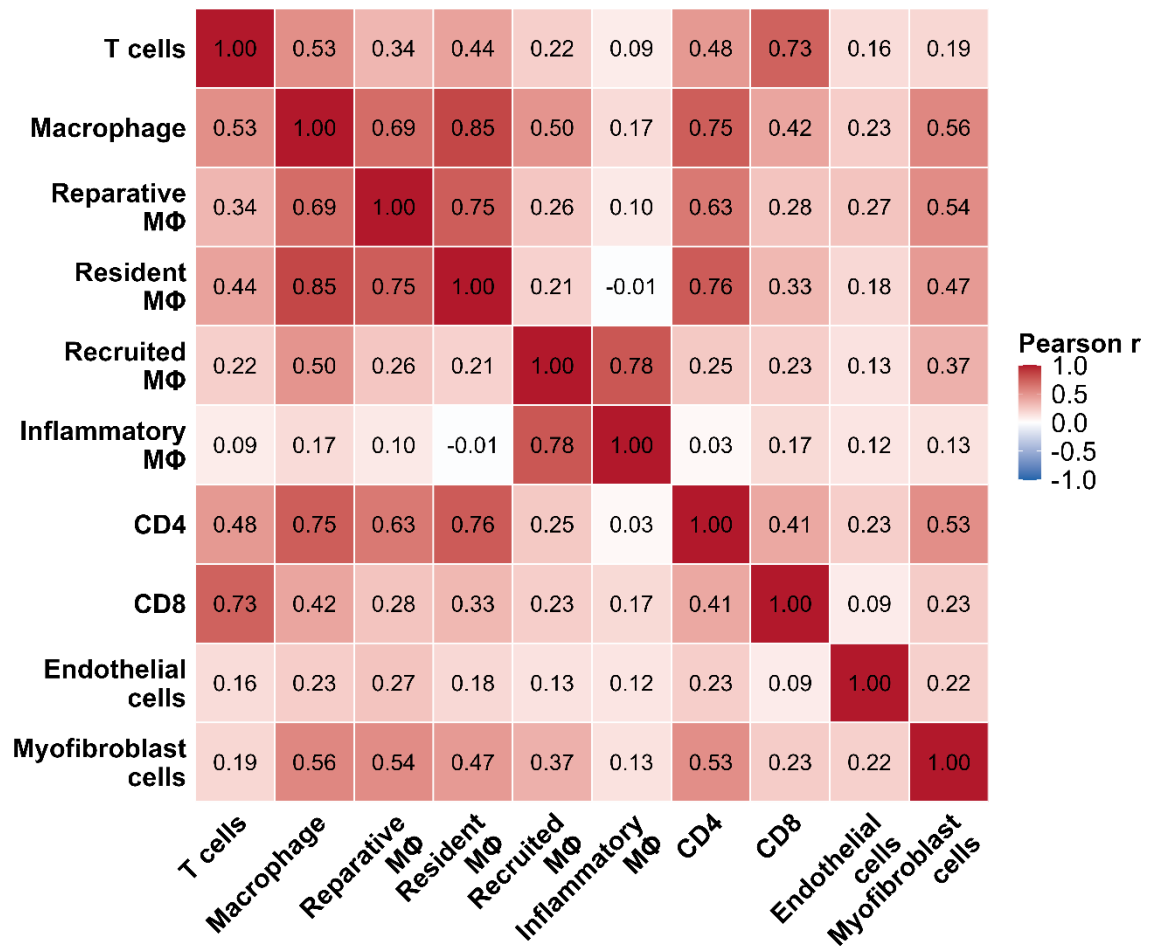

**Figure S8. Pairwise correlation matrix of immune and stromal gene modules.** Tiles show Pearson r values across standardised module scores.

(A)

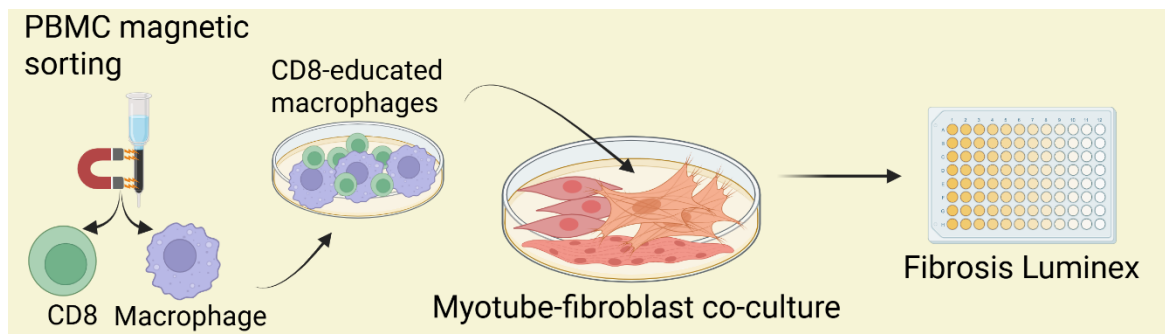

(B)

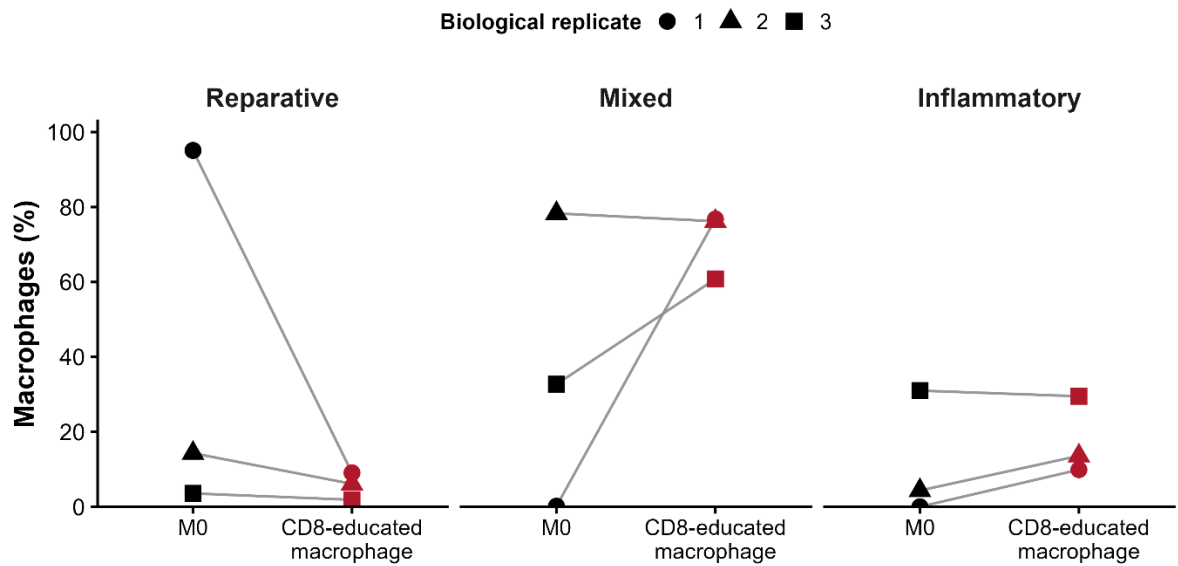

(C)

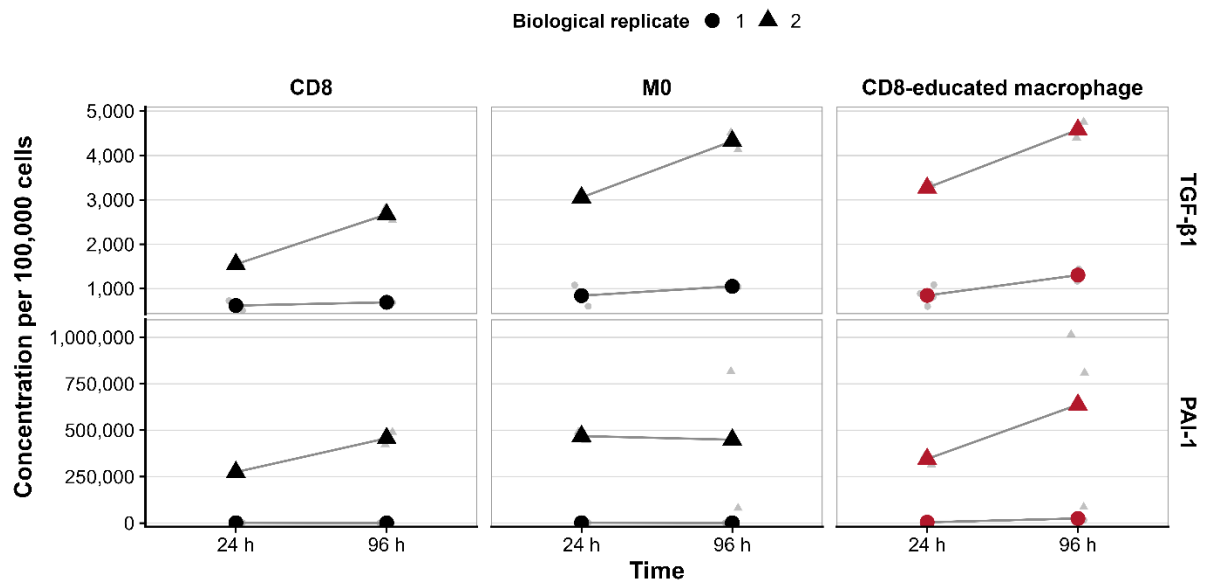

(D)

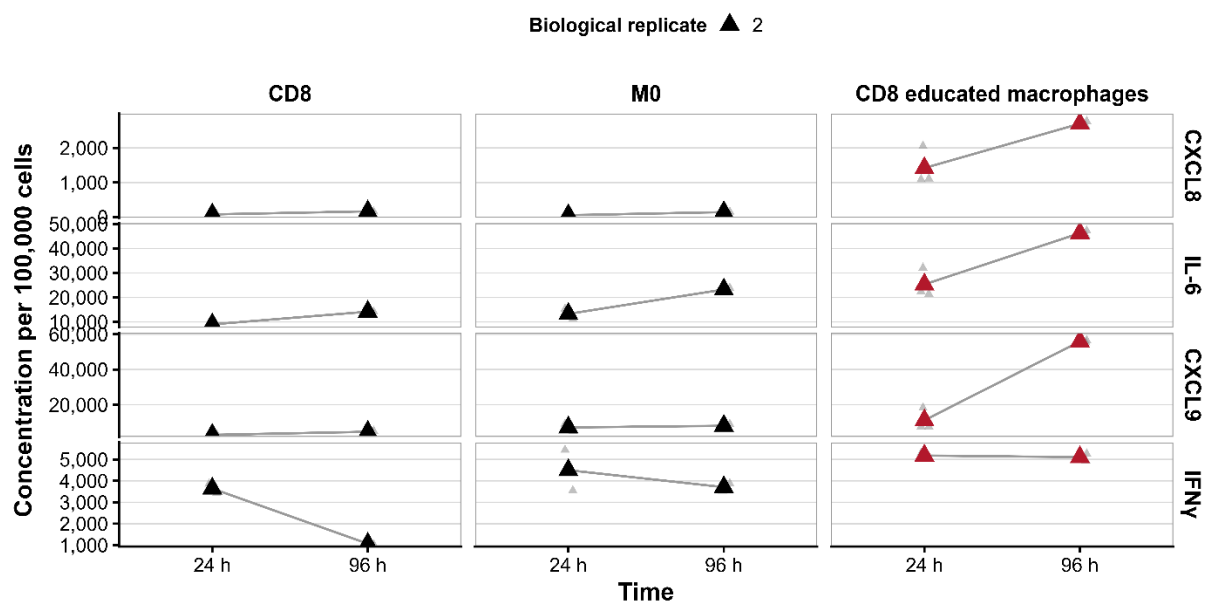

**Figure S9. Immune-stromal co-culture experiments** (A) Experimental schematic. (B) Macrophage phenotype after 48h of CD8 education, from n=3 biological replicates. CD163/CD86 quadrant frequencies among HLA-DR+CD11b+ macrophages (gating, Methods Figure M4). Reparative CD163+ CD86-; mixed CD163+CD86+; inflammatory CD86+ CD163. (C) Secreted total TGF-β1 (pg/mL) and PAI-1 (pg/mL) normalised to cell count at 24h and 96h of immune–stromal co-culture for CD8, M0 and CD8-educated macrophages. Data from n=2 independent PBMC donors and n=2 myotube-fibroblast donor pairs with technical replicates specified in Table S25. Donors are shown individually because of the scale difference. Values are descriptive. (D) Inflammatory and chemokine secretome of the immune–stromal co-culture. CXCL9, IFNγ, IL-6 and CXCL8 concentrations (pg/mL, log scale) across co-culture conditions. Data are from a single biological donor; grey points show individual technical replicates and triangles the technical mean. All values are descriptive. Corresponding numerical values are given in Table S26.

### Supplementary Tables

| Characteristic | Total | Human discovery cohort |  | Human validation cohort |  |
| --- | --- | --- | --- | --- | --- |
|  |  | Robust | Frail | Robust | Frail |
| N | 55 | 18 | 4 | 17 | 16 |
| Age, years* | 40.00<br>(33.00-71.50) | 32.50<br>(27.20-36.00) | 76.00<br>(71.00-81.00) | 37.00<br>(33.00-66.00) | 72.00<br>(64.20-75.20) |
| Older adults, n (%) | 27 (49.10%) | 3 (16.70%) | 4 (100.00%) | 6 (35.30%) | 14 (87.50%) |
| Male, n (%) | 28 (50.90%) | 10 (55.60%) | 2 (50.00%) | 7 (41.20%) | 9 (56.20%) |
| Race |  |  |  |  |  |
| Chinese | 47 (85.50%) | 16 (88.90%) | 4 (100.00%) | 12 (70.60%) | 15 (93.80%) |
| Malay | 2 (3.60%) | 0 (0.00%) | 0 (0.00%) | 2 (11.80%) | 0 (0.00%) |
| Indian | 3 (5.50%) | 1 (5.60%) | 0 (0.00%) | 2 (11.80%) | 0 (0.00%) |
| Caucasian | 2 (3.60%) | 0 (0.00%) | 0 (0.00%) | 1 (5.90%) | 1 (6.20%) |
| Others | 1 (1.80%) | 1 (5.60%) | 0 (0.00%) | 0 (0.00%) | 0 (0.00%) |
| CMV serology positive, n (%) | 43 (78.20%) | 9 (50.00%) | 4 (100.00%) | 16 (94.10%) | 14 (87.50%) |
| Age-adjusted CCI* | 0.00 (0.00-1.00) | 0.00 (0.00-0.00) | 1.50 (1.00-2.20) | 0.00 (0.00-0.00) | 0.50 (0.00-1.00) |
| 3-m TUG, s* | 8.66 (7.06-10.19) | 6.79 (5.94-7.23) | 11.03 (10.86-11.22) | 8.33 (7.65-8.87) | 10.82 (9.96-12.86) |

**Table S1. Demographics of the discovery and validation cohorts.** Continuous variables marked with an asterisk are median (IQR). CCI, Charlson Comorbidity Index; CMV, cytomegalovirus; TUG, timed up-and-go. "Other" race comprises one Boyanese participant.

| Sample ID | Age, years | Sex | CMV serostatus | Physically Frail |
| --- | --- | --- | --- | --- |
| 6 | 28 | Male | Negative | No |
| 9 | 36 | Female | Negative | No |
| 10 | 72 | Male | Positive | Yes |
| 11 | 27 | Female | Positive | No |
| 13 | 23 | Female | Positive | No |
| 14 | 71 | Female | Positive | Yes |
| 15 | 71 | Female | Positive | No |
| 18 | 81 | Female | Positive | Yes |

**Table S2. Participants included in the baseline single-cell RNA-sequencing subcohort.**

Slow TUG was defined as baseline 3-m TUG $\geq 9.45$ s. CMV, cytomegalovirus.

| Module | Adjustment | N | Standardized $\beta$<br>(95% CI) | Adjusted<br>$R^2$ | Adjusted<br>p value |
| --- | --- | --- | --- | --- | --- |
| *Discovery cohort* |  |  |  |  |  |
| IFNγ-CXCR3 | CCI | 21 | 0.53 (0.04-1.03) | 0.16 | 0.04 |
| Inflammaging | CCI | 21 | 0.55 (0.07-1.02) | 0.19 | 0.04 |
| IFNγ-CXCR3 | CCI + CMV | 21 | 0.54 (0.03-1.06) | 0.12 | 0.04 |
| Inflammaging | CCI + CMV | 21 | 0.55 (0.06-1.05) | 0.14 | 0.04 |
| *Validation cohort* |  |  |  |  |  |
| IFNγ-CXCR3 | CCI | 33 | 0.21 (-0.17-0.59) | -0.02 | 0.54 |
| Inflammaging | CCI | 33 | 0.07 (-0.30-0.44) | -0.06 | 0.71 |
| IFNγ-CXCR3 | CCI + CMV | 33 | 0.22 (-0.14-0.57) | 0.09 | 0.45 |
| Inflammaging | CCI + CMV | 33 | 0.08 (-0.28-0.43) | 0.05 | 0.66 |

**Table S3A. Associations of plasma immune modules with age-adjusted TUG residuals in the discovery and validation cohorts.** Primary models adjusted for CCI; secondary models additionally adjusted for CMV serostatus. $\beta$ values are standardised. BH correction was applied within the prespecified module family. One discovery-cohort sample (participant ID 19) was excluded after failing assay quality control.

| Module | Adjustment | N | Standardized $\beta$<br>(95% CI) | Adjusted $R^2$ | Adjusted p value |
| --- | --- | --- | --- | --- | --- |
| *Age-residualised Maximum Handgrip Strength* |  |  |  |  |  |
| IFNγ-CXCR3 | CCI | 21 | 0.02 (-0.54-0.58) | -0.06 | 0.93 |
| Inflammaging | CCI | 21 | 0.35 (-0.16-0.87) | 0.05 | 0.34 |
| IFNγ-CXCR3 | CCI + CMV | 21 | 0.08 (-0.45-0.62) | 0.06 | 0.74 |
| Inflammaging | CCI + CMV | 21 | 0.31 (-0.18-0.81) | 0.14 | 0.40 |
| *Age-residualised Sit-to-stand Test* |  |  |  |  |  |
| IFNγ-CXCR3 | CCI | 20 | -0.24 (-0.72-0.23) | 0.05 | 0.35 |
| Inflammaging | CCI | 20 | -0.22 (-0.69-0.26) | 0.04 | 0.35 |
| IFNγ-CXCR3 | CCI + CMV | 20 | -0.20 (-0.69-0.30) | 0.04 | 0.41 |
| Inflammaging | CCI + CMV | 20 | -0.24 (-0.72-0.23) | 0.07 | 0.41 |

**Table S3B. Sensitivity analyses with maximum handgrip strength and sit-to-stand test.**

Similar analyses to S3A, in discovery cohort.

| Module | Analysis | Pooled standardized $\beta$ (95% CI) | p value | BH q value | Q | I<sup>2</sup> |
| --- | --- | --- | --- | --- | --- | --- |
| IFNγ-CXCR3 | CCI-adjusted | 0.33 (0.05-0.62) | 0.02 | 0.04 | 1.18 | 15.50% |
| IFNγ-CXCR3 | CCI + CMV-adjusted | 0.33 (0.05-0.61) | 0.02 | 0.04 | 1.19 | 15.60% |
| Inflammaging | CCI-adjusted | 0.25 (-0.02-0.53) | 0.07 | 0.07 | 2.68 | 62.70% |
| Inflammaging | CCI + CMV-adjusted | 0.24 (-0.03-0.52) | 0.08 | 0.08 | 2.63 | 62.00% |

**Table S4. Cross-cohort fixed-effect meta-analysis of plasma immune-module associations with age-adjusted TUG residuals.** Primary models adjusted for CCI; secondary models additionally adjusted for CMV serostatus (n = 54, one discovery-cohort sample (participant ID 19) was excluded after failing assay quality control). Pooled estimates indicate cross-cohort consistency rather than definitive validation. Q and I<sup>2</sup> quantify between-cohort heterogeneity. BH, Benjamini–Hochberg.

| Predictor module | Bootstrap mean $\beta$ (SD) | BCa 95%CI | Two-sided bootstrap p value | Directional consistency |
| --- | --- | --- | --- | --- |
| IFNγ-CXCR3 | 0.46 (0.22) | 0.07–0.93 | 0.01 | 99.40% |
| Inflammaging | 0.26 (0.14) | 0.00–0.56 | 0.06 | 96.80% |

**Table S5. Bootstrap sensitivity analyses for plasma immune-module associations with age-adjusted TUG residuals.** Results are based on 5,000 bootstrap resamples of models adjusted for CCI and CMV serostatus (n = 54, one discovery-cohort sample (participant ID 19) was excluded after failing assay quality control). Standardised $\beta$ represents the SD difference in residualised TUG per 1-SD higher module score. BCa, bias-corrected and accelerated.

| Diagnostic | Result |
| --- | --- |
| Correlation between IFNγ-CXCR3 and inflammaging modules | r = 0.72 |
| VIF: IFNγ-CXCR3 module | 2.33 |
| VIF: inflammaging module | 2.08 |
| VIF: CCI | 1.12 |
| VIF: CMV | 1.24 |
| **IFNγ-CXCR3 + CCI + CMV model** |  |
| Largest Cook's distance | 0.13 |
| Most influential observations | ID 52, 47, 38 |
| Highest leverage observation | Hat = 0.24, ID 49 |
| Other high-leverage observations | Hat = 0.21, ID 38; Hat = 0.15, ID 18; Hat = 0.14, ID 15 |
| Largest absolute DFBETA | \|DFBETA\| = 0.65, ID 38 |
| LOO | $\beta$ range 0.23 to 0.42, median 0.32; p value range 0.01 to 0.12 |
| **inflammaging + CCI + CMV model** |  |
| Largest Cook's distance | 0.13 |
| Most influential observations | Rows 52, 47, 20, 16, 10 |
| Highest leverage observation | Hat = 0.22, ID 49 |
| Other high-leverage observations | Hat = 0.20, ID 18; Hat = 0.17, ID 15; Hat = 0.15, ID 13 |
| Largest absolute DFBETA | \|DFBETA\| = 0.41, ID 16 |
| LOO | $\beta$ range 0.18 to 0.29, median 0.23; p value range 0.05 to 0.20 |

**Table S6. Collinearity and influence diagnostics for plasma immune-module models.**

VIF, variance-inflation factor; LOO, leave-one-out. Models adjusted for CCI and CMV serostatus.

| Rank | Marker | Pooled<br>standardized $\beta$<br>(95% CI) | p value | BH q value | I<sup>2</sup> |
| --- | --- | --- | --- | --- | --- |
| 1 | CXCL9 | 0.33 (0.02-0.63) | 0.04 | 0.09 | 61.10% |
| 2 | IFNγ | 0.27 (-0.01-0.55) | 0.06 | 0.09 | 0.00% |
| 3 | IL18R1 | 0.19 (-0.09-0.47) | 0.18 | 0.18 | 0.00% |

**Table S7. Representative-marker prioritisation within the IFNγ–CXCR3 plasma module.**

Markers were ranked by inverse variance fixed-effect pooling of cohort-specific standardised associations with age-adjusted TUG residuals. $I^2$ quantifies between-cohort heterogeneity.

| Analysis | Discovery $\beta$ | Discovery p value | Validation $\beta$ | Validation p value | Pooled $\beta$ (95% CI) | Pooled p value | BH q value |
| --- | --- | --- | --- | --- | --- | --- | --- |
| CCI-adjusted | 0.71 | 0.02 | 0.16 | 0.40 | 0.33 (0.02 to 0.63) | 0.04 | 0.09 |
| CCI + CMV-adjusted | 0.74 | 0.02 | 0.13 | 0.48 | 0.29 (-0.01 to 0.59) | 0.06 | 0.09 |

**Table S8. Cohort-specific and pooled associations of plasma CXCL9 with age-adjusted TUG residuals.** Primary models adjusted for CCI; secondary models additionally adjusted for CMV serostatus. Pooled estimates were obtained by fixed-effect meta-analysis.

| Measure (%) | N | Standardised $\beta$ (95% CI) | p value | BH q value |
| --- | --- | --- | --- | --- |
| CD8 TEMRA | 19 | 0.55 (0.05 to 1.05) | 0.03 | 0.08 |
| PD-1+ CD8 TEMRA | 19 | -0.32 (-0.84 to 0.20) | 0.21 | 0.35 |
| CD56+ CD8 TEMRA | 19 | -0.38 (-1.16 to 0.40) | 0.32 | 0.63 |
| CD28- CD8 TEMRA | 19 | 0.18 (-0.39 to 0.74) | 0.52 | 0.52 |
| PD-1+ CD4- CD8- T cells | 19 | 0.17 (-0.37 to 0.71) | 0.51 | 0.52 |
| CD28- CD4- CD8- T cells | 19 | 0.57 (0.12 to 1.02) | 0.02 | 0.08 |

**Table S9A. Adjusted associations of peripheral flow-cytometry phenotypes with age-adjusted TUG residuals.** Each model regressed the age-adjusted TUG residual z score on a 1-SD higher flow-cytometry measure with adjustment for CCI and CMV serostatus. $\beta$ values

are standardised. BH q values correct across all 36 flow measures tested in Tables S9A and S9B.

| **CXCR3 measure** | **N** | **Standardised $\beta$ (95% CI)** | **p value** | **BH q value** |
| --- | --- | --- | --- | --- |
| CXCR3+ %, Total CD3 | 19 | -0.24 (-0.78 to 0.30) | 0.36 | 0.63 |
| CXCR3+ %, Total CD4 | 19 | -0.28 (-0.82 to 0.26) | 0.29 | 0.63 |
| CXCR3+ %, Total CD8 | 19 | -0.14 (-0.69 to 0.42) | 0.60 | 0.76 |
| CXCR3 gMFI, Tfh | 19 | -0.21 (-0.75 to 0.34) | 0.43 | 0.68 |
| CXCR3+ %, Tfh | 19 | -0.05 (-0.65 to 0.55) | 0.86 | 0.97 |
| CXCR3 gMFI, Th1 | 19 | -0.15 (-0.70 to 0.41) | 0.58 | 0.76 |
| CXCR3 gMFI, Th17 | 19 | -0.28 (-0.81 to 0.26) | 0.29 | 0.63 |
| CXCR3 gMFI, Treg | 19 | -0.08 (-0.66 to 0.49) | 0.76 | 0.91 |
| CXCR3+ %, Treg | 19 | -0.04 (-0.61 to 0.54) | 0.89 | 0.97 |
| CXCR3 gMFI, CD4- CD8- (DN) | 19 | -0.38 (-0.99 to 0.22) | 0.20 | 0.63 |
| CXCR3+ %, CD4- CD8- (DN) | 19 | -0.28 (-0.81 to 0.25) | 0.28 | 0.63 |
| CXCR3 gMFI, CD28- CD4- CD8- | 19 | -0.26 (-0.80 to 0.28) | 0.32 | 0.63 |
| CXCR3 gMFI, PD-1+ CD4- CD8- | 19 | -0.32 (-0.84 to 0.21) | 0.21 | 0.63 |
| CXCR3 gMFI, CD8 TEMRA | 19 | -0.29 (-0.84 to 0.27) | 0.29 | 0.63 |
| CXCR3+ %, CD8 TEMRA | 19 | 0.00 (-0.54 to 0.55) | 0.99 | 0.99 |
| CXCR3 gMFI, CD28- CD8 TEMRA | 19 | -0.27 (-0.86 to 0.32) | 0.35 | 0.63 |
| CXCR3+ %, CD28- CD8 TEMRA | 19 | -0.05 (-0.60 to 0.50) | 0.85 | 0.97 |
| CXCR3 gMFI, PD-1+ CD8 TEMRA | 19 | -0.37 (-0.88 to 0.14) | 0.15 | 0.63 |
| CXCR3+ %, PD-1+ CD8 TEMRA | 19 | 0.00 (-0.56 to 0.57) | 0.99 | 0.99 |
| CXCR3 gMFI, KLRG1+CD57+ CD8 | 19 | -0.24 (-0.77 to 0.29) | 0.34 | 0.63 |
| CXCR3+ %, KLRG1+CD57+ CD8 | 19 | -0.15 (-0.77 to 0.47) | 0.62 | 0.76 |
| CXCR3 gMFI, NK | 19 | 0.25 (-0.34 to 0.85) | 0.38 | 0.63 |

| CXCR3 measure | N | Standardised $\beta$ (95% CI) | p value | BH q value |
| --- | --- | --- | --- | --- |
| CXCR3+ %, NK | 19 | 0.55 (-0.02 to 1.12) | 0.06 | 0.63 |
| CXCR3 gMFI, CD56+ CD8 TEMRA | 18 | -0.42 (-1.03 to 0.18) | 0.15 | 0.63 |
| CXCR3+ %, CD56+ CD8 TEMRA | 18 | -0.02 (-0.60 to 0.56) | 0.95 | 0.99 |
| CXCR3 gMFI, PD-1+ CD8 | 19 | -0.23 (-0.76 to 0.30) | 0.37 | 0.63 |
| CXCR3+ %, PD-1+ CD8 | 19 | -0.19 (-0.77 to 0.38) | 0.49 | 0.72 |

**Table S9B. Adjusted associations of peripheral flow-cytometry CXCR3 measures with age-adjusted TUG residuals.** Each model regressed the age-adjusted TUG residual z score on a 1-SD higher CXCR3 measure with adjustment for CCI and CMV serostatus. $\beta$ values are standardised. BH q values correct across all 36 flow measures tested in Tables S9A and S9B. No CXCR3 association survived correction across the full flow panel.

| Characteristic | Whole cohort (N=340) | Sarcopenic (N=102) | Non-sarcopenic (N=238) |
| --- | --- | --- | --- |
| Age, median (IQR) | 66.00 (57.00-72.00) | 71.00 (66.00-76.80) | 63.00 (55.20-70.00) |
| Older adults, n (%) | 186 (54.70%) | 79 (77.50%) | 107 (45.00%) |
| Male, n (%) | 199 (58.50%) | 66 (64.70%) | 133 (55.90%) |
| CXCL9 expression, median (IQR) | 6.22 (5.92-6.59) | 6.21 (5.91-6.54) | 6.23 (5.92-6.64) |

**Table S10. Demographic characteristics of the muscle transcriptomic cohort.** Values are median (IQR) or n (%). Sarcopenia was classified using Asian Working Group for Sarcopenia 2019 criteria.

| Module | Genes, n | Genes included | Cronbach $\alpha$ | Mean inter-item r | PC1 variance, % |
| --- | --- | --- | --- | --- | --- |
| Leukocyte infiltration | 1 | PTPRC | N/A | N/A | N/A |
| T-cell lineage | 5 | CD3E, TRAC, TRBC2, CD2, CD247 | 0.81 | 0.45 | 56.70 |
| CD4 | 1 | CD4 | N/A | N/A | N/A |
| CD8 | 1 | CD8A | N/A | N/A | N/A |
| Endothelial | 3 | PECAM1, VWF, CDH5 | 0.83 | 0.62 | 74.40 |
| Myofibroblast | 4 | POSTN, LOX, TNC, FAP | 0.74 | 0.54 | 66.00 |
| Macrophage | 9 | CD14, ITGAM, CSF1R, LST1, AIF1, TYROBP, FCER1G, SPI1, CD68 | 0.90 | 0.55 | 60.90 |
| Inflammatory macrophage | 7 | IL1B, TNF, IL6, NLRP3, CXCL2, CXCL3, PTGS2 | 0.89 | 0.62 | 68.30 |
| M2 macrophage | 6 | MSR1, MERTK, CCL18, MAF, CD163, MRC1 | 0.76 | 0.34 | 51.40 |
| Resident macrophage | 7 | FOLR2, LYVE1, STAB1, SELENOP, C1QA, C1QB, C1QC | 0.91 | 0.58 | 70.00 |
| Recruited macrophage | 7 | CCL2, CCL4, CCL8, FCN1, VCAN, S100A8, S100A9 | 0.82 | 0.44 | 53.60 |

**Table S11. Immune and stromal gene-module definitions and internal-consistency metrics.** Cronbach $\alpha$ and mean inter-item $r$ are not applicable to single-gene signatures. PC1 variance is the percentage of module variance explained by the first principal component.

| Module 1 | Module 2 | Pearson $r$ (95% CI) | p value | BH q value |
| --- | --- | --- | --- | --- |
| Macrophage | Resident macrophage | 0.85 (0.82-0.88) | <0.001 | <0.001 |
| Recruited macrophage | Inflammatory macrophage | 0.78 (0.74-0.82) | <0.001 | <0.001 |
| Resident macrophage | CD4 | 0.76 (0.71-0.80) | <0.001 | <0.001 |
| M2 macrophage | Resident macrophage | 0.75 (0.70-0.79) | <0.001 | <0.001 |
| Macrophage | CD4 | 0.75 (0.70-0.79) | <0.001 | <0.001 |
| T cells | CD8 | 0.73 (0.68-0.78) | <0.001 | <0.001 |
| Macrophage | M2 macrophage | 0.69 (0.63-0.74) | <0.001 | <0.001 |
| M2 macrophage | CD4 | 0.63 (0.56-0.69) | <0.001 | <0.001 |
| Macrophage | Myofibroblast cells | 0.56 (0.48-0.63) | <0.001 | <0.001 |
| M2 macrophage | Myofibroblast cells | 0.54 (0.46-0.61) | <0.001 | <0.001 |

**Table S12A. Top 10 pairwise Pearson correlations among immune and stromal modules.** Correlations used complete cases across the muscle cohort. BH q values account for all pairwise module comparisons.

| Module 1 | Module 2 | Partial r | p value | BH q value |
| --- | --- | --- | --- | --- |
| Recruited macrophage | Inflammatory macrophage | 0.81 | <0.001 | <0.001 |
| T cells | CD8 | 0.66 | <0.001 | <0.001 |
| Macrophage | Resident macrophage | 0.65 | <0.001 | <0.001 |
| Macrophage | Recruited macrophage | 0.57 | <0.001 | <0.001 |
| M2 macrophage | Resident macrophage | 0.40 | <0.001 | <0.001 |
| Macrophage | Inflammatory macrophage | -0.32 | <0.001 | <0.001 |
| Resident macrophage | Recruited macrophage | -0.32 | <0.001 | <0.001 |
| Resident macrophage | CD4 | 0.28 | <0.001 | <0.001 |
| M2 macrophage | Myofibroblast cells | 0.26 | <0.001 | <0.001 |
| T cells | Myofibroblast cells | -0.22 | <0.001 | <0.001 |

**Table S12B. Top 10 partial correlations among immune and stromal modules.** Each partial correlation controls for the remaining modules in the diagnostic set. BH q values account for all pairwise partial-correlation tests.

| Module | VIF |
| --- | --- |
| Macrophage | 7.71 |
| Resident macrophage | 6.21 |
| Recruited macrophage | 5.00 |
| Inflammatory macrophage | 3.39 |
| CD4 | 2.92 |
| M2 macrophage | 2.68 |
| T cells | 2.66 |
| CD8 | 2.32 |
| Myofibroblast cells | 1.80 |

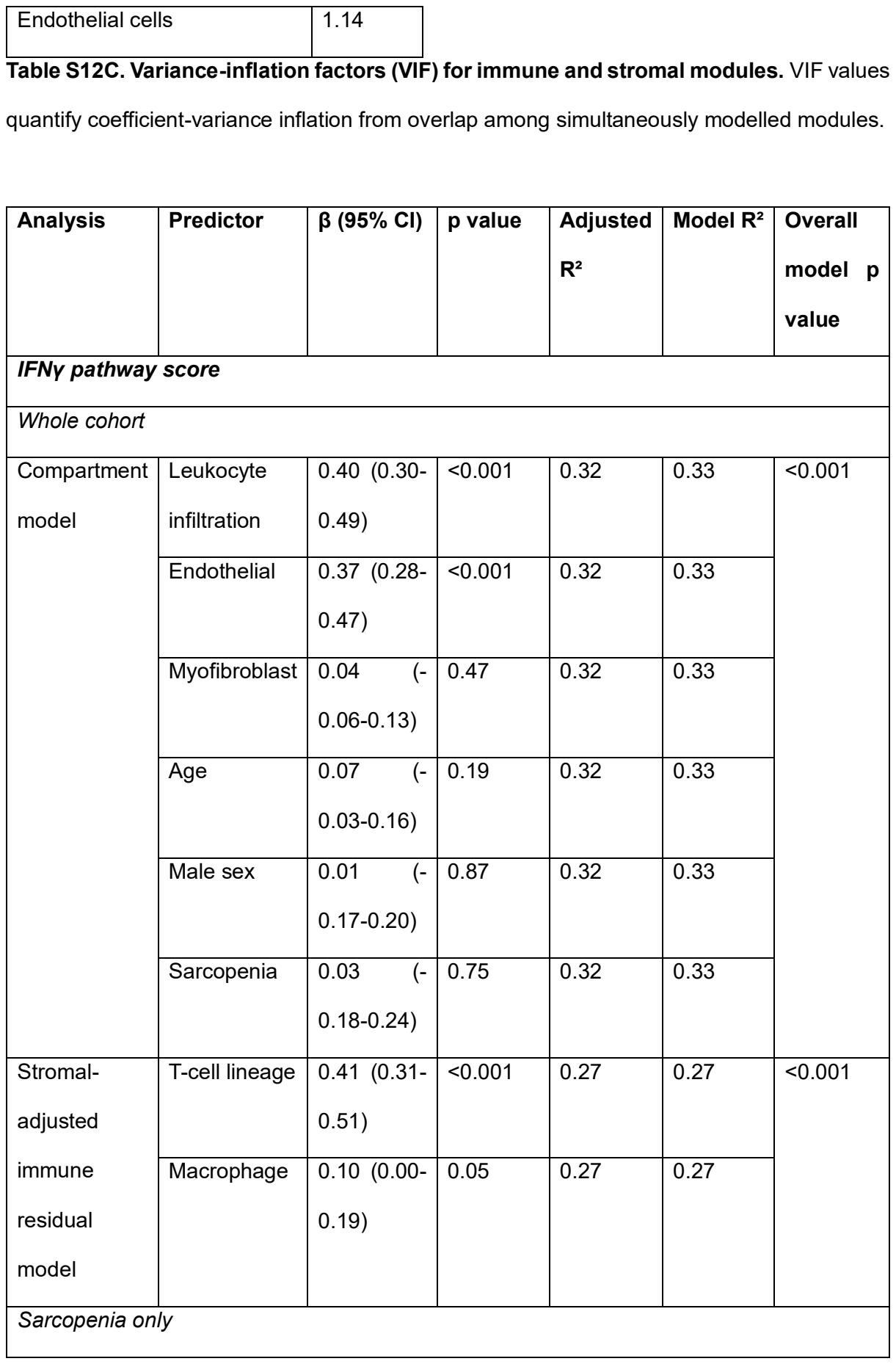

|  |  |  |  |  |  |  |
| --- | --- | --- | --- | --- | --- | --- |
| Compartment model | Leukocyte infiltration | 0.50 (0.34-0.66) | <0.001 | 0.41 | 0.44 | <0.001 |
|  | Endothelial | 0.34 (0.19-0.49) | <0.001 | 0.41 | 0.44 |  |
|  | Myofibroblast | 0.15 (-0.01-0.32) | 0.06 | 0.41 | 0.44 |  |
|  | Age | 0.03 (-0.13-0.19) | 0.70 | 0.41 | 0.44 |  |
|  | Male sex | 0.10 (-0.22-0.43) | 0.52 | 0.41 | 0.44 |  |
| Stromal-adjusted immune residual model | T-cell lineage | 0.64 (0.47-0.80) | <0.001 | 0.44 | 0.45 | <0.001 |
|  | Macrophage | -0.08 (-0.24-0.09) | 0.36 | 0.44 | 0.45 |  |
| Hallmark IFNγ ssGSEA score |  |  |  |  |  |  |
| Whole cohort |  |  |  |  |  |  |
| Compartment model | Leukocyte infiltration | 0.40 (0.31-0.49) | <0.001 | 0.38 | 0.39 | <0.001 |
|  | Endothelial | 0.34 (0.25-0.43) | <0.001 | 0.38 | 0.39 |  |
|  | Myofibroblast | 0.17 (0.08-0.26) | <0.001 | 0.38 | 0.39 |  |
|  | Age | 0.07 (-0.02-0.17) | 0.12 | 0.38 | 0.39 |  |
|  | Male sex | 0.07 (-0.11-0.24) | 0.44 | 0.38 | 0.39 |  |
|  | Sarcopenia | 0.07 (-0.13-0.27) | 0.50 | 0.38 | 0.39 |  |
| Stromal-adjusted immune residual model | T-cell lineage | 0.22 (0.12-0.32) | <0.001 | 0.13 | 0.14 | <0.001 |
|  | Macrophage | 0.14 (0.04-0.24) | 0.006 | 0.13 | 0.14 |  |
| Sarcopenia only |  |  |  |  |  |  |
| Compartment model | Leukocyte infiltration | 0.55 (0.41-0.69) | <0.001 | 0.57 | 0.59 | <0.001 |
|  | Endothelial | 0.30 (0.17-0.43) | <0.001 | 0.57 | 0.59 |  |
|  | Myofibroblast | 0.27 (0.13-0.41) | <0.001 | 0.57 | 0.59 |  |
|  | Age | -0.08 (-0.21-0.06) | 0.25 | 0.57 | 0.59 |  |
|  | Male sex | 0.26 (-0.02-0.54) | 0.07 | 0.57 | 0.59 |  |
| Stromal-adjusted immune residual model | T-cell lineage | 0.50 (0.33-0.66) | <0.001 | 0.36 | 0.37 | <0.001 |
|  | Macrophage | 0.01 (-0.16-0.17) | 0.91 | 0.36 | 0.37 |  |

**Table S13. Determinants of IFNγ-response activity in muscle.** Analyses used the custom IFNγ-response score and Hallmark IFNγ-response ssGSEA sensitivity score in the whole cohort (n = 340) and sarcopenic subgroup (n = 102). Whole-cohort models adjusted for age, sex, and sarcopenia status; subgroup models adjusted for age and sex. Stromal-adjusted immune models evaluated T-cell and macrophage contributions after accounting for endothelial and myofibroblast signatures.

| Model | Predictor | Standardized $\beta$ (95% CI) | Partial r | Partial R<sup>2</sup> | Unique $\Delta R^2$ | p value | BH q value |
| --- | --- | --- | --- | --- | --- | --- | --- |
| Broad-lineage model | T-cell lineage | 0.40 (0.30-0.49) | 0.40 | 0.16 | 0.11 | <0.001 | <0.001 |
|  | Macrophage lineage | 0.23 (0.11-0.34) | 0.21 | 0.04 | 0.03 | <0.001 | <0.001 |
| T-cell subset model | CD4 expression | -0.18 (-0.31--0.06) | -0.15 | 0.02 | 0.01 | 0.005 | 0.005 |
|  | CD8A expression | 0.37 (0.28-0.46) | 0.41 | 0.17 | 0.11 | <0.001 | <0.001 |
|  | Macrophage lineage | 0.43 (0.30-0.56) | 0.34 | 0.12 | 0.07 | <0.001 | <0.001 |
| CD4-only model | CD4 expression | -0.10 (-0.23-0.04) | -0.08 | 0.01 | 0.00 | 0.16 | 0.16 |
| CD8-only model | CD8A expression | 0.35 (0.26-0.44) | 0.39 | 0.15 | 0.10 | <0.001 | <0.001 |

**Table S14A. Predictor-level associations with standardised muscle IFNγ-response activity.** The broad-lineage model included endothelial and myofibroblast modules, age, sex, total T-cell lineage, and macrophage lineage (n = 340). The exploratory subset model replaced the total T-cell module with CD4 and CD8A expression.

| Model | R<sup>2</sup> | Adjusted R<sup>2</sup> | ΔR<sup>2</sup> vs base | AIC | BIC |
| --- | --- | --- | --- | --- | --- |
| Base model | 0.19 | 0.18 | — | 902.76 | 925.74 |
| Broad-lineage model | 0.45 | 0.44 | 0.25 | 778.06 | 808.69 |
| T-cell subset model | 0.45 | 0.44 | 0.26 | 776.37 | 810.83 |
| CD4-only model | 0.34 | 0.33 | 0.15 | 836.62 | 867.26 |
| CD8-only model | 0.44 | 0.43 | 0.25 | 782.59 | 813.22 |

**Table S14B. Overall fit of models explaining standardised muscle IFNγ-response activity.** ΔR<sup>2</sup> is relative to the base model comprising endothelial and myofibroblast modules, age, and sex. AIC, Akaike information criterion; BIC, Bayesian information criterion.

| Terms added | Reduced model | Added df | F | p value |
| --- | --- | --- | --- | --- |
| CD4 and CD8A expression | Base covariates + macrophage lineage | 2 | 34.53 | <0.001 |
| Macrophage lineage | Base covariates + CD4 + CD8A | 1 | 43.08 | <0.001 |

**Table S14C. Nested tests of incremental T-cell-subset and macrophage contributions to standardised muscle IFNγ-response activity.** Subset analyses are exploratory because CD4 and CD8A are single-gene markers nested within the aggregate T-cell signature.

| Predictor | $\beta$ (95% CI) | p value |
| --- | --- | --- |
| All T-cell module | 0.74 (0.64-0.85) | <0.001 |
| Macrophage module | -0.10 (-0.21-0.02) | 0.10 |
| Endothelial | 0.05 (-0.03-0.13) | 0.20 |
| Myofibroblast | 0.25 (0.14-0.37) | <0.001 |
| Age | -0.01 (-0.01--0.00) | 0.06 |
| Male sex | 0.13 (-0.00-0.26) | 0.05 |
| Sarcopenia | -0.11 (-0.26-0.04) | 0.15 |

**Table S15A. Determinants of muscle CXCL9 expression in the whole cohort.** The model included 340 participants ($R^2 = 0.49$; adjusted $R^2 = 0.48$).

| Predictor | $\beta$ (95% CI) | p value |
| --- | --- | --- |
| All T-cell module | 0.78 (0.61-0.96) | <0.001 |
| Macrophage module | -0.12 (-0.38-0.14) | 0.36 |
| Endothelial | -0.12 (-0.31-0.07) | 0.22 |
| Myofibroblast | 0.34 (0.11-0.58) | 0.005 |
| Age | -0.01 (-0.03-0.01) | 0.18 |
| Male sex | 0.13 (-0.14-0.41) | 0.34 |

**Table S15B. Determinants of muscle CXCL9 expression in the sarcopenic subgroup.**

The model included 102 participants ($R^2 = 0.61$; adjusted $R^2 = 0.58$).

| Model | Predictor | Standardized<br>$\beta$ (95% CI) | Partial<br>r | Partial<br>$R^2$ | p value | BH q<br>value |
| --- | --- | --- | --- | --- | --- | --- |
| Broad-lineage<br>model | T-cell<br>lineage | 0.68 (0.58-0.77) | 0.62 | 0.39 | <0.001 | <0.001 |
|  | Macrophage<br>lineage | -0.12 (-0.23- -<br>0.01) | -0.12 | 0.01 | 0.03 | 0.03 |
| Exploratory T-cell subset model | CD4 expression | 0.12 (-0.01-0.25) | 0.10 | 0.01 | 0.07 | 0.10 |
|  | CD8A expression | 0.50 (0.41-0.59) | 0.50 | 0.25 | <0.001 | <0.001 |
|  | Macrophage lineage | 0.01 (-0.13-0.14) | 0.01 | 0.00 | 0.90 | 0.90 |
| CD4-only sensitivity model | CD4 expression | 0.24 (0.09-0.39) | 0.17 | 0.03 | 0.002 | 0.002 |
| CD8A-only sensitivity model | CD8A expression | 0.51 (0.42-0.60) | 0.52 | 0.27 | <0.001 | <0.001 |

**Table S16A. Predictor-level associations with standardised muscle CXCL9 expression.**

All analyses included 340 complete cases. The exploratory subset model replaced the aggregate T-cell module with CD4 and CD8A expression.

| Model | N | R<sup>2</sup> | Adjusted R<sup>2</sup> | AIC | BIC |
| --- | --- | --- | --- | --- | --- |
| Base model | 340 | 0.13 | 0.12 | 929.36 | 952.33 |
| Broad-lineage model | 340 | 0.50 | 0.49 | 741.43 | 772.06 |
| Exploratory T-cell subset model | 340 | 0.41 | 0.40 | 802.89 | 837.35 |
| CD4-only sensitivity model | 340 | 0.21 | 0.20 | 899.46 | 930.09 |
| CD8A-only sensitivity model | 340 | 0.40 | 0.39 | 804.34 | 834.97 |

**Table S16B. Overall fit of models explaining standardised muscle CXCL9 expression.**

$\Delta R^2$ is relative to the base model comprising endothelial and myofibroblast modules, age, and sex.

| Terms added | Reduced model | Full model | F | p value |
| --- | --- | --- | --- | --- |
| Addition of CD4 and CD8A expression | Base + macrophage | Base + macrophage + CD4 + CD8A | 62.35 | <0.001 |
| Addition of macrophage lineage | Base + CD4 + CD8A | Base + CD4 + CD8A + macrophage | 0.02 | 0.90 |

**Table S16C. Nested tests of incremental T-cell-subset and macrophage contributions to standardised muscle CXCL9 expression.**

| Pathway interaction | N | Interaction $\beta$ (95% CI) | p value | BH q value |
| --- | --- | --- | --- | --- |
| CXCL9 × sarcopenia → T-cell | 340 | 0.23 (0.08 to 0.39) | 0.003 | 0.02 |
| CXCL9 × sarcopenia → fibrosis | 340 | 0.02 (-0.18 to 0.23) | 0.82 | 0.82 |
| CXCL9 × sarcopenia → fibrosis, adjusted | 340 | 0.07 (-0.12 to 0.25) | 0.48 | 0.73 |
| T-cell × sarcopenia → macrophage | 340 | -0.05 (-0.23 to 0.13) | 0.58 | 0.73 |
| Macrophage × sarcopenia → fibrosis | 340 | 0.12 (-0.07 to 0.31) | 0.23 | 0.57 |

**Table S16D. Sarcopenia interaction tests for key CXCL9–T-cell–macrophage–fibrosis relationships.** Interaction terms tested whether the focal association differed by sarcopenia status. Models used the covariates of the corresponding whole-cohort pathway analyses; BH correction was applied across the five interaction tests shown.

| Analysis | Non-sarcopenic slope (95% CI) | Sarcopenia slope (95% CI) |
| --- | --- | --- |
| CXCL9 → T-cell | 0.56 (0.45 to 0.67) | 0.80 (0.68 to 0.91) |
| CXCL9 → fibrosis | 0.24 (0.10 to 0.38) | 0.27 (0.11 to 0.42) |
| CXCL9 → fibrosis, adjusted | 0.11 (-0.03 to 0.26) | 0.18 (0.01 to 0.35) |
| T-cell → macrophage | 0.50 (0.35 to 0.65) | 0.45 (0.29 to 0.60) |
| Macrophage → fibrosis | 0.49 (0.37 to 0.61) | 0.61 (0.44 to 0.79) |

**Table S16E. Sarcopenia-stratified slopes for key pathway associations.** Values are standardised slopes (95% CI) estimated from the corresponding interaction models. Formal between-group inference is provided by the interaction terms in Table S16D.

| Model | Predictor | $\beta$ (95% CI) | p value |
| --- | --- | --- | --- |
| Full stromal-adjusted fibrosis<br>model | CXCL9 | -0.06 (-0.16-0.04) | 0.24 |
|  | All T-cell module | 0.05 (-0.06-0.17) | 0.34 |
|  | Macrophage module | 0.13 (0.03-0.24) | 0.01 |
|  | Endothelial module | 0.10 (0.02-0.17) | 0.010 |
|  | Myofibroblast module | 0.65 (0.56-0.74) | <0.001 |
|  | Age | -0.01 (-0.09-0.07) | 0.78 |
|  | Male sex | 0.03 (-0.12-0.18) | 0.69 |
|  | Sarcopenia status | -0.00 (-0.17-0.17) | 0.99 |
| Stromal-adjusted<br>immune residual<br>model | CXCL9 | -0.08 (-0.18-0.01) | 0.10 |
|  | All T-cell module | 0.07 (-0.03-0.18) | 0.16 |
|  | Macrophage module | 0.09 (0.01-0.18) | 0.03 |

**Table S17. Associations of immune and stromal modules with the muscle fibrosis score.** The full model included CXCL9, T-cell, macrophage, endothelial, and myofibroblast modules, age, sex, and sarcopenia status. The stromal-adjusted immune-residual model evaluated CXCL9, T-cell, and macrophage contributions after accounting for stromal signatures. All analyses included 340 complete cases.

| Model | Predictor | Standardized<br>$\beta$ (95% CI) | Partial r | Partial<br>$R^2$ | p value | BH q<br>value |
| --- | --- | --- | --- | --- | --- | --- |
| Broad-lineage<br>model | CXCL9<br>expression | 0.14 (0.02 to<br>0.26) | 0.12 | 0.02 | 0.02 | 0.02 |
|  | T-cell<br>lineage | -0.18 (-0.32 to<br>-0.05) | -0.15 | 0.02 | 0.01 | 0.01 |
|  | Macrophage<br>lineage | 0.53 (0.42 to<br>0.64) | 0.47 | 0.22 | 0.00 | 0.00 |
| Exploratory<br>macrophage-subset model | CXCL9<br>expression | 0.15 (0.02 to<br>0.27) | 0.13 | 0.02 | 0.02 | 0.03 |
|  | T-cell<br>lineage | -0.13 (-0.26 to<br>0.00) | -0.11 | 0.01 | 0.05 | 0.06 |
|  | Reparative<br>macrophage<br>module | 0.11 (-0.04 to<br>0.25) | 0.08 | 0.01 | 0.15 | 0.15 |
|  | Resident<br>macrophage<br>module | 0.35 (0.20 to<br>0.49) | 0.25 | 0.06 | 0.00 | <0.001 |
|  | Recruited<br>macrophage<br>module | 0.22 (0.06 to<br>0.37) | 0.15 | 0.02 | 0.01 | 0.01 |
|  | Inflammatory<br>macrophage<br>module | -0.25 (-0.40 to<br>-0.10) | -0.18 | 0.03 | 0.00 | 0.00 |

**Table S18A. Predictor-level associations with standardised muscle fibrosis expression.**

All analyses included 340 complete cases. The exploratory subset model replaced the aggregate macrophage-cell module with reparative, resident, recruited and inflammatory macrophage expression.

| **Model** | **N** | **R<sup>2</sup>** | **Adjusted R<sup>2</sup>** | **AIC** | **BIC** |
| --- | --- | --- | --- | --- | --- |
| Base model | 340 | 0.12 | 0.11 | 935.95 | 966.58 |
| Broad-lineage model | 340 | 0.12 | 0.11 | 935.95 | 966.58 |
| Exploratory macrophage-subset model | 340 | 0.32 | 0.30 | 853.15 | 887.61 |
| Reparative macrophage-only sensitivity model | 340 | 0.32 | 0.30 | 854.88 | 900.83 |
| Resident<br>macrophage -<br>only sensitivity<br>model | 340 | 0.24 | 0.22 | 888.46 | 922.92 |
| Recruited<br>macrophage-only sensitivity<br>model | 340 | 0.30 | 0.28 | 861.85 | 896.31 |
| Inflammatory<br>macrophage-only sensitivity<br>model | 340 | 0.13 | 0.11 | 935.73 | 970.19 |

**Table S18B. Overall fit of models explaining standardised muscle fibrosis expression.**

$\Delta R^2$ is relative to the base model comprising endothelial and myofibroblast modules, age, and sex.

| Terms added | Reduced<br>model | Full model | F | p value |
| --- | --- | --- | --- | --- |
| Addition of aggregate<br>macrophage lineage | Base | Base +<br>macrophage<br>lineage | 94.04 | <0.001 |
| Addition of reparative,<br>inflammatory,<br>resident and recruited<br>macrophage subsets | Base | Base +<br>macrophage-subset modules | 24.63 | <0.001 |

**Table S18C. Nested tests of incremental macrophage subset contributions to standardised muscle fibrosis expression.**

| Path | Estimate (95% CI) | p value |
| --- | --- | --- |
| CXCL9 → All T cells | 0.68 (0.47-0.83) | <0.001 |
| Macrophage → Fibrosis | 0.13 (0.03-0.25) | 0.02 |
| All T cells → Macrophage | 0.51 (0.41-0.62) | <0.001 |
| Serial indirect effect:<br>CXCL9 → T cells →<br>Macrophage → Fibrosis | 0.05 (0.01-0.09) | 0.02 |
| Indirect via T cells only | 0.04 (-0.04 to 0.12) | 0.39 |
| Indirect via Macrophages only | -0.02 (-0.04 to 0) | 0.09 |
| Direct effect | -0.06 (-0.18 to 0.06) | 0.30 |
| Total indirect effect | 0.07 (-0.01-0.15) | 0.10 |
| Total effect | 0.01 (-0.08-0.08) | 0.87 |

**Table S19. Cross-sectional sequential mediation model linking CXCL9 to fibrosis through T-cell and macrophage modules.** Estimates and 95% confidence intervals were obtained by bootstrap (5000 samples). Models adjusted for age, sex, and sarcopenia status; n = 340. The model assesses pathway consistency.

| Mediator | Indirect effect<br>(95% CI) | p value | Direct effect<br>(95% CI) | p value |
| --- | --- | --- | --- | --- |
| *Whole cohort* |  |  |  |  |
| Macrophage | 0.01 (0.01-0.02) | <0.001 | 0.01 (0.00-0.02) | 0.03 |
| T-cell | 0.01 (0.01-0.02) | <0.001 | 0.01 (-0.01-0.02) | 0.28 |
| *Sarcopenia only* |  |  |  |  |
| Macrophage | 0.02 (0.00-0.03) | 0.02 | 0.00 (-0.01-0.02) | 0.72 |
| T-cell | 0.01 (0.00-0.03) | 0.04 | 0.01 (-0.01-0.03) | 0.37 |

**Table S20. Single-mediator models of the CXCL9-fibrosis association.** Indirect and direct effects were estimated with bootstrap models (5000 samples) adjusted for age, sex, and sarcopenia status in the whole cohort (n = 340 and age and sex in the sarcopenic subgroup (n = 102).

| Path or effect | Estimate (95% CI) | p value |
| --- | --- | --- |
| CXCL9 → CD8 | 0.58 (0.36-0.73) | <0.001 |
| CD8 → Resident macrophage | 0.20 (0.08-0.33) | 0.002 |
| Resident macrophage → Fibrosis | 0.20 (0.11-0.30) | <0.001 |
| Sequential indirect:<br>CXCL9 → CD8 → Resident<br>macrophage → Fibrosis | 0.02 (0.01-0.04) | 0.01 |
| Indirect via CD8 only | -0.02 (-0.07-0.04) | 0.52 |
| Indirect via resident macrophage<br>only | 0.01 (-0.01-0.05) | 0.29 |
| Direct effect | -0.02 (-0.11 to 0.08) | 0.75 |
| Total indirect effect | 0.02 (-0.04-0.08) | 0.48 |
| Total effect | 0.01 (-0.08-0.08) | 0.88 |

**Table S21. Exploratory sequential mediation through CD8A and resident-macrophage signatures.** Estimates and 95% confidence intervals were obtained by bootstrap (5000 samples) and adjusted for age, sex, and sarcopenia status. CD8A is a single-gene surrogate. Analysis is hypothesis-generating.

|  |  |
| --- | --- |
|  | CXCL9 (pg/ml) |

| Biological Replicate | Myotube | Fibroblast | Cell-normalised fibroblast | Expected additive | Observed co-culture | Excess over additive (%) | Synergy Index |
| --- | --- | --- | --- | --- | --- | --- | --- |
| 1 | 136.61 | 31.65 | 22.61 | 159.21 | 291.32 | 82.97 | 1.83 |
| 2 | 659.04 | 163.54 | 116.81 | 775.85 | 2251.88 | 190.25 | 2.90 |

**Table S22. CXCL9 secretion from fibroblast monoculture, myotube monoculture and myotube:fibroblast co-culture.** Co-culture used the same number of myotubes as per the mono-culture, 15000 per well. Fibroblasts were 7000 cells in the monoculture and 5000 in the co-culture, and scaling was used.

| Cell Population | Biological replicates (n) | Group 1 (mean of %) | Group 2 (mean of %) | Mean % difference | p-value |
| --- | --- | --- | --- | --- | --- |
|  |  | *IFNγ negative* | *IFNγ positive* |  |  |
| CD8 TEMRA | 3 | 9.44 | 27.81 | 18.37 | **0.046** |
| CXCR3+ CD8 TEMRA | 3 | 5.17 | 15.74 | 10.57 | 0.059 |
| PD1+ CD8 TEMRA | 3 | 1.31 | 4.52 | 3.21 | 0.057 |
| CD56+ CD8 TEMRA | 3 | 3.84 | 11.57 | 7.73 | **0.049** |
|  |  | *Isotype* | *Anti-CXCL9* |  |  |
| CD8 TEMRA | 3 | 27.17 | 19.20 | -7.97 | 0.087 |
| CXCR3+ CD8 TEMRA | 3 | 17.92 | 10.40 | -7.52 | **0.035** |

|  |  |  |  |  |  |  |
| --- | --- | --- | --- | --- | --- | --- |
| PD1+<br>TEMRA | CD8 | 3 | 4.97 | 2.61 | -2.36 | **0.009** |
| CD56+<br>TEMRA | CD8 | 3 | 10.96 | 6.79 | -4.16 | 0.100 |
|  |  |  | **DMSO control** | **AMG487** |  |  |
| CD8 TEMRA |  | 3 | 16.80 | 11.03 | -5.77 | **0.019** |
| CXCR3+<br>TEMRA | CD8 | 3 | 10.59 | 5.99 | -4.60 | **0.031** |
| PD1+<br>TEMRA | CD8 | 3 | 3.97 | 1.46 | -2.51 | 0.093 |
| CD56+<br>TEMRA | CD8 | 3 | 5.69 | 3.81 | -1.87 | **0.016** |

**Table S23. CD8 TEMRA subset transwell migration** toward IFNγ-negative or IFNγ-conditioned myotube–fibroblast supernatant, with isotype/DMSO controls, CXCL9 neutralisation (anti-CXCL9) or CXCR3 antagonism (AMG487) for CD8 TEMRA, CXCR3+ CD8 TEMRA, PD1+ CD8 TEMRA and CD56+ CD8 TEMRA populations, shown as volume-normalised counts. Blockade significantly reduced migration of PD1+ CD8 TEMRA with anti-CXCL9 ($p = 0.018$) and of CD8 TEMRA ($p = 0.038$) and CD56+ CD8 TEMRA ($p = 0.032$) with AMG487; IFNγ priming increased CD8 TEMRA migration at trend level ($p = 0.09$). Data from $n = 3$ independent PBMC donors and $n = 2$ myotube–fibroblast donor pairs, each with 2 technical replicates. Dots show per-donor means; bars show across-donor mean ± SEM. Statistics are one-sided paired t-tests on per-donor means.

| Macrophage Phenotype | Biological replicates (n) | M0 (% of all macrophages) | CD8-educated macrophages (% of all macrophages) | Mean difference | pvalue |
| --- | --- | --- | --- | --- | --- |
| Reparative | 3 | 37.7 | 5.69 | -32.0 | 0.36 |
| Mixed | 3 | 37.1 | 71.3 | 34.2 | 0.27 |
| Inflammatory | 3 | 11.8 | 17.7 | 5.88 | 0.25 |

**Table S24. Comparison of macrophage polarisation states** between naive (M0) and CD8-educated macrophages. Frequencies of reparative (CD163+CD86-), mixed (CD163+CD86+) and inflammatory (CD86+CD163-) phenotypes, as a percentage of HLA-DR+CD11b+ macrophages, were compared after 48h of CD8 education across n = 3 biological replicates. Values are mean percentages; mean difference is CD8-educated minus M0. p values are from paired t-tests on per-replicate frequencies. CD8 education shifted macrophages away from the reparative state (-32.0%) toward the mixed (+34.2%) and, to a lesser extent, inflammatory (+5.9%) states; none of the individual shifts reached statistical significance (p = 0.36, 0.27 and 0.25, respectively), consistent with a reproducible but non-canonical phenotype across donors rather than a discrete polarisation change. Gating is shown in Methods Figure M4.

| Condition | Biological replicate | Number of technical duplicates | Cell Count | TGF-β1, pg/ml | Cell-normalised TGF-β1 Concentration (per 100,000 cells) |
| --- | --- | --- | --- | --- | --- |
| *24h* |  |  |  |  |  |
| CD8 | 1 | 2 | 100000 | 619.5 ± 108.9 | 619.5 ± 108.9 |
|  | 2 | 2 | 100000 | 1549.62 ± 12.04 | 1549.62 ± 12.04 |
| M0 | 1 | 2 | 80000 | 673.5 ± 190.5 | 841.87 ± 238.12 |
|  | 2 | 2 | 54950 | 1678.44 ± 14.91 | 3054.49 ± 27.13 |
| CD8 educated macrophages | 1 | 4 | 81500 | 690.29 ± 82.38 | 846.99 ± 101.08 |
|  | 2 | 2 | 57550 | 1883.47 ± 30.15 | 3272.75 ± 52.38 |
| *96h* |  |  |  |  |  |
| CD8 | 1 | 2 | 100000 | 691.8 ± 6.6 | 691.8 ± 6.6 |
|  | 2 | 2 | 100000 | 2673.09 ± 133.65 | 2673.09 ± 133.65 |
| M0 | 1 | 2 | 80000 | 842.1 ± 9.3 | 1052.62 ± 11.63 |
|  | 2 | 2 | 54950 | 2376.75 ± 105.85 | 4325.3 ± 192.63 |
| CD8 educated macrophages | 1 | 2 | 76000 | 991.5 ± 104.1 | 1304.61 ± 136.97 |
|  | 2 | 2 | 57550 | 2635.49 ± 102.43 | 4579.48 ± 177.99 |

**Table S25A. Secreted total TGF-β1 concentrations in the immune–stromal co-culture.**

IFNγ-conditioned myotube–fibroblast stroma was co-cultured for 24 h and 96 h with the indicated immune populations: magnetically sorted CD8 T cells (CD8), naive monocyte-derived macrophages (M0), CD8-educated macrophages, and CD8-educated macrophages generated under anti-IFNγ or isotype-control antibody during the 48h education step. Total TGF-β1 was measured by immunoassay; values are mean±SD across the technical replicates indicated for each condition, given both as raw pg/mL and normalised to 100,000 cells using the corresponding cell counts. Data are from n = 2 independent PBMC donors paired with n = 2 separately expanded myotube–fibroblast donor pairs. The two replicates differed in protein secretion and values are therefore reported separately by replicate. All comparisons are descriptive.

| Condition | Biological replicate | Number of | Cell Count | PAI-1, pg/ml | Cell-normalised PAI-1 |
| --- | --- | --- | --- | --- | --- |

|  |  | technical replicates |  |  | Concentration (per 100,000 cells) |
| --- | --- | --- | --- | --- | --- |
| *24h* |  |  |  |  |  |
| CD8 | 1 | 2 | 100000 | 1836.81 | 1836.81 |
|  | 2 | 2 | 100000 | 274574.61 ± 3959.85 | 274574.61 ± 3959.85 |
| M0 | 1 | 1 | 80000 | 1587.79 | 1984.73 |
|  | 2 | 2 | 54950 | 257473.04 ± 11954.72 | 468558.76 ± 21755.64 |
| CD8 educated macrophages | 1 | 1 | 81500 | 3598.38 | 4415.2 |
|  | 2 | 3 | 57550 | 198718.38 ± 8863.85 | 345296.92 ± 15402 |
| *96h* |  |  |  |  |  |
| CD8 | 1 | 2 | 100000 | 981.22 ± 167.29 | 981.22 ± 167.29 |
|  | 2 | 2 | 100000 | 456878.34 ± 33150.53 | 456878.34 ± 33150.53 |
| M0 | 1 | 2 | 80000 | 910.41 ± 236.66 | 1138.01 ± 295.83 |
|  | 2 | 2 | 54950 | 246756.48 ± 202394.87 | 449056.38 ± 368325.51 |
| CD8 educated macrophages | 1 | 2 | 76000 | 18795.14 ± 6365.22 | 24730.45 ± 8375.29 |
|  | 2 | 3 | 57550 | 366886.42 ± 162259.49 | 637508.98 ± 281945.24 |

**Table S25B. Secreted PAI-1 concentrations in the immune–stromal co-culture.**

| Condi<br>on | Cell<br>s | Tec<br>hnic<br>al<br>repli<br>cate<br>s (n) | CXC<br>L8<br>(pg/<br>ml) | CXCL<br>8,<br>norm<br>alised | IL-6<br>(pg/<br>ml) | IL-6,<br>norm<br>alise<br>d | CXC<br>L9<br>(pg/<br>ml) | CXCL<br>9,<br>norma<br>lised t | IFNγ<br>(pg/<br>ml) | IFNγ,<br>norm<br>alise<br>d |
| --- | --- | --- | --- | --- | --- | --- | --- | --- | --- | --- |
| 24h |  |  |  |  |  |  |  |  |  |  |
| CD8 | 1000<br>00 | 2 | 77.5<br>5 ±<br>3.95 | 77.55<br>± 3.95 | 9004.<br>05 ±<br>457.8<br>5 | 9004.<br>05 ±<br>457.8<br>5 | 2962.<br>9 ±<br>120.6 | 2962.9<br>±<br>120.6 | 3640<br>.55 ±<br>215.<br>25 | 3640.<br>55 ±<br>215.2<br>5 |
| M0 | 5495<br>0 | 2 | 30.0<br>5 ±<br>11.0<br>5 | 54.69<br>±<br>20.11 | 7295.<br>35 ±<br>1120.<br>95 | 13276<br>.34 ±<br>2039.<br>95 | 3937.<br>95 ±<br>1102.<br>85 | 7166.4<br>2 ±<br>2007.0<br>1 | 2471<br>.3 ±<br>519.<br>6 | 4497.<br>36 ±<br>945.5<br>9 |
| CD8<br>educate<br>d<br>macroph<br>ages | 5755<br>0 | 3 | 816.<br>7 ±<br>183.<br>49 | 1419.<br>11 ±<br>318.8<br>3 | 1457<br>7.4 ±<br>1956.<br>77 | 25329<br>.97 ±<br>3400.<br>12 | 6453.<br>67 ±<br>2040.<br>72 | 11214.<br>02 ±<br>3545.9<br>9 | 2980<br>.67 ±<br>30.8<br>5 | 5179.<br>26 ±<br>53.6 |
| 96h |  |  |  |  |  |  |  |  |  |  |
| CD8 | 1000<br>00 | 2 | 165.<br>95 ±<br>10.2<br>5 | 165.9<br>5 ±<br>10.25 | 1418<br>0.15<br>±<br>76.15 | 14180<br>.15 ±<br>76.15 | 4697.<br>4 ±<br>411.7 | 4697.4<br>±<br>411.7 | 1067<br>.55 ±<br>43.4<br>5 | 1067.<br>55 ±<br>43.45 |
| M0 | 5495<br>0 | 2 | 79.1<br>5 ±<br>4.95 | 144.0<br>4 ±<br>9.01 | 1277<br>6.25<br>±<br>327.1<br>5 | 23250<br>.68 ±<br>595.3<br>6 | 4441.<br>65 ±<br>525.1<br>5 | 8083.0<br>8 ±<br>955.69 | 2036<br>.25 ±<br>95.9<br>5 | 3705.<br>64 ±<br>174.6<br>1 |
| CD8<br>educate<br>d<br>macroph<br>ages | 5755<br>0 | 2 | 1554<br>.95 ±<br>41.8<br>5 | 2701.<br>91 ±<br>72.72 | 2663<br>5.65<br>±<br>565.3<br>5 | 46282<br>.62 ±<br>982.3<br>6 | 3218<br>2.15<br>±<br>359.9<br>5 | 55920.<br>33 ±<br>625.46 | 2938<br>.8 ±<br>92 | 5106.<br>52 ±<br>159.8<br>6 |

**Table S26. Inflammatory and chemokine secretome of the immune–stromal co-culture.**

CXCL8, IL-6, CXCL9 and IFNγ concentrations after 24h and 96h of co-culture of IFNγ-conditioned myotube-fibroblast stroma with CD8 T cells, naive macrophages (M0), CD8-educated macrophages, or CD8-educated macrophages generated under anti-IFNγ or isotype control. Values are mean±SD across technical replicates (n indicated per condition) from a single biological donor, given as raw pg/mL and normalised to 100,000 cells. CXCL9 was strictly IFNγ-dependent and markedly enriched in the CD8-educated condition at 96h, whereas CXCL8 rose about 40-fold over M0 in the CD8-educated condition. IFNγ is a QC condition and reflects residual conditioning stimulus. As a single-donor dataset, all values are descriptive.
